# 45-Color Full Spectrum Flow Cytometry Panel for Deep Immunophenotyping of the Major Lineages Present in Human Peripheral Blood Mononuclear Cells with Emphasis on the T cell Memory Compartment

**DOI:** 10.1101/2024.04.27.591472

**Authors:** Lily M. Park, Joanne Lannigan, Quentin Low, Maria C. Jaimes, Diana L. Bonilla

**Affiliations:** Cytek Biosciences, Inc., Scientific Commercialization, 47215 Lakeview Boulevard, Fremont, CA 94538; Flow Cytometry Support Services, LLC, Alexandria, VA 22314

**Keywords:** Full spectrum flow cytometry, Aurora, blood immunophenotyping, high-dimensional flow cytometry, OMIP-069, Memory T cells, T cell exhaustion, T cell activation

## Abstract

The need for more in-depth exploration of the human immune system has moved the flow cytometry field forward with advances in instrumentation, reagent development and user-friendly implementations of data analysis methods. The increase in the number of markers evaluated simultaneously requires a careful selection of highly overlapping dyes to avoid introducing detrimental spread and compromising population resolution. In this manuscript, we present the strategy used in the development of a high-quality human 45-color panel which allows for comprehensive characterization of major cell lineages present in circulation including T cells, gamma delta T cells, NKT-like cells, B cells, NK cells, monocytes, basophils, dendritic cells, and ILCs, as well as more in-depth characterization of memory T cells. The steps taken to ensure that each marker in the panel was optimally resolved are discussed in detail. We highlight the outstanding discernment of cell activation, exhaustion, memory, and differentiation states of CD4+ and CD8+ T cells using this 45-color panel, enabling an in-depth description of very distinct phenotypes associated with the complexity of the T cell memory response. Furthermore, we present how this panel can be effectively used for cell sorting on instruments with a similar optical layout to achieve the same level of resolution. Functional evaluation of sorted specific rare cell subsets demonstrated significantly different patterns of immunological responses to stimulation, supporting functional and phenotypic differences within the T cell memory subsets. In summary, the combination of flow cytometry full spectrum technology, careful assay design and optimization, results in high resolution multiparametric assays. This approach offers the opportunity to fully characterize immunological profiles present in peripheral blood in the context of infectious diseases, autoimmunity, neurodegeneration, immunotherapy, and biomarker discovery.

**PURPOSE AND APPROPRIATE SAMPLE TYPES:** This 45-color flow cytometry-based panel was developed as an expansion of the previously published OMIP-069 [1] and serves as an in-depth immunophenotyping of the major cell subsets present in human peripheral blood. The goal of this panel is to maximize the amount of high-quality data that can be acquired from a single sample, not only for more in-depth characterization of the immune system, but also to address the issue of limited sample availability. The panel’s development included identifying fluorochromes that could improve the performance of the original 40-color panel and expanding the number of markers for deeper delineation of memory status of T cell subpopulations. To increase the number of markers, it was critical that any expansion did not negatively impact the resolution and quality of the data. To achieve this, the fluorochrome combinations were carefully characterized to ensure optimal resolution of each marker. The panel allows for deep characterization of the major cell lineages present in circulation (CD4 T cells, CDS T cells, regulatory T cells, yo T cells, NKT-like cells, B cells, NK (Natural Killer) cells, monocytes, and dendritic cells), while also providing an in-depth characterization of the T cell compartment, with a combination of activation, inhibitory, exhaustion, and differentiation markers. The panel supports deep exploration of the memory status of CD4^+^ T cells, CDS^+^ T cells, and NKT-like cells. The steps taken in the optimization of the panel ensured outstanding resolution of each marker within the multicolor panel and unequivocal identification of each cell subset. This panel design and optimization will enhance the ability to characterize immunological profiles present in peripheral blood in the context of oncology, infectious diseases, autoimmunity, neurodegeneration, immunotherapy, and biomarker discovery.

The panel was developed using fresh and cryopreserved human peripheral blood mononuclear cells (PBMCs) from healthy adults. We have not tested the panel on whole blood or biopsies; hence it is anticipated that the panel might require further optimization to be used with other sample types.

## BACKGROUND

Characterization of immune cell populations has played a critical role in understanding disease pathogenesis, progression, and response to treatment [2]. Full spectrum flow cytometry (FSFC) has enabled a deeper characterization of the complexity of immune responses through the development of highly multiparametric panels for broad immunophenotyping [3–7]. OMIP-069 [1] was the first optimized panel demonstrating that 40 different fluorochromes can be effectively used in combination, without compromising the resolution of each individual marker. This hallmark was achieved by using a full spectrum cytometer with optimized setup, well characterized optical performance, extensive knowledge of panel design and assay optimization [S], and advances in fluorochrome development [1]. Since its publication, the panel has been adopted in many laboratories worldwide [9,10].

After the publication of OMIP-069, new fluorochromes were developed, opening the opportunity to improve and expand panel performance. Using a rigorous strategy, new fluorochromes were evaluated to identify expanded combinations that could perform optimally (Online Figures 1-6, and 7-14, Online Table 2). The parameters taken into consideration to evaluate performance included the resolution of each individual maker when stained alone and in the full panel (Online Figures 2-5), unequivocal gating, identification of all expected cell subsets, and robust performance across multiple donors and instruments. The result is a 45-color panel (Online Figure 6), optimized in a Cytek Aurora cytometer equipped with 5 lasers and 64 detectors (Online Table 1) that allows for assessment of the frequency of B cells, NK cells, monocytes, basophils, CD4 and CDS T cells, regulatory T cells (Tregs), yo T cells, NKT-like cells, innate lymphoid cells (ILCs), dendritic cells (DCs), and their main subpopulations in human peripheral blood samples, using the majority of markers (Online Table 3) previously described in OMIP-069 [1]. We show supportive evidence of reagent titration (Online Figure 7), reference control selection (Online Figure S, Online Table 4), unmixing assessment (Online Figure 9), staining protocol optimization (Online Figure 10), and marker resolution evaluation (Online Figure 11-13).

The panel was designed for full evaluation of main lineages in blood (Figure 1 and Online Figure 14), and deep characterization of the T cell compartment, by including memory and effector differentiation markers (CD27, CD2S, CD45RA, CD127, CD95, GPR56, and DNAM-1), activation markers (CD3S, CD39, and HLA-DR), homing markers (CCR5, CCR6, CCR7, CXCR3, CXCR5, and CD11b), inhibitory receptors (PD-1, TIM-3, KLRG1, CD161, and TIGIT), and markers of tissue residency (CD103). The differentiation of T cells into memory or effector states can be highly heterogeneous, requiring the use of additional markers for deeper immunological evaluations [11]. To address this, the panel includes S additional targets (TIM-3, TIGIT, KLRG1, CD161, GPR56, CD103, DNAM-1, and CD11b) for an in-depth characterization of functionality in memory T cell subsets.

**Figure 1A.**
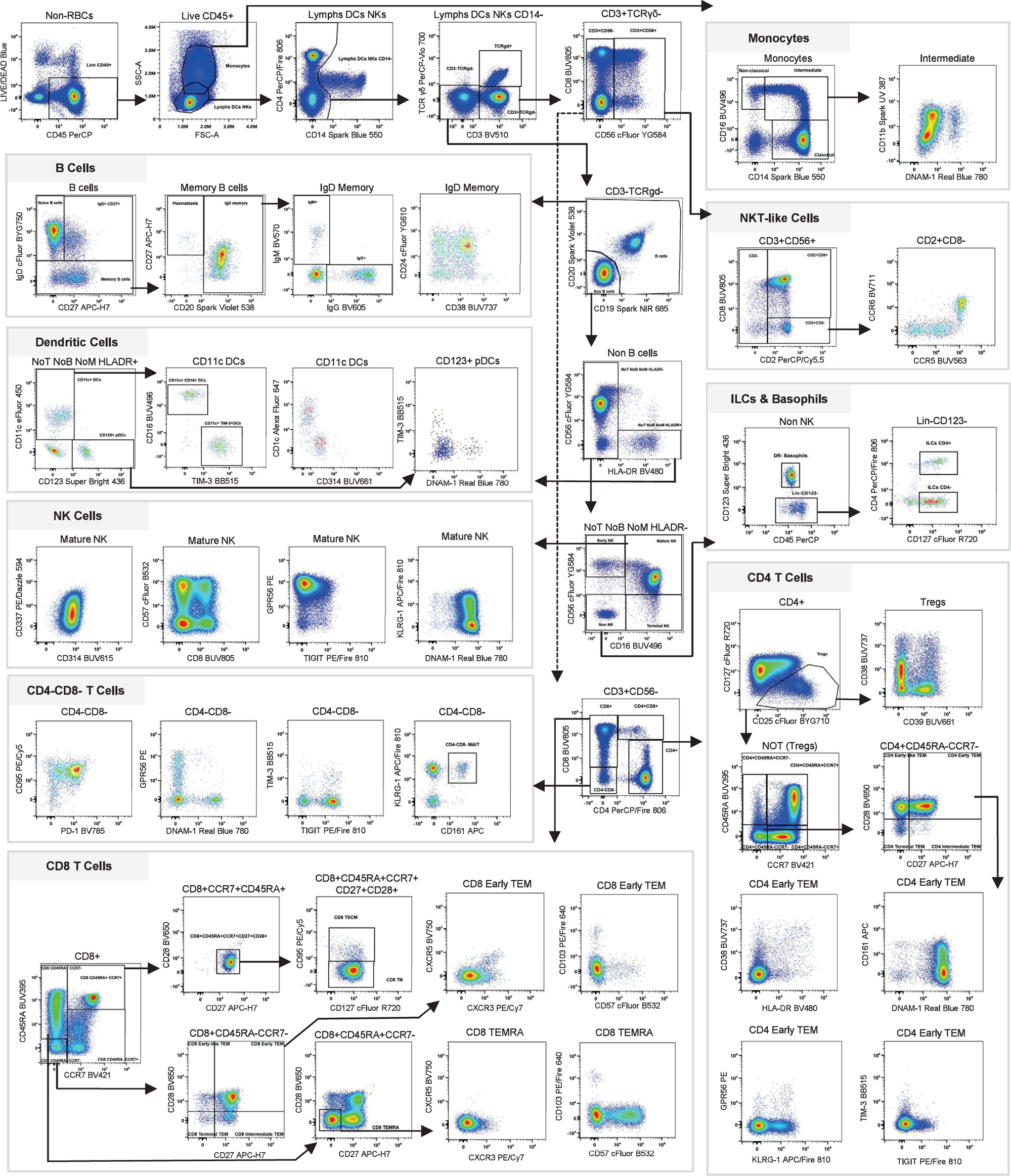
Manual gating strategy. A summary version of the full gating strategy used to identify the main cellular subsets is presented, with all markers in the panel being displayed in bivariate plots. Arrows are used to visualize the relationships across plots. First, cell doublets and red blood cells were excluded (not shown). Next, cells were gated on viability (Live dead blue) and CD45 positivity. Live CD45+ cells were then segregated, based on FSCA/SSC-A properties, into Lymphocytes/DCs/NK cells and monocytes. Monocytes were further classified by CD14 and CD16 expression as non-classical (CD14^−^CD16^+^), intermediate (CD14^+^CD16^+^), and classical (CD14^+^CD16^−^). The expression of DNAM-1 and CD11b is shown for the intermediate population. From the lymphocyte/DC/NK gate, CD14^+^ events were excluded, and the following populations were identified: CD3^−^TCRy6^−^, CD3^+^TCRy6^+^, and CD3^+^TCRy6^−^. The CD3^+^TCRy6^−^ population was divided into CD19+CD20+ and CD19-CD20-. B cells were defined as CD3^−^TCRy6^−^CD19^+^CD20^+^, and gated as IgD^+^CD27^−^, IgD^+^CD27^+^, or IgD^−^; the lgD^−^ subset was divided into plasmablasts or lgD^−^ memory B cells, based on CD20 and CD27 expression. The expressions of lgG, lgM, CD24, and CD38 were assessed within the lgD^−^ memory B cells. CD19^−^ CD20^−^events were classified based on CD56 and HLA-DR expression. Dendritic cells (DCs) were identified by gating on CD14^−^CD3^−^CD19^−^CD20^−^CD56^−^HLA-DR^+^ and from there CD123^+^ (pDCs) and CD11c^+^ DCs were identified. CD11c^+^ DCs were further divided into CD16^+^ or TIM-3^+^. The expression of TlM-3 and DNAM-1 is shown as pDCs, and the expression of CD1c and CD314 is shown as CD11c^+^ DCs. NK cells were defined as CD14^−^CD3^−^TCRy6^−^CD19^−^CD20^−^HLA-DR^−^ and classified as early NK (CD56^+^CD16^−^), mature NK (CD56^+^CD16^+^), and terminal NK (CD56^−^CD16^+^) cells. Expression of CD337, CD314, CD57, CD8, GPR56, TlGlT KLRG1, and DNAM-1 is shown for the mature NK cells. CD3^+^CD56^−^TCRy6^−^ events were classified based on CD8 and CD56 expression. NKT-like cells were defined as CD3^+^TCRy6^−^CD56^+^ and subclassified as CD2^−^, CD2^+^CD8^−^, and CD2^+^CD8^+^. The expression of CCR6 and CCR5 is shown in the CD2^+^CD8^−^ NKT-like subset. Subsequently, basophils and innate lymphoid cells (lLCs) were identified out of CD14^−^CD3^−^TCRy6^−^ CD19^−^CD20^−^HLA-DR^−^CD56^−^CD16^−^ population. Basophils are CD123 positive and lLCs are CD123 negative. lLCs were further identified using CD127^+^ and 2 subsets identified based on CD2 expression. The CD3^+^CD56^−^ events were separated based on the expression of CD4 and/or CD8. From the CD4^+^ population, regulatory T-cells (Tregs) were identified as CD127^−^CD25^+^. The expression of CD38 and CD39 is shown for Tregs. The non-T regs (CD127^+^CD25^−^) were subclassified using CCR7, CD45RA, CD27, CD28, and CD95 for definition of naïve, memory, and effector CD4 T cell subsets. The expressions of CD161, DNAM-1, HLA-DR, CD38, GPR56, KLRG1, TlM-3, and TlGlT are shown for CD4^+^ early T effector memory cells (early TEM). Similarly, CD8 T cells were profiled for memory status. Naïve T cells (TN) were defined as CCR7^+^CD45RA^+^CD27^+^CD28^+^CD95^−^, T memory stem cells (TSCM) as CCR7^+^CD45RA^+^CD27^+^CD28^+^CD95^+^, early TEM as CCR7^+^CD45RA^−^ CD27^+^CD28^+^ CD95^+^, early-like TEM as CCR7^+^CD45RA^−^CD27^−^CD28^+^CD95^+^, intermediate TEM as CCR7^+^CD45RA^−^CD27^+^CD28^−^CD95^+^, terminal TEM as CCR7^+^CD45RA^−^CD27^−^CD28^−^CD95^+^, and effector memory T cells re-expressing CD45RA (TEMRA) as CCR7^−^CD45RA^+^CD27^−^CD28^−^CD95^+^. The expressions of CXCR5, CXCR3, CD103, and CD57 are shown for CD8^+^ early TEM and TEMRA. All data presented are derived from fresh PBMCs of one healthy donor. Data was analyzed using SpectroFlo software and the full gating strategy is shown in the online data.

**Figure 1B.**
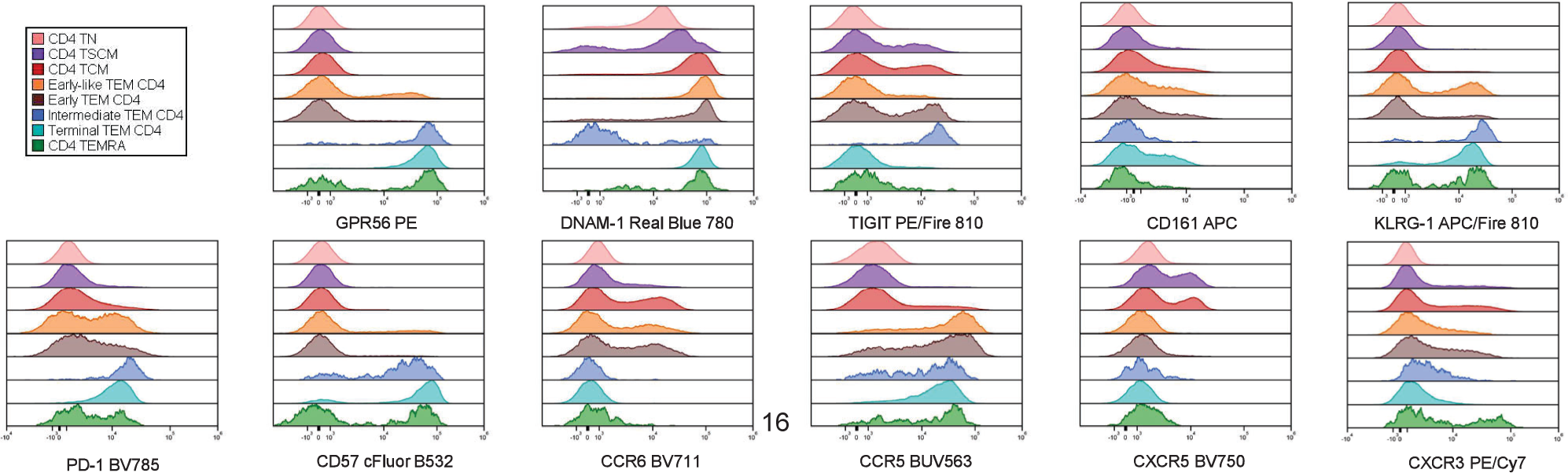
Marker expression in CD4^+^ T cell subsets manually gated. Overlaid histograms for GPR56, DNAM-1, TIGIT, PD-1, CD161, KLRG1, CD57, CCR6, CCR5, CXCR5, HLA-DR, and CXCR3 are shown. Each color represents a gated population, including TN (pink), TSCM (purple), central memory T cells (TCM) (red), early-like TEM (orange), early TEM (brown), intermediate TEM (blue), terminal TEM (cyan), and TEMRA (green). Data were analyzed using FCS Express software.

Naïve T cells (TN) have not been exposed to antigens and have a strong proliferative potential when stimulated. Upon antigen recognition, TN can proliferate, differentiate to effector cells, and generate protective and perdurable responses [12]. Some of the effector populations can turn into long-lived memory cells that provide lasting protection against aggressors, including microorganisms, inflammatory signals, or tumor cells [13]. Memory T cells are identified by the lack of expression of CD45RA and are comprised of a heterogeneous pool of subsets with distinct homing, renewal capacity, effector functions, and marker expression [4]. For example, the CC-chemokine receptor CCR7 is a lymph node homing marker that facilitates distinction of two main memory T cell categories: long-lived central memory (TCM: CCR7^+^CD45RA^−^) and short-lived effector memory (TEM: CCR7^−^CD45RA^−^). TCM have cell renewal capabilities, traffic to secondary lymphoid tissues, and can confer protective immunity against cancer when transferred adoptively [14]. TEM have strong cytolytic capacity, secrete interferon gamma (IFN-y), are responsible for toxicity against target cells, and undergo terminal differentiation after a few cell divisions [11]. CD27 and CD2S are useful to further refine the memory effector classification, identifying early effector (CD45RA^−^ CCR7^−^CD2S^+^CD27^+^), early-like effector (CD45RA^−^CCR7^−^CD2S^+^CD27^−^), intermediate effector (CD45RA^−^CCR7^−^ CD2S^−^CD27^+^), terminal effector (CD45RA^−^CCR7^−^CD2S^−^CD27^−^), and RA-terminal effector (TEMRA: CD45RA^+^CCR7^−^CD2S^−^CD27^−^) [15]. In contrast, TN express both CCR7 and CD45RA. TCM and TEM can circulate in blood and migrate to secondary lymphoid tissues, while resident memory T cells (TRM) are found within the tissues, without recirculation, for immediate functional responses after restimulation at the primary inflammatory site [16]. TRM cells are highly activated and cytotoxic and have been defined by the expression of markers included in this panel e.g., CD103, CD39, PD-1, and TIM-3. CD103 favors the retention of TRM in epithelial tissues and is required for cytotoxic activity [17]. TRM cells have limited proliferative capacity and can become terminally differentiated following reactivation [1S]. With this panel, we were able to successfully identify a small percentage of TRM cells within the CDS^+^ TCM and all the TEM subsets, and in the CD4^+^ early-like and early effector TEM subpopulations.

Immunological memory depends on self-renewing T cells, namely stem cell memory T cells (TSCM). TSCM cells respond efficiently to antigens, have the multipotent ability to derive all memory T cell differentiation clusters, and are critical for efficient T cell cytotoxicity and persistence [19]. TSCM cells express TN markers such as CD45RA, CCR7, CD27, CD2S, and CD127, but in contrast to TN cells, TSCM cells express CD95 [20], and compared to other memory populations, these cells have higher proliferative capacity. With this panel we identify TSCMs in PBMC as CCR7^+^CD45RA^+^CD95^+^. In the samples tested, we found, in agreement with Gattinoni L. et al., that TSCM cells represent less than 5% of the circulating CD4^+^ and CDS^+^ T cells in blood from healthy donors [20]. The identification of these subsets is of immense importance, since there is evidence of the therapeutic potential in targeting both exhausted T cells and TSCM cells for adoptive cancer immunotherapy, by reprograming their memory status to improve anti-tumor control [19,21]. T cell dysfunction is a common finding in cancer. Fraietta J. et al. showed that the presence of early memory T cells and exhausted T cells conditions the effectiveness of CAR-T cell therapy of chronic lymphocytic leukemia [22], while Miller B. et al. showed that exhausted CDS^+^ T cells can mediate anti-tumoral responses after immune check point blockade [23]. In cancer, deeper characterizations of T cell memory subsets are critical for disease characterization, and for development of immunotherapy strategies. The presence of certain markers in tumor-infiltrating lymphocytes (TILs) has been associated with cancer prognosis and clinical response [24–26]. Memory T cell differentiation does not always translate into protective responses and some dysfunctionalities might occur. Under persistent antigenic stimulation, memory T cells can show an exhausted profile (TEX), characterized by limited effector functions such as secretion of IFN-y or Tumor Necrosis Factor-a (TNF-a), cytotoxicity, or proliferation capacity in response to antigen stimulation, as well as higher expression of checkpoint inhibitory receptors [1S,27]. We have included in the panel multiple checkpoint markers associated with cell exhaustion, whose blockade results in T cell and NK cell re-activation [2S-32]. PD-1 is the programmed cell death protein 1, also known as CD279. It is expressed by T cells or B cells, and acts as a checkpoint inhibitor, downregulating T cell responses [33]. TIGIT is the T cell immunoreceptor with Ig and ITIM domain present on T cells and NK cells, binding to CD155 and CD122 on DCs and macrophages. TIGIT is a checkpoint inhibitor that blocks NK and CDS^+^ T cell cytotoxicity [34]. TIM-3 is the T cell immunoglobulin and mucin-domain containing-3 expressed on CD4^+^ T cells, CDS^+^ T cells, T regulatory cell, NK cells, macrophages, and dendritic cells. TIM-3 is also a checkpoint inhibitor upregulated by tumor-infiltrating lymphocytes and associated with T cell exhaustion [29,35]. In addition, the expression of CD57 can be used as an indication of cellular senescence, failure to proliferate, and susceptibility to cell death [36]. In the panel, TEX cells were identified using CD57 and GPR56 in both CD4^+^ and CDS^+^ T cells. GPR56 is the G protein-couple receptor 56, an adhesion molecule expressed in neural cells, muscle cells, hematopoietic precursors, and exhausted terminal effector cytotoxic T and NK cells [16]. Galletti G. et al., characterized the heterogeneity of the human CD95^+^CDS^+^ memory T cell pool in healthy donors by single cell RNA sequencing and flow cytometry [19]. Using part of the gating strategy from this publication, we were able to dissect the heterogeneity of the memory pool and identify TRM, TSCM, TN, TCM, and TEM subpopulations (TEX, early, early-like, intermediate, and terminal). Manual characterization of memory T cell populations is described in detail in Online Figures 15-19.

Other new markers included in the panel are CD11b, KLRG1, CD161, DNAM-1, and CD103. CD11b is integrin aM and is important for cell migration and adhesion in myeloid cells and lymphocytes during inflammation, binding to components of the extracellular matrix and the endothelium. CD11b signaling can negatively regulate T cell activation [1S,37]. KLRG1 is the member 1 of the killer cell lectin-like receptor subfamily G, a glycoprotein expressed by NK cells and on late-differentiated effector and effector memory CD4^+^ and CDS^+^ T cell subsets as a marker of differentiation. KLRG1 is an inhibitory checkpoint receptor that binds to E-cadherin and N-cadherin, and is associated with T cell exhaustion, T cell senescence, and limited proliferative and cytotoxic capacity [16,3S]. KLRG1 has been considered as a biomarker to predict immunotherapy response, being more highly expressed in patients with lung adenocarcinoma that respond to anti-PD-1 therapy [25]. CD161 is the member 1 of the killer cell lectin-like receptor subfamily B expressed by NK cells, CD4^+^ and CDS^+^ T cells, also known as KLRG1 [39]. CD161 binds to Lectin-like transcript-1 (LLT-1) and triggers inhibitory signals and T cell exhaustion [16,26]. DNAM-1 is the DNAX accessory molecule-1 that mediates cell adhesion. DNAM-1, also known as CD226, is a co-stimulatory receptor expressed in NK cells, NKT-like cells, Tregs, B cells, platelets, monocytes, and T cells, and is associated with T cell activation, cytokine secretion, proliferation, migration, and cytotoxicity [40,41]. The presence of these markers in tumor-infiltrating lymphocytes (TILs) has been associated with cancer prognosis and clinical response [24–26]. CD103 is the integrin aE, part of the aEβ7 complex, that binds to e-cadherin and mediates adhesion and homing of T cells within epithelial tissues. CD103 is expressed by intraepithelial lymphocytes (αβ and γδ T cells), lamina propria T cells, regulatory T cells, and a subset of DCs. The presence of CD103^+^ resident memory T cells has been linked to better survival upon treatment in patients with head and neck squamous cell carcinoma [24] and esophageal cancer [42]. For example, Gattas G et al. demonstrated that the presence of CD103^+^ CDS^+^ tumor-infiltrating lymphocytes is associated with better responses to chemoimmunotherapy in patients with head and neck squamous cell carcinoma [43].

Mucosal associated invariant T cells (MAIT) can be found scarcely in peripheral blood, being more represented in tissues, such as liver and intestine, where they play a role in mucosal defense against microbes. MAIT cells are either CD4^−^CDS^−^ or CDS^int^ T cells and have high expression of CD161, CD127, and KLRG1 with low expression of CCR7, CD3S, CD39, GPR56, CD314, CD103, CD57, CXCR3, TIGIT, PD1, TIM-3, and HLA-DR. In some of the donors tested with this 45-color panel, we identified MAIT cells that can express an effector memory phenotype (CD45RA^−^CCR7^−^CD95^+^CD27^+^CD2S^+^CD127^+^), with a chemokine receptor pattern (CCR5^+^CCR6^+^) indicating preferential tissue homing, in agreement with the literature [19,44]. Two subsets were identified by manual gating in the MAIT population, based on the presence or absence of DNAM-1. Other receptors expressed by T cells, already present in OMIP-069 and involved in stimulating or inhibiting responses, were used to support the full characterization of all described subsets. Those receptors include: CD337, the natural cytotoxicity triggering receptor 3, also known as NKp30, with a role in activating cell cytotoxicity [45]; CCR5 (C-C chemokine receptor type 5) and CCR6 (C-C chemokine receptor type 6), involved in chemokine binding, T cell homing, and memory development [46,47]; CD314, the member D of the activating receptor natural killer group 2, important for NK cell activation [4S]; CD39, the [46] ectonucleoside triphosphate diphosphohydrolase-1 with a role in T regulatory suppressive functions [49], chemokine receptors CXCR3 and CXCR5 which regulate T cell trafficking and priming [50,51] the interleukin-7 receptor a (CD127), involved in memory and effector T cell functions [52]; CD3S, the cyclic ADP ribose hydrolase, important for T cell activation and calcium signaling [53]; and then MHC (Major Histocompatibility Complex) class II receptor, human leukocyte antigen-DR isotype (HLA-DR), widely known for its upregulation during T cell activation [54].

Display of the manual gating with focus on the T cell compartment based on the descriptions provided here, is shown in Figure 1A and Online Figure 14. The inclusion of new markers expanded the subclassification of multiple lineage populations in the panel. CD4^+^ T cells, CDS^+^ T cells, CD4^+^CDS^+^ T cells, CD4^−^CDS^−^ T cells, NK cells, and NKT-like cells can be further divided into subsets based on the absence or presence of KLRG1, CD161, GPR56, CD103, DNAM-1, and TIGIT. T cell differentiation and memory status can be clearly sub-compartmentalized based on the expression pattern of GPR56, DNAM-1, TIGIT, PD-1, CD161, KLRG1, CD57, CCR6, CCR5, CXCR5, HLA-DR, and CXCR3 for TN, TSCM, TCM, TEM, and TEMRA CD4^+^ T cells (Figure 1A and Online Figures 15-19). Moreover, the inclusion of additional markers facilitated the characterization of subsets derived from the non-T cell compartment. CD11b and DNAM-1 can be used to subclassify monocytes. We observed a progression from low CD11b expression, in non-classical monocytes, to intermediate levels in intermediate monocytes, to high intensity in classical monocytes. DNAM-1 leads to clear separation of 2 populations from each subset. In yo T cells, the use of DNAM-1, TIGIT, CD27, GPR56, TIM-3, and CD161 leads to compartmentalization of the initially defined CD45RA/CCR7 subsets. For example, CCR7^−^CD45RA^+^ yo T cells show high expression of TIGIT and GPR56, in contrast to CCR7^−^CD45RA^+^ yo T cells that have increased levels of CD27 and DNAM-1. As shown for other subsets, DNAM-1 was identified as a highly informative marker to subclassify populations, it can be used in combination with CD3S to distinguish 2 subsets of basophils, and with CD25 for ILC subsets. CD1c^+^ DCs can be subclassified based on the presence or absence of TIM-3. Similarly, NK cell can be divided into additional subsets according to CD161 expression. No additional subclassification was identified for B cells. Manual analysis for highly parametric panels can be time consuming and cannot effectively capture the variety of subpopulations present in the sample. To explore the heterogeneity of the T cell memory pool, FlowSOM clustering and UMAP dimensionality reduction analysis were performed separately in CD4^+^ T cells, CDS^+^ T cells, and NKT-like T cells, with identification of 35 distinct metaclusters per subpopulation. We present the computationally derived cluster analogues of manually gated established subsets for CD4 and CDS T cells, with a phenotype description included in Online Figures 20-25, with comparisons across donors, populations, fixation conditions, and instruments.

We report the development of a high quality optimized 45-color panel for in-depth surface characterization of activation and exhaustion in the lymphoid compartment, that can highly resolve all subsets describe in OMIP-069 and additional subsets including MAIT, TRM, TSCM, TN, TCM, and TEM subpopulations (TEX, early, early-like, intermediate, and terminal), with further refinement using activation, exhaustion, and inhibitory markers. Understanding of memory T cell generation, function, adaptability, compartmentalization, lifetime persistence, and senescence requires definition of distinct populations by using single cell analysis technologies and contributes to understanding disease and treatment processes.

This panel highlights why FSFC continues to be a technological breakthrough for deeper explorations of immune compartments. Both panels can be applied for a variety of immunological studies in the context of immunotherapy, cancer monitoring, drug, and vaccine development, as well as biomarker discovery. We were able to build a high quality 45-color panel that preserves the biological identity and resolution of all populations of interest, and furthermore, which can be used to sort populations of interest using an optically compatible cell sorter (Online Figure 26). Peripheral blood T cells represent a small percentage of the total T cell compartment, most residing within tissues, thus additional testing would be necessary to identify the usage of this panel in other sample types or tissues.

**OMIP xxx Table 1..** Summary table for application of OMIP.

|  |  |
| --- | --- |
| PURPOSE | Deep subset profiling of immune cells to include subsets of T, B, NK, NKT, monocyte, and dendritic cells, and in-depth memory T cell evaluation |
| Species | Human |
| Cell type | PBMCs |
| Cross references | OMIP-003, OMIP-004, OMIP-006, OMIP-013, OMIP-015, OMIP-017, OMIP-021, OMIP-023, OMIP-024, OMIP-029, OMIP-030, OMIP-033, OMIP-034, OMIP-039, OMIP-042, OMIP-044, OMIP-050, OMIP-051, OMIP-058, OMIP-060, OMIP-063, OMIP-069, OMIP-070, OMIP-071, OMIP-074, OMIP-077, OMIP-078, OMIP-080, OMIP-084, OMIP-087, OMIP-090, OMIP-091, and OMIP-094 |

**Table 2.** Reagents Used in OMIP-xxx.

| Specificity | Fluorochrome | Clone | Purpose |
| --- | --- | --- | --- |
| **Leukocytes** |  |  |  |
| CD45 | PerCP | 2D1 | Leukocyte definition |
| **B cells** |  |  |  |
| CD19 | Spark NIR 685 | HIB19 | B cell definition |
| CD20 | Spark Violet 538 | 2H7 | B cell definition |
| CD24 | cFluor YG610 | SN3 | B cell differentiation |
| IgD | cFluor BYG750 | IgD26 | B cell differentiation |
| IgG | BV605 | G18-145 | B cell differentiation |
| IgM | BV570 | MHM-88 | B cell differentiation |
| **Monocytes** |  |  |  |
| CD14 | Spark Blue 550 | 63D3 | Monocyte definition |
| CD16 | BUV496 | 3G8 | Monocytes and NK cell differentiation |
| CD11b | Spark UV 387 | ICRF44 | Monocytes, dendritic cells and some lymphocyte differentiation |
| **Dendritic cells** |  |  |  |
| CD1c | Alexa Fluor 647 | L161 | Dendritic cell differentiation |
| CD123 | Super Bright 436 | 6H6 | Dendritic cell differentiation |
| CD11c | eFluor 450 | 3.9 | Dendritic cell definition |
| **NK cells** |  |  |  |
| CD56 | cFluor YG584 | 5.1H11 | NK cell identification and γδ T cell differentiation |
| CD2 | PerCP/Cy5.5 | TS1/8 | T cell and NK cell differentiation |
| CD337 | PE/Dazzle 594 | P30-15 | NK cell differentiation |
| CD314 | BUV615 | 1D11 | NK cell differentiation |
| **T cells** |  |  |  |
| CD3 | BV510 | SK7 | T cell and NKT-like cell definition |
| CD4 | PerCP/Fire 806 | SK3 | T helper T cell and NKT-like cell definition |
| CD8 | BUV805 | SK1 | Cytotoxic T cell, NK cell, and NKT-like cell definition |
| CD25 | cFluor BYG710 | 4000 | Regulatory T cell definition |
| TCRγδ | PerCP-Vio 700 | REA591 | γδ T cell definition |
| **T cell Memory** |  |  |  |
| CCR7 | BV421 | G043H7 | T cell differentiation and memory status |
| CD27 | APC-H7 | M-T271 | T cell and B cell differentiation and memory status |
| CD45RA | BUV395 | 5H9 | T cell differentiation and memory status |
| CD95 | PE/Cy5 | DX2 | T cell differentiation |
| CD127 | cFluor R720 | A019D5 | T cell differentiation and ILC identification |
| CD28 | BV650 | CD28.2 | T cell and NK cell differentiation |
| GPR56 | PE | CG4 | T cell differentiation and memory status |
| DNAM-1 | Real Blue 780 | DX11 | Monocyte, dendritic cell, T cell, and B cell differentiation |
| CD103 | PE/Fire 640 | Ber-ACT8 | Intraepithelial lymphocytes and regulatory T cells |
| CD161 | APC | HP-3G10 | T cell differentiation and memory status |
| KLRG1 | APC/Fire 810 | SA231A2 | T cell differentiation and memory status |
| **T cell Activation/Differentiation** |  |  |  |
| HLA-DR | BV480 | L203 | T cell and monocyte activation, and dendritic cell identification |
| CD38 | BUV737 | HB7 | Monocyte, dendritic cell, T cell, and B cell differentiation |
| CD39 | BUV661 | TU66 | B cell, T regulatory, and monocyte differentiation |
| CCR5 | BUV563 | 2D7/CCR5 | Monocyte, dendritic cell, T cell, and B cell differentiation |
| CCR6 | BV711 | G034E3 | T cell and B cell differentiation |
| CXCR3 | PE/Cy7 | G025H7 | Dendritic cell, T cell, and B cell differentiation |
| CXCR5 | BV750 | RF8B2 | T cell differentiation |
| **T cell Exhaustion** |  |  |  |
| CD57 | cFluor B532 | HNK-1 | T cell and NK cell differentiation and memory status |
| TIM-3 | BB515 | 7D3 | T cell inhibitory receptor |
| TIGIT | PE/Fire 810 | A15153G | T cell inhibitory receptor |
| PD-1 | BV785 | EH12.2H7 | T cell inhibitory receptor |
| **Other** |  |  |  |
| Viability | LIVE/DEAD Blue | - | Viability |

Abbreviations: APC, allophycocyanin; Spark UV, Spark ultraviolet; BUV, brilliant ultraviolet; BV, brilliant violet; BB, brilliant blue; FITC, fluorescein; PE, phycoerythrin; PerCP, Peridinin chlorophyll protein; and cFluor YG, yellow green; B-blue, R-red.

## Supporting information

Online Materials 45 Color Full Spectrum Panel

## SIMILARITY TO PUBLISHED OMIPS

Similar OMIPs with emphasis on broad human immune system evaluation have been published (OMIPs -015, -023, -024, -030, -033, -034, -042, -050, -05S, -063, -069, -071, -077, -07S, -0S0, and -094). The panel partially overlaps with OMIPs dedicated to evaluating T cells (OMIPs -004, -006, -013, -015, -017, -021, -030, -060, -0S4, -0S7, -090, and -091), dendritic cells (OMIP-044); monocytes (OMIP-0S3), B cells (OMIPs -003, -033, -051, and -074); ILCs OMIP-0S2), and NK cells (OMIPs -029, -039, and -070).

## STATEMENT OF ETHICAL USE OF HUMAN SAMPLES

Human PBMCs used in this study were obtained from AllCells Alameda, 1301 Harbor Bay Parkway, Suite 200, Alameda, CA 94502, Phone: (510) 726-2700. Ethical review and regulatory compliance were conducted by Alpha Independent Review Board, 1001 Avenida Pico, Suite C #497, San Clemente, CA 92673, (SSS) 265-5766 (toll free) under Protocol number: 7000-SOP-045.

## ACKNOWLEDGMENTS

The authors would like to thank Patrick C. Duncker for careful review of the manuscript.

## AUTHOR CONTRIBUTIONS

Lily M. Park: data acquisition; data curation; formal analysis; methodology; validation and visualization. Joanne Lannigan: Data curation; writing and editing. Quentin Low: data acquisition; data curation and formal analysis. Maria C. Jaimes: conceptualization; data curation; methodology; project administration; resources; supervision; validation; writing review and editing. Diana L. Bonilla: Conceptualization; data acquisition; data curation; formal analysis; visualization; validation and writing.

