## Supplementary material for "45-Color Full Spectrum Flow Cytometry Panel for Deep Immunophenotyping of the Major Lineages Present in Human Peripheral Blood Mononuclear Cells with Emphasis on the T cell Memory Compartment": Online Materials 45 Color Full Spectrum Panel

### Online Information

### Table of Contents

1. Instrument configuration and instrument setup
2. Fluorochrome selection
    a. Fluorochrome replacements from OMIP-069
    b. Identification of new fluorochromes to add to selection from OMIP-069
    c. Initial evaluation of 45-color combination using CD4 stained cells
3. Panel design strategy
4. 45-color assay optimization
    a. Antibody titrations
    b. Reference controls optimization
    c. Evaluation of unmixing accuracy in multicolor samples
    d. Spillover spread assessment
    e. Individual marker resolution assessment in single stained vs. multicolor samples
    f. Optimization of staining procedure
    g. Assessment of resolution of markers assigned to highly overlapping fluorochromes
5. 45-color assay performance
    a. Manual gating and cellular subsets identified across donors
    b. Clustering and dimensionality reduction analysis

   c. Impact of paraformaldehyde fixation on assay performance
   d. Assessment of assay performance across analyzers and sorter with the same optical configuration
   e. Sorting of T cell memory subsets and functional characterization by intracellular cytokine staining
6. Sample preparation and staining protocol
7. References

### 1. Instrument configuration and instrument setup

This panel was developed on a Cytek Aurora (Cytek Biosciences, Fremont, California) equipped with 5 lasers (355, 405, 488, 561, 640 nm) and 64 detectors as described in OMIP-069 (1) (**Online Table 1**).

Settings provided by the manufacturer (referred to as CytekAssaySetting in SpectroFlo® software) were used as a starting point for instrument setup. When those settings were developed by the manufacturer, there were no fluorochromes commercially available with a peak emission in the first detector of the UV module (center wavelength 372 nm). For this reason, the recommended setup was checked using CD4 cells stained with all commercially available fluorochromes excited by the UV laser, including Spark UV 387. The following criteria were used to determine the optimal UV1 gain: 1. fluorochrome spectra showing expected peak of emission and full emission profile 2. optimal resolution calculated using stain index 3. minimal spillover spread with fluorochromes emitting in neighboring channels (BUV395). For this assessment Peripheral Blood Mononuclear Cells (PBMCs) were stained with CD4 antibodies conjugate to Spark UV 387 and all BUV fluorochromes. CD4 Spark UV 387 stain index was doubled when using a 2-fold UV1 gain increase. Increasing the UV1 gain further lead to an improvement in resolution, but also to higher spread from Spark UV 387 into BUV395, and changes in the expected BUV395 peak emission (data not shown). We settled at increasing the UV1 gain by 2-fold and the new settings were saved as “CytekAssaySetting 2x UV1” and used throughout the development of the panel.

| Laser | Laser wavelength (nm) | Laser power (mW) | Laser type | Detector name | Detector type | Center wavelength (nm) | Bandwidth (nm) | Fluorochrome |
|---|---|---|---|---|---|---|---|---|
| Ultraviolet | 355 | 20 | Solid state | UV1 | APD | 372.5 | 15 | Spark UV 387 |
| | | | | UV2 | APD | 387.5 | 15 | BUV395 |
| | | | | UV3 | APD | 427.5 | 15 | |
| | | | | UV4 | APD | 443 | 15 | |
| | | | | UV5 | APD | 458 | 15 | |
| | | | | UV6 | APD | 473 | 15 | LIVE/DEAD Blue |
| | | | | UV7 | APD | 514 | 28 | BUV496 |
| | | | | UV8 | APD | 542 | 28 | |
| | | | | UV9 | APD | 581.5 | 31 | BUV563 |
| | | | | UV10 | APD | 612.5 | 31 | BUV615 |
| | | | | UV11 | APD | 664 | 27 | BUV661 |
| | | | | UV12 | APD | 691.5 | 28 | |
| | | | | UV13 | APD | 720 | 29 | |
| | | | | UV14 | APD | 749.5 | 30 | BUV737 |
| | | | | UV15 | APD | 779.5 | 30 | |
| | | | | UV16 | APD | 811.5 | 34 | BUV805 |
| Violet | 405 | 100 | Solid state | V1 | APD | 427.5 | 15 | BV421 |
| | | | | V2 | APD | 443 | 15 | Super Bright 436 |
| | | | | V3 | APD | 458 | 15 | eFluor 450 |
| | | | | V4 | APD | 473 | 15 | |
| | | | | V5 | APD | 508 | 20 | BV480 |
| | | | | V6 | APD | 524.5 | 17 | |
| | | | | V7 | APD | 541.5 | 17 | BV510 |
| | | | | V8 | APD | 580.5 | 19 | Spark Violet 538<br>BV570 |
| | | | | V9 | APD | 598 | 20 | |
| | | | | V10 | APD | 615 | 20 | BV605 |
| | | | | V11 | APD | 664 | 27 | BV650 |
| | | | | V12 | APD | 691.5 | 28 | |
| | | | | V13 | APD | 720 | 29 | BV711 |
| | | | | V14 | APD | 749.5 | 30 | BV750 |
| | | | | V15 | APD | 779.5 | 30 | BV785 |
| | | | | V16 | APD | 811.5 | 34 | |
| Blue | 488 | 50 | Solid state | B1 | APD | 508 | 20 | BB515 |
| | | | | B2 | APD | 524.5 | 17 | cFluor B532 |
| | | | | B3 | APD | 541.5 | 17 | Spark Blue 550 |
| | | | | B4 | APD | 580.5 | 19 | |
| | | | | B5 | APD | 598 | 20 | |
| | | | | B6 | APD | 615 | 20 | |
| | | | | B7 | APD | 661 | 17 | |
| | | | | B8 | APD | 679 | 18 | PerCP |
| | | | | B9 | APD | 697 | 19 | PerCP/Cy5.5 |
| | | | | B10 | APD | 717 | 20 | PerCP-Vio700 |
| | | | | B11 | APD | 738 | 21 | |
| | | | | B12 | APD | 760 | 23 | |
| | | | | B13 | APD | 783 | 23 | |
| | | | | B14 | APD | 811.5 | 34 | Real Blue 780<br>PerCP/Fire 806 |
| Yellow green | 561 | 50 | Solid state | YG1 | APD | 577 | 20 | YG584<br>PE |
| | | | | YG2 | APD | 598 | 20 | |
| | | | | YG3 | APD | 615 | 20 | PE-Dazzle 594<br>cFluor YG610 |
| | | | | YG4 | APD | 661 | 17 | PE/Fire 640 |
| | | | | YG5 | APD | 679 | 18 | PE-Cy5 |
| | | | | YG6 | APD | 697 | 19 | |
| | | | | YG7 | APD | 720 | 29 | cFluor BYG710 |
| | | | | YG8 | APD | 749.5 | 30 | cFluor BYG750 |
| | | | | YG9 | APD | 779.5 | 30 | PE/Cy7 |
| | | | | YG10 | APD | 811.5 | 34 | PE/Fire 810 |
| Red | 640 | 80 | Solid state | R1 | APD | 661 | 17 | APC |
| | | | | R2 | APD | 679 | 18 | Alexa Fluor 647 |
| | | | | R3 | APD | 697 | 19 | Spark NIR 685 |
| | | | | R4 | APD | 717 | 20 | cFluor 720 |
| | | | | R5 | APD | 738 | 31 | |
| | | | | R6 | APD | 760 | 23 | |
| | | | | R7 | APD | 783 | 23 | APC-H7 |
| | | | | R8 | APD | 811.5 | 34 | APC/Fire 810 |

**Online Table 1. Instrument Optical Configuration**
List of laser wavelength, power, and type, detector name, range and fluorochrome measured for each detector. The panel detailed within this OMIP was developed for a Cytek Aurora equipped with 5 lasers and 64 detectors. Avalanche Photodiode Detectors (APDs) are used.

### 2. Fluorochrome selection

For fluorochrome selection, the 40 fluorochromes described in OMIP-069 were used as a starting point. Seven fluorochromes were replaced due to issues with reagent performance and/or availability, which were discovered after the panel publication in 2020. After identifying the most efficient fluorochrome replacements, five additional fluorochromes were selected to further expand the panel. The new fluorochromes were chosen to maximize resolution of new markers, without perturbing the performance of the original conjugates.

#### a. Fluorochrome replacements from OMIP-069

Since the publication of OMIP-069, the panel has been adopted in laboratories worldwide (personal communications, and (2,3)). Throughout its adoption, challenges that limited the ability to use the full panel have been reported. Those challenges can be grouped into two main categories: reagent availability and reagent performance. Concerning reagent availability, four reagents CD20 Pacific Orange (Thermo Fisher, Cat. MHCD2030), CD25 Phycoerythrin (PE) PE-Alexa-Fluor 700 (Thermo Fisher, Cat. MHCD2524), CD24 PE-Alexa Fluor 610 (Thermo Fisher, Cat. MHCD2422), and CD127 Allophycocyanin (APC) APC-R700 (BD Biosciences, Cat. 565185) were often on backorder, taking up to several months to become available. In addition, technical issues with individual reagent performance were reported to the authors that involved tandem fluorochrome degradation observed for HLA-DR PE/Fire 810 (BioLegend, Cat. 307683), and changes in the spectrum profile of TCRγδ Peridinin Chlorophyll protein (PerCP) PerCP-eFluor 710 (Thermo Fisher, Cat. 49-9959-42). Finally, it was also documented that spread between Fluorescein isothiocyanate (FITC) and BB515, the most challenging fluorochrome combination in the panel (Similarity Index 0.98; Spillover Spread Value of Brilliant Blue (BB)

BB515 into FITC 29), was not always consistent with that in the publication and could be higher, making it challenging to use that fluorochrome combination.

We identified fluorochromes which could be used to overcome the availability and performance issues of the seven reagents listed above. Five reagents had a direct fluorochrome replacement with new fluorochromes having similar emission spectrum (similarity index of 0.94 or higher) and also similar or higher resolution compared to the original fluor: Spark Violet 538 for Pacific Orange cFluor Blue Yellow Green (BYG) BYG710 for PE-Alexa Fluor 700; cFluor R720 for APC-R700 and PerCP-Vio 700 for PerCP-eFluor 710. **(Online Figure 1A).** cFluor Yellow Green (YG) YG610 was identified as an option to replace PE-Alexa Fluor 610, with low excitation by the blue laser, reducing the spread into blue laser excited fluorochromes with similar emission. Resolution was first assessed by measuring stain index of PBMCs labeled with anti-human CD4 conjugated to the new fluorochromes **[**not shown, see Commentary **Ref when available].**

To overcome the high spread between FITC and BB515, FITC was replaced by cFluor B532. We observed a significantly lower similarity index and lower spillover spread value between BB515 and cFluor B532 (similarity index of 0.98 for BB515-FITC vs 0.89 for BB515-cFluor 532) **(Online Figures 1, 2, and 4E)**. The spread impact of Spark Blue 550 is also **shown (Online Figure 4E).**

With careful PE/Fire 810 handling in sample staining, we did not observe stability issues during the development of OMIP-069. However, because of the performance reports we received regarding this reagent, we evaluated alternative fluorochromes to replace it. As mentioned before, we focused on fluorochromes with similar 561 nm excitation and far-red emission, stable to exposure to light and fixation, and with minimal spread impact into other fluorochromes. cFluor BYG750 was selected as a replacement **(Online Figure 1).**

1A

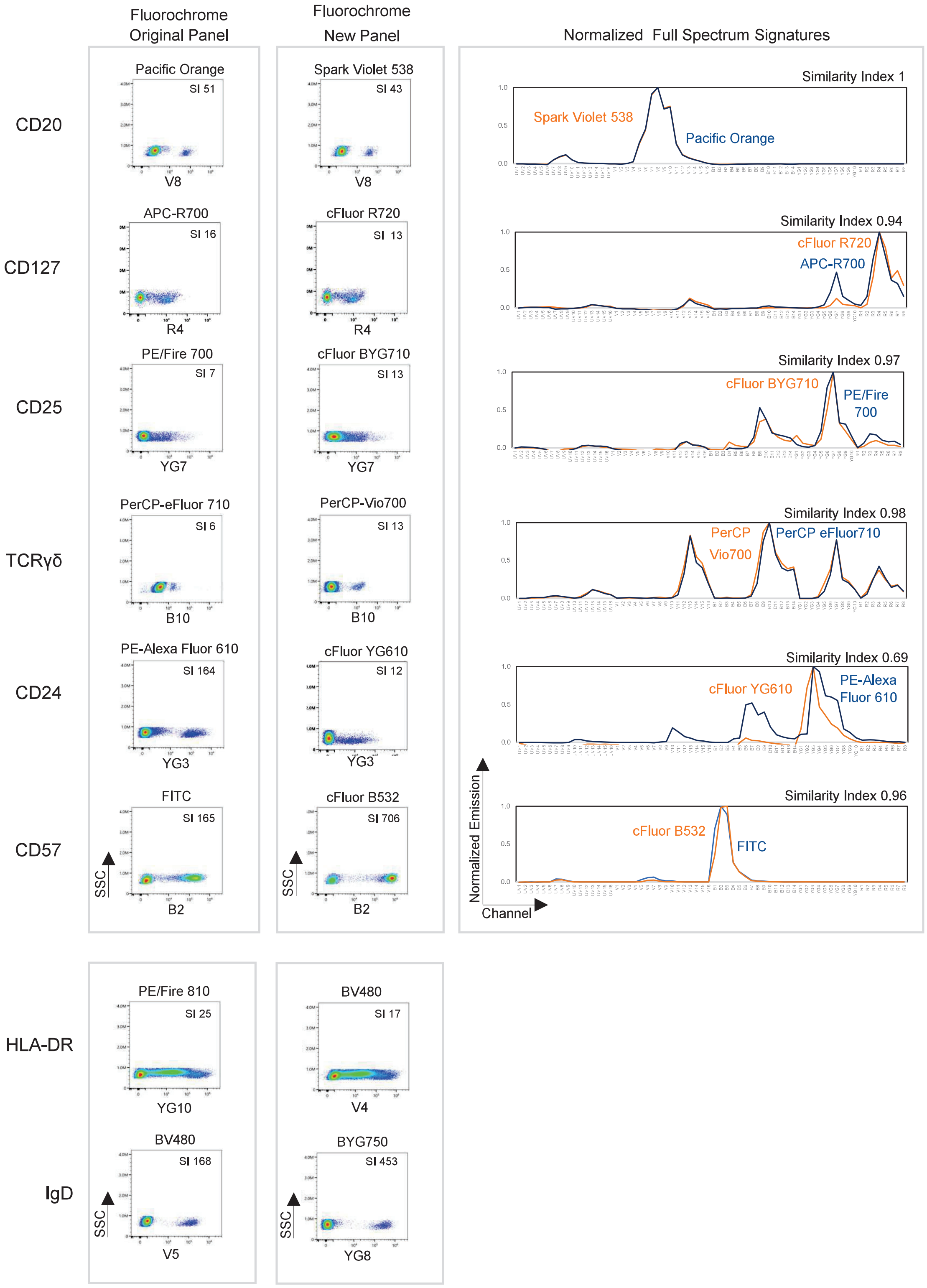
Fluorochrome Original Panel
Fluorochrome New Panel
Normalized Full Spectrum Signatures
CD20
Pacific Orange
SI 51
V8
Spark Violet 538
SI 43
V8
Similarity Index 1
Spark Violet 538
Pacific Orange
CD127
APC-R700
SI 16
R4
cFluor R720
SI 13
R4
Similarity Index 0.94
cFluor R720
APC-R700
CD25
PE/Fire 700
SI 7
YG7
cFluor BYG710
SI 13
YG7
Similarity Index 0.97
cFluor BYG710
PE/Fire 700
TCRγδ
PerCP-eFluor 710
SI 6
B10
PerCP-Vio700
SI 13
B10
Similarity Index 0.98
PerCP Vio700
PerCP eFluor710
CD24
PE-Alexa Fluor 610
SI 164
YG3
cFluor YG610
SI 12
YG3
Similarity Index 0.69
cFluor YG610
PE-Alexa Fluor 610
CD57
FITC
SI 165
SSC
B2
cFluor B532
SI 706
SSC
B2
Similarity Index 0.96
cFluor B532
FITC
Normalized Emission
Channel
HLA-DR
PE/Fire 810
SI 25
YG10
BV480
SI 17
V4
IgD
BV480
SI 168
SSC
V5
BYG750
SI 453
SSC
YG8

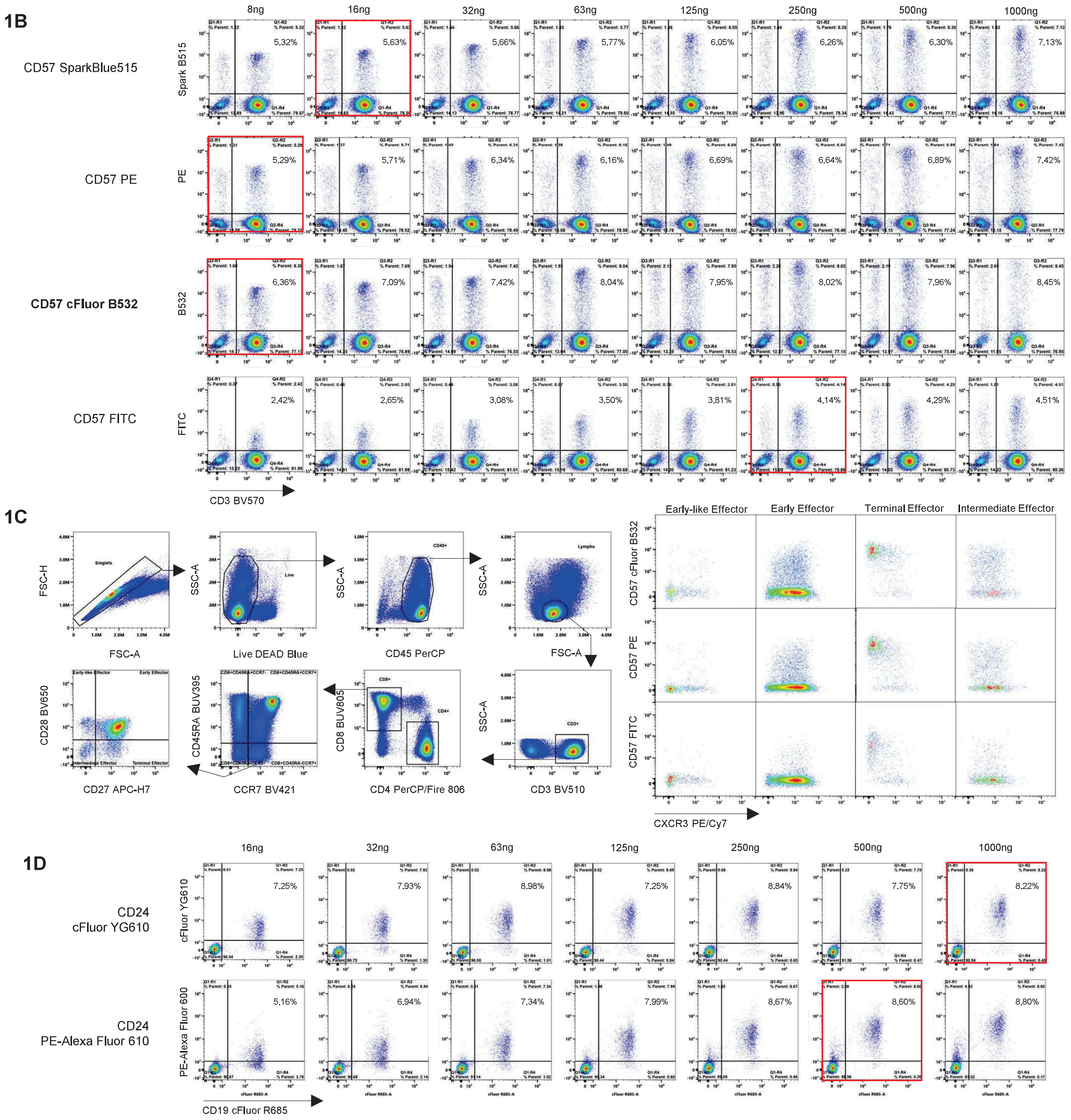

**Online Figure 1. Evaluation of OMIP-069 Replacement Reagents.**

Eight reagents were replaced from the original panel. Comparisons of the staining pattern (gated on lymphocytes) and resolution measured by Stain Index are presented, as normalized full spectrum profiles for reagents identified as direct replacements. (**A**) Similarity indices for each pair are shown, except for HLA-DR and IgD, which did not have a direct replacement with similar emission profile. (**B**) Data generated with the same anti-CD57 clone (HNK-1) for multiple conjugates (Spark Blue 515, PE, FITC, and cFluor B532), the one used in the panel. CD3 BV570 counter stain was used. Similar orthogonal populations were identified regardless the fluorochrome. Optimal titers shown in a red box. Similar staining patterns were observed between PE and the cFluor conjugate, but not for the FITC reagent, which identified lower percentage of CD57+ cells. (**C**) Multicolor panel tested to evaluate the use of CD57 cFluor B532 for identification of cell subsets in effector CD8 T cells. Data was compared against the same clone conjugated with PE or FITC, and similar patterns were observed. Events was gated on single CD45+CD3+CD8+CCR7+CD45RA-. Early-like Effector cells were defined as CD28+ CD27-, Early Effectors as CD28+ CD27+, Terminal Effectors as CD28- CD27+, and Intermediate Effectors as CD28- CD27-. Similar staining patterns were observed. (**D**) cFluor YG610 and PE-Alexa Fluor 610 CD24 conjugates (clone SN3) were tested to evaluate antibody performance. Cells were counter stained with CD19 cFluor R685. The same expression patterns were observed with both conjugates. Optimal titers are highlighted in red boxes.

Reagents of the same or equivalent clones conjugated to the new fluorochromes were then tested, first by antibody titrations and comparing performance at the single-color level to the original reagent. As shown in **Online Figure 1**, most of the new conjugates showed a similar resolution and pattern at optimal titer compared to the predicate reagent. However, the staining patterns for CD57 cFluor B532 and CD24 cFluor YG610 were slightly different compared to the original reagents. For CD57 cFluor B532, we evaluated its performance compared to conjugates to PE (considered as a Gold Standard), FITC and Spark Blue 515, all from the same clone. Cells were also stained with CD3 to evaluate expression in T and non-T cells. The staining pattern obtained with CD57 cFluor B532 was equivalent to the ones with the other reagents **(Online Figure 1B),** whereas the original CD57 FITC exhibited a lower % of positive cells in both compartments. We next investigated the differences in performance in a multicolor panel that allowed us to identify different T cell subsets. We found that CD57 cFluor B532 performed very similarly to the PE conjugate across T-cell subsets **(Online Figure 1C)**. Based on all these findings, CD57 cFluor B532 was identified as a suitable and superior reagent for the panel compared to the original CD57 FITC. Concerning CD24 cFluor YG610, co-staining with CD19, to identify B cells, and comparison with the original reagent revealed that the staining was specific and provided adequate resolution, and that the new reagent had less non-specific binding to non-B cells and hence included in the final panel **(Online Figure 1D)**.

The performance of the these newly identified fluorochromes was evaluated in fully stained samples comparing the two 40-color combinations (the one originally published in OMIP-069, and the one with the replacement fluorochromes). Similarity index matrices show the similarity for each pair of fluorochromes identified for use, accompanied by a reduction in complexity index for the revised panel. The new reagent performance in fully stained samples is shown in the **Online Figure 2** and in detail in the OMIP-069 v2 commentary in Cytometry Part A **[Ref when**

**2A**

Original Panel

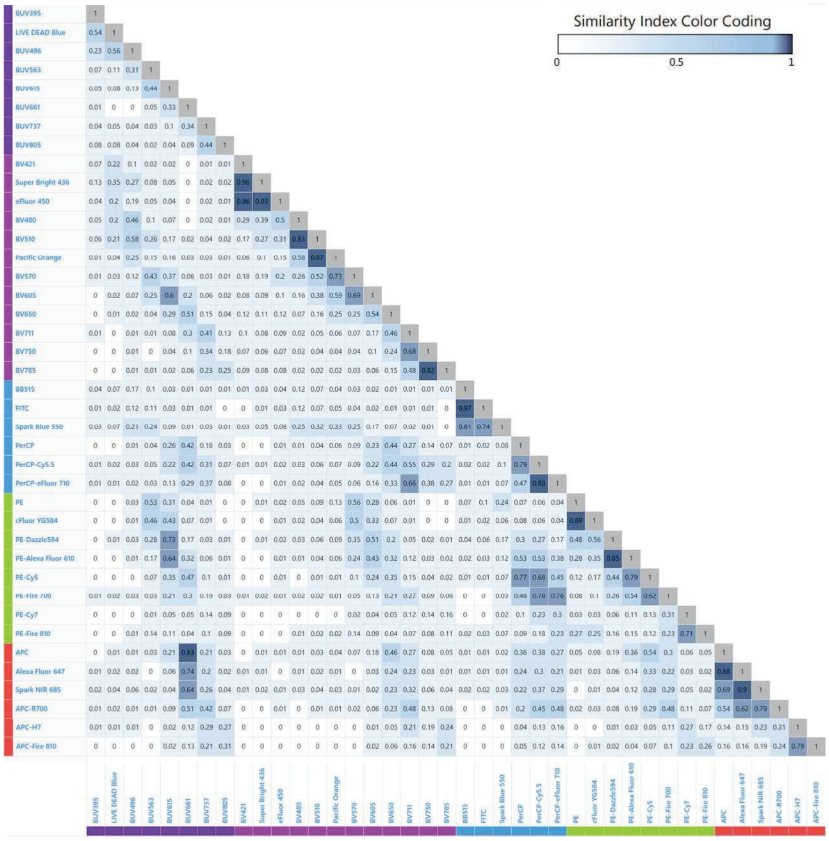

Complexity Index 54.97

Revised Panel

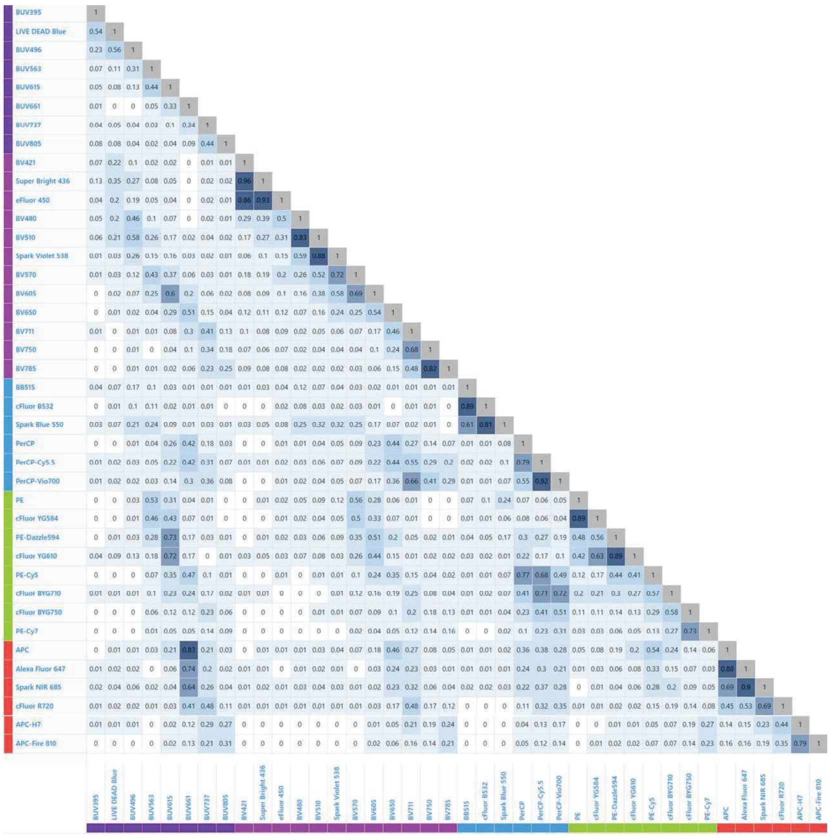

Complexity Index 50.81

**2B**

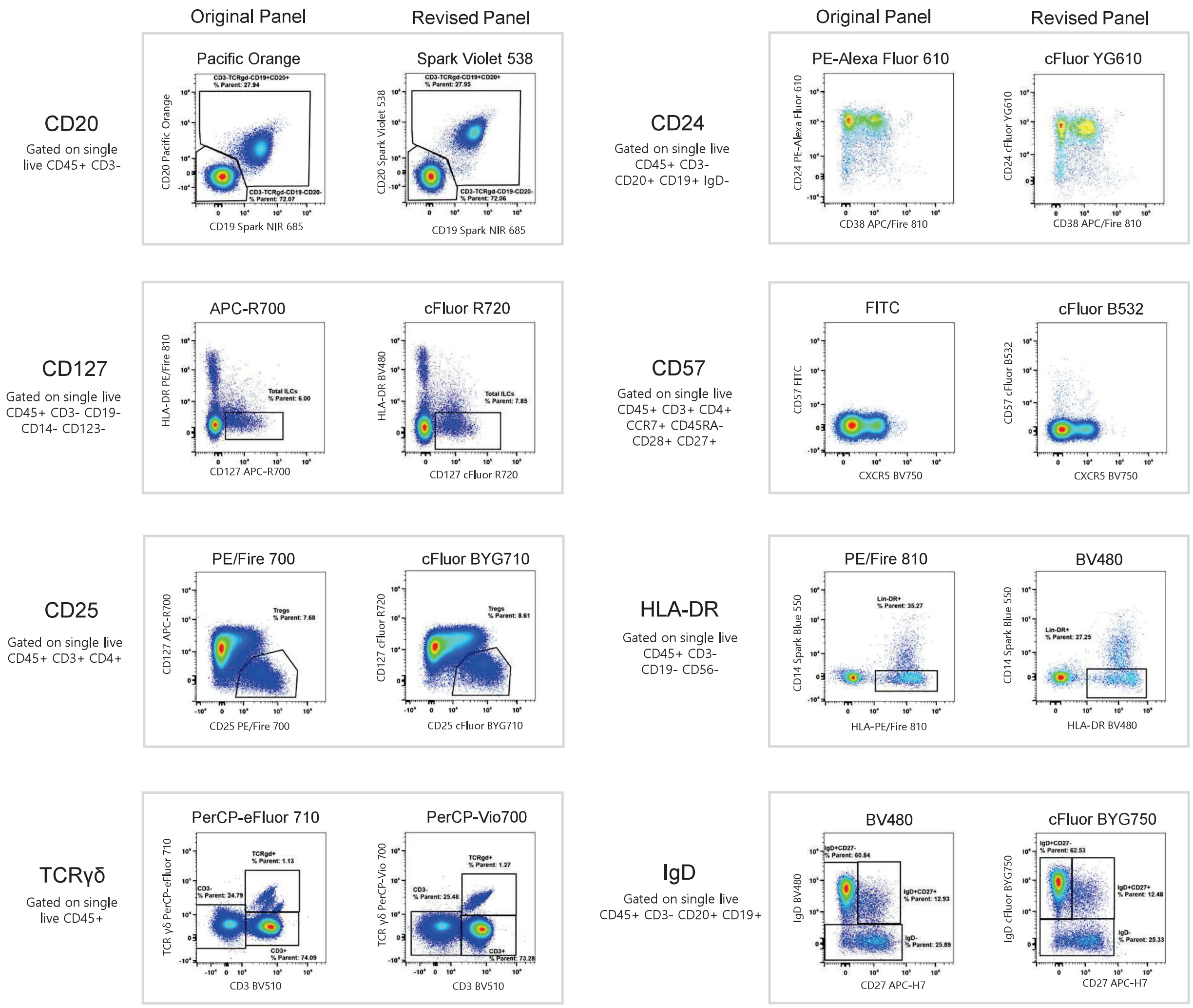

**Online Figure 2. Evaluation of OMIP-069 Replacement Reagents in the Multicolor Scenario.**
Fully stained samples were evaluated to compare the two 40-color combinations. (**A**) The Similarity Index Matrix (SIM) displays the numerical value for each pair of fluorochromes identified for use. Similarity matrices are shown for the Original Panel and Revised Panel. At the bottom of the matrices, the complexity indices are shown. A reduction in complexity is observed for the revised panel. Values were calculated with antigen-specific stained cells using SpectroFlo. (**B**) Representative plots showing the performance of newly introduced reagents compared to those from the original panel. Plots with gates are used for further subsetting, while plots without gates are the final population of interest.

**available].** Differences in the TCRγδ expression profile are expected since the PerCP-Vio 700 conjugate (Miltenyi clone REA771) recognizes the variable delta 2 chain of the gamma delta receptor complex, which is present in the main population of circulating γδ T cells.

#### b. Identification of additional fluorochromes compatible with OMIP-069 fluorochromes

We next aimed to identify additional fluorochromes that would be compatible with the 40 fluorochromes already selected. 33 fluorochromes from OMIP-069 were preserved, and 7 new fluorochromes were identified in the section above, 6 direct replacements (Spark Violet 538, cFluor R720, cFluor BYG710, PerCP-Vio 700, cFluor YG610, and cFluor B532), and 1 new fluorochrome addition (BYG750). A two-step approach was used to identify suitable additional fluorochromes. First, fluorochromes with unique spectra (similarity index lower than 0.98) were pre-selected using the database of 226 fluorochromes available through the Cytek® Cloud at the time this screening was done (August 2023). Of note, three families of fluorochromes were excluded from this screening: Quantum dots (Qdots), NovaFluors, and non-commercially available Brilliant Blue (BB). Qdots were not included due to known quenching issues requiring special handling (4). We have also observed aggregation, non-specific binding, and fluorochrome-fluorochrome interaction when using Qdots in complex panels. In addition, we have observed aggregation, non-specific binding, spread of the positive populations, and subpar performance characterized by loss of intensity in multicolor panels with NovaFluor fluorochromes. Finally, the reduced commercial availability of the Brilliant Blue custom conjugates, except for BB515 and BB700, limited our ability to do iterations of the panel in a timely manner.

Based on the initial screening, eleven fluorochromes with unique signatures were identified as candidates: Spark Ultraviolet (UV) UV387, StarBright UV445, StarBright UV795, cFluor V610, StarBright B580, StarBright B615, StarBright B810, Real Blue 780, PerCP/Fire 806, PE/Fire 640, and PE/Fire 810. Anti-human CD4 antibodies conjugated to the identified fluorochromes were used to generate similarity, complexity and stain indices, and spillover spread matrices. Each fluorochrome's characteristics and performance were fully characterized individually as well as in combination with the already selected fluors from the revised OMIP-069. First, we found issues related to high complexity index and spillover spread. StarBright B580 and StarBright B615 lead to a significant increase in the complexity index with higher spread into fluorochromes with similar emission. Next, we found that StarBright B810 and PerCP/Fire 806 introduced significant spread into each other, as well StarBright UV795 and BUV805, impacting multicolor performance. Because of clone and reagent availability, PerCP/Fire 806 and BUV805 were preferred over StarBright B810 and StarBright UV795, respectively. In addition, issues related to brightness were found: CD4 StarBright UV445 showed significant loss in brightness between the single stained control and when the reagent was used in combination with other fluorochromes. cFluor V610, a fluorochrome with low stain index, was not selected since the requirement was an additional bright fluorochrome to be paired with a tertiary antigen. Results of the testing of some reagents not included in the final panel are shown, emphasizing if they had optimal or suboptimal performance in single or multicolor evaluations **(Online Figure 3 and Online Table 2).**

To accommodate the addition of new markers, and since PE/Fire 810 fulfilled all the needed characteristics to be included in this fluorochrome combination, it was reincorporated into the panel. However, since stability issues had been reported from OMIP-069 adopters, we investigated fluorochrome stability. Fresh PBMCs were stained with either HLA-DR PE/Fire 810, CXCR3 PE/Cy7, or IgD cFluor BYG750, and acquired immediately. Then, cells were fixed with

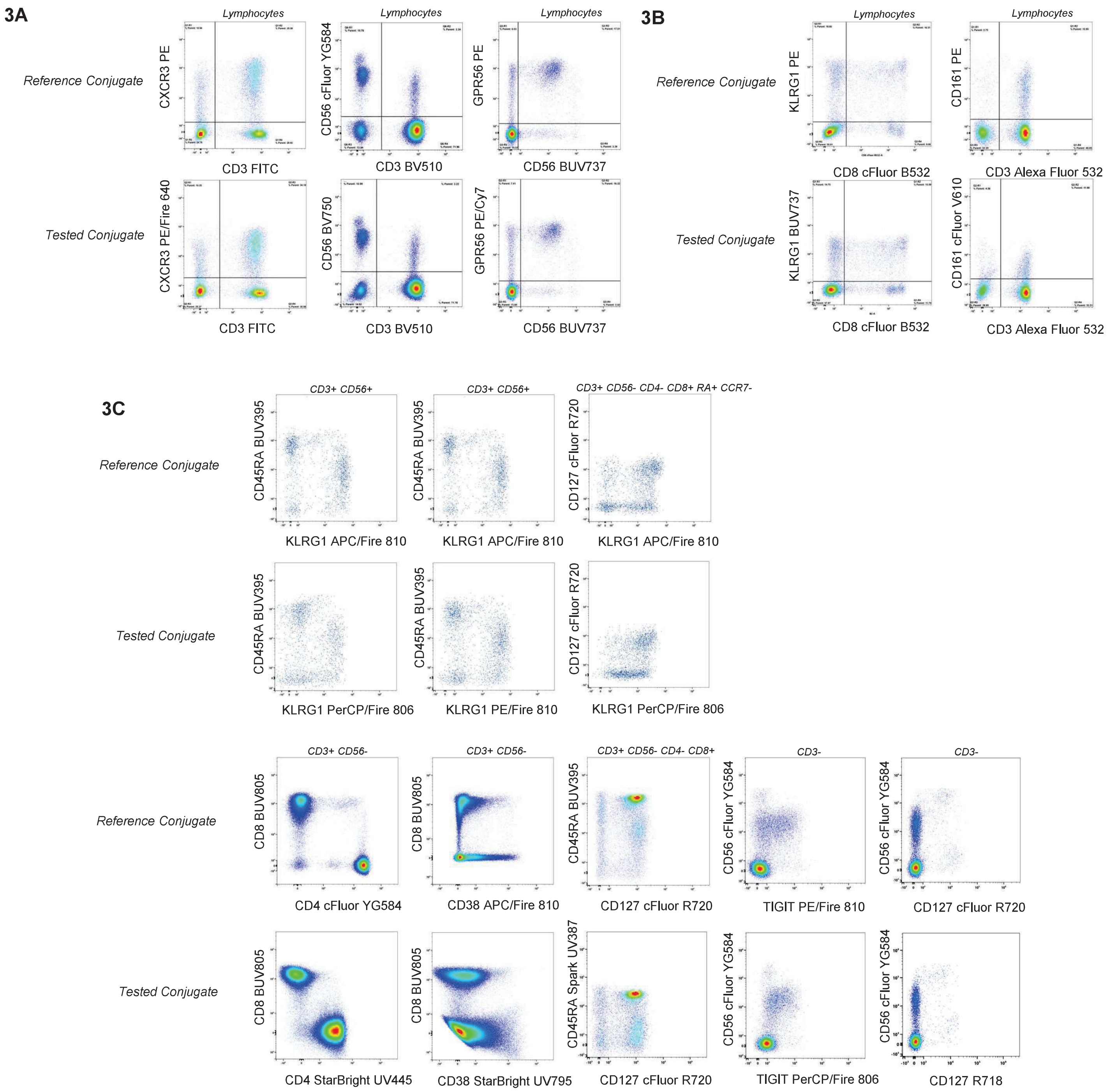

**Online Figure 3. Reagent conjugates tested but not used.**
(**A**) Examples of tested reagents which show adequate performance, when compared with a bright well characterized conjugate, but were not included in the final panel due to reagent selection, results for CXCR3 PE/Fire 640, CD56 BV750, and GPR56 PE/Cy7 are shown. (**B**) Examples of reagents with suboptimal performance either because of brightness or because of pattern differences when compared to a well characterized reagent, when used as single or co-stained. Results for KLRG1 BUV737 and CD161 cFluor V610 are shown. (**C**) Examples of reagents with suboptimal performance when used in fully stained samples. Results for KLRG1 PerCP/Fire 806, KLRG1 PE/Fire 810, CD4 StarBright UV445, CD38 StarBright UV795, CD45RA Spark UV 387, TIGIT PerCP/Fire 806, and CD127 R718 are shown. Information about reagent specificity, conjugate, clone, and catalog number are provided in the online table 2, as well as a summary of the testing results.

| | Specificity | Fluorochrome | Clone | Catalog # | Vendor | Testing Result |
|---|---|---|---|---|---|---|
| Adequate Performance Not Used in Final Panel | TIM-3 | PE/Fire 810 | F38-2E2 | 345059 | BioLegend | Good performance based on titration compared to reference, in the final panel PE/Fire 810 was assigned to TIGIT |
| | CD11b | PerCP/Cy5.5 | ICRF44 | 301328 | BioLegend | Good performance based on titration compared to reference, in the final panel we decided to reserve PerCP/Cy5.5 to CD2 |
| | CXCR3 | PE/Fire640 | G-25H7 | 353763 | BioLegend | Good performance based on titration compared to reference, in the final panel PE/Fire 640 was assigned to CD103 |
| | GPR56 | PECy7 | CG4 | 358206 | BioLegend | Good performance based on titration compared to reference, in the final panel GPR56 was assigned to PE |
| | CD38 | BV711 | HIT2 | 303528 | BioLegend | Good performance based on titration compared to reference, in the final panel CD38 was assigned to BUV737 |
| | CD127 | R718 | HIL-7R-M21 | 566968 | BD Biosciences | Good performance based on titration compared to reference and multicolor testing, in the final panel CD127 was assigned to cFluor R720 |
| | CD56 | BV750 | 5.1H11 | 362555 | BioLegend | Good performance based on titration compared to reference and multicolor testing, in the final panel CD56 was assigned to cFluor YG584 |
| Performance issues identified during titration and/or multicolor testing | KLRG1 | PE/Fire 810 | SA231A2 | 367733 | BioLegend | Based on multicolor testing, decreased resolution in some of the subsets of interests compared to reference |
| | KLRG1 | PerCP/Fire 806 | SA231A2 | 367747 | BioLegend | Based on multicolor testing, suboptimal resolution across subsets |
| | KLRG1 | BUV737 | 13F12F2 | 367-9488-42 | Thermo Fisher | Based on titration using co-staining, significant lower resolution compared to reference |
| | CD38 | StarBright UV795 | AT13/5 | MCA1019SBUV795 | Bio-Rad | Significant spread introduced by StarBright UV795 to CD8 BUV805. |
| | CD45RA | Spark UV387 | HI100 | 304179 | BioLegend | Based on titration and multicolor testing, suboptimal resolution |
| | CD4 | StarBright UV445 | RPA-T4 | MCA1267SBUV445 | Bio-Rad | Decreased intensity in multicolor testing |
| | CD161 | cFluor V610 | HP-3G10 | R0-00119 | Cytek Biosciences | Based on titration using co-staining, significant lower resolution compared to reference |
| | TIGIT | PerCP/Fire 806 | A15153G | 372749 | BioLegend | Based on titration and multicolor testing, supoptimal resolution |
| | CD319 | PerCP/Fire 806 | 162.1 | 331827 | BioLegend | Based on titration using co-staining, significant lower resolution compared to reference |
| | NKp80 | PE | 5D12 | 346706 | BioLegend | Based on multicolor testing, suboptimal resolution |

**Online Table 2. Tested Reagents not Included in the Panel.** Information about reagent specificity, conjugate, clone, catalog number, vendor, and testing results are provided. On top, the tested reagents which show adequate performance but were not included in the final panel. On the bottom, the reagents with suboptimal performance either because of brightness or because of significant differences when compared to a well characterized reagent.

either 1% or 4% Paraformaldehyde in PBS, exposed to light, and a time course was performed. All fluorochromes were stable for up to 2 hours under the two fixation conditions, i.e., no changes in spillover were found when compared to a fresh sample. However, PE/Fire 810 showed a significant increase of spillover into PE starting 2 hours after light exposure **[Commentary, Ref when available].** This increase was not seen with the other two tandems tested. Our findings indicate that PE/Fire 810 is more sensitive to light exposure than other tandem fluorochromes and that it needs to be handled with special care. We decided to include it as part of our expansion panel, since it remains stable under our normal staining conditions.

In summary, based on spectrum characteristics, brightness, spillover spread, and reagent availability, 5 fluorochromes were added to the modified 40-color combination: Spark UV 387, Real Blue 780, PerCP/Fire 806, PE/Fire 640, and PE/Fire 810.

#### c. Characterization of performance of 45-fluorophore combination

To fully characterize the identified fluorochrome combination for panel design, thawed PBMCs were stained with anti-human CD4 antibodies conjugated to the 44 identified fluorochromes, and the selected viability dye (data not shown). This allowed us to confirm the information provided by the Cytek Cloud tool and to accurately assess brightness ranking and spillover spread in the used instrument. The normalized spectra of the 45 fluorochromes is shown in **Online Figure 4A.** The vast majority of the fluorochromes have distinct peaks of emission, except for Spark Violet 538 and BV570, Real Blue 780 and PerCP/Fire 806, PE and cFluor YG584, and PE-Dazzle 594 and cFluor YG610. The similarity index matrix **(Online Figure 4B)** indicates that the pairs with the highest similarity were Brilliant Violet (BV) BV421 and Super Bright 436 (0.95), and PE/Cy5 and PE/Fire 640 (0.95).

We next assessed fluorochrome brightness by calculating stain index in CD4 stained samples, using the peak channel in raw data. We used this strategy to focus on the brightness of each fluorochrome, as unmixing the data will introduce spread. As shown in **Online Figure 4C,** this fluorochrome combination has a good balance between bright, mid, and dim fluorochromes. Importantly, fluorochromes emitting longer wavelengths (higher than 700 nm) are found among the brightest fluorochromes (Real Blue 780, PE/Cy5, cFluor BYG710, cFluor Red R720, etc.). NOTE: Panel design decisions based on brightness might not apply to other spectral platforms whose detectors do not exhibit the same sensitivity in the red and far-red areas of emission (5). The Complexity Index, a metric developed by Cytek Biosciences that correlates with the overall performance of the fluorophore combination, was 55. The lower the complexity index, the lower the probability that spillover spread will detrimentally impact panel performance, and in our experience a value of 55 is an acceptable number when this number of fluorochromes are used in combination on the Cytek Aurora. As a reference, the complexity index of the original 40-color combination of OMIP-069 was 53.72 (1).

The last part of the fluorochrome characterization using CD4 stained cells was the calculation of the spillover spread matrix (SSM) (5) **(Online Figure 4D).** SSM shows that the selected fluorochrome combination, when calculated for Aurora data, has a small number of combinations that require attention in terms spread. Of the 1,980 combinations of fluorochromes, only eleven had an SSM greater than 10. The fluorochrome combinations exhibiting a high level of spread (SSM values higher than 10) were: Spark UV 387 and Brilliant Ultraviolet (BUV) BUV395 (in both directions); BV421 into Super Bright 436; eFluor 450 into Super Bright 436; BV480 into BV510; PerCP and PerCP-Cy5.5 into PE/Cy5; PerCP into PE/Fire 640; PerCP-Vio 700 into PerCP-Cy5.5; and PE/Fire 640 and PE/Cy5 (in both directions). The fluorochromes introducing the most spread into others (highest row sum) are PerCP-Cy5.5 and PerCP-Vio 700; and the ones receiving the most spread (highest column sum) are Alexa Fluor 647 and PerCP-Cy5.5. NOTE:

**4A**

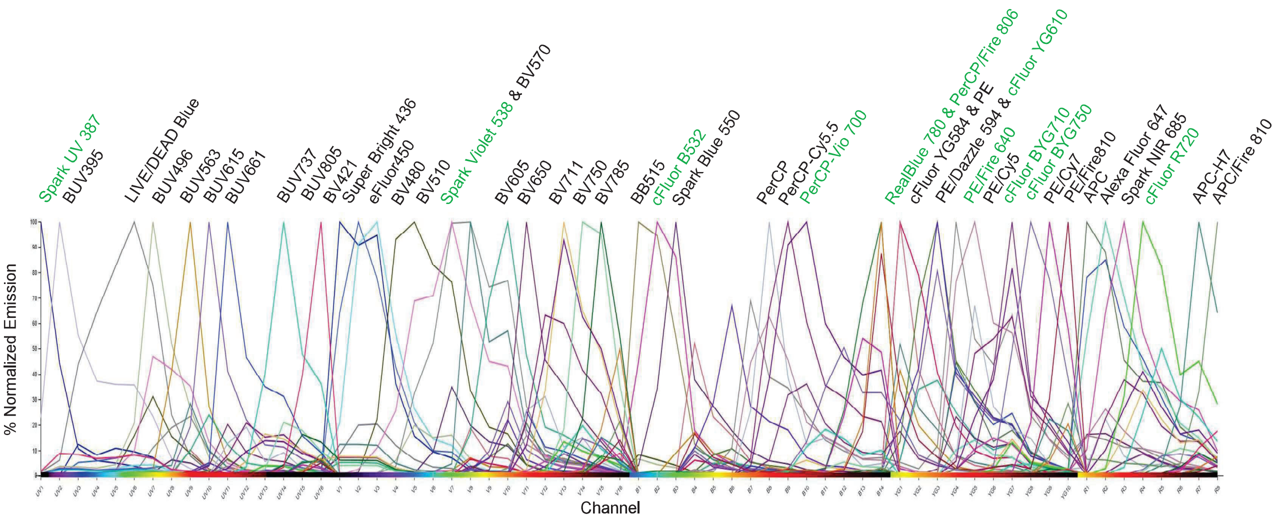
Spark UV 387
BUV395
LIVE/DEAD Blue
BUV496
BUV563
BUV615
BUV661
BUV737
BUV805
BV421
Super Bright 436
eFluor450
BV480
BV510
Spark Violet 538 & BV570
BV605
BV650
BV711
BV750
BV785
BB515
cFluor B532
Spark Blue 550
PerCP
PerCP-Cy5.5
PerCP-Vio 700
RealBlue 780 & PerCP/Fire 806
cFluor YG584 & PE
PE/Dazzle 594 & cFluor YG610
PE/Fire 640
PE/Cy5
cFluor BYG710
cFluor BYG750
PE/Cy7
PE/Fire810
APC
Alexa Fluor 647
Spark NIR 685
cFluor R720
APC-H7
APC/Fire 810
% Normalized Emission
Channel

**4B**

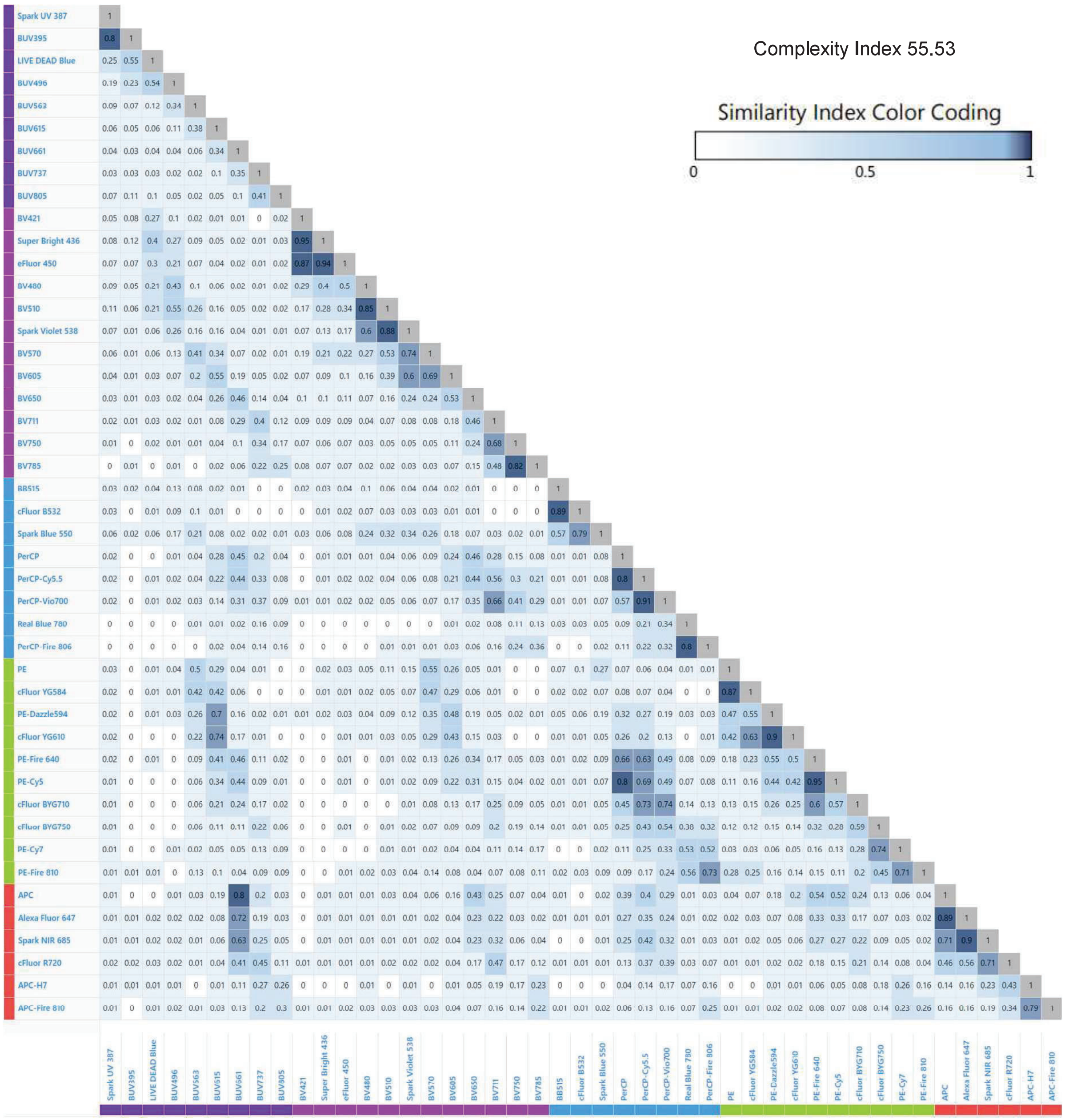
Complexity Index 55.53
Similarity Index Color Coding
0
0.5
1
Spark UV 387
BUV395
LIVE DEAD Blue
BUV496
BUV563
BUV615
BUV661
BUV737
BUV805
BV421
Super Bright 436
eFluor 450
BV480
BV510
Spark Violet 538
BV570
BV605
BV650
BV711
BV750
BV785
BB515
cFluor B532
Spark Blue 550
PerCP
PerCP-Cy5.5
PerCP-Vio700
Real Blue 780
PerCP-Fire 806
PE
cFluor YG584
PE-Dazzle594
cFluor YG610
PE-Fire 640
PE-Cy5
cFluor BYG710
cFluor BYG750
PE-Cy7
PE-Fire 810
APC
Alexa Fluor 647
Spark NIR 685
cFluor R720
APC-H7
APC-Fire 810

**4C**

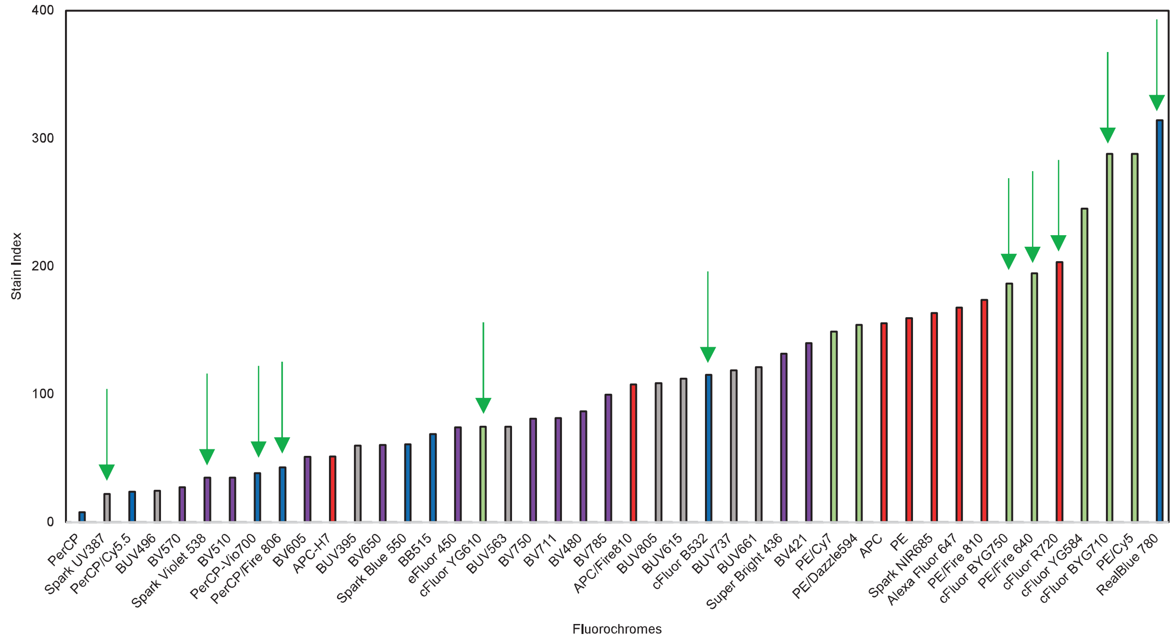

**4D**

| CD4 SSM | Spark UV387 | BUV395 | LIVE /EAD Blue | BUV496 | BUV563 | BUV615 | BUV661 | BUV737 | BUV805 | BV421 | Super Bright 436 | eFluor 450 | BV480 | BV510 | Spark Violet 538 | BV570 | BV605 | BV650 | BV711 | BV750 | BV785 | BB515 | cFluor B532 |
|---|---|---|---|---|---|---|---|---|---|---|---|---|---|---|---|---|---|---|---|---|---|---|---|
| Spark UV387 | | 10.32 | 3.46 | 0.00 | 0.00 | 0.00 | 0.00 | 0.34 | 0.00 | 0.00 | 0.00 | 0.00 | 0.00 | 0.00 | 0.00 | 0.80 | 0.00 | 0.45 | 0.00 | 0.00 | 0.48 | 0.00 | 0.00 |
| BUV395 | 12.02 | | 2.87 | 0.00 | 0.00 | 0.00 | 0.00 | 0.00 | 0.00 | 0.00 | 1.40 | 0.98 | 0.53 | 1.41 | 0.00 | 0.00 | 0.00 | 0.00 | 0.21 | 0.00 | 0.00 | 0.00 | 0.00 |
| LIVE/DEAD Blue | 2.10 | 3.55 | | 3.20 | 0.00 | 0.80 | 0.00 | 0.00 | 0.44 | 2.83 | 4.31 | 3.58 | 1.62 | 5.04 | 3.07 | 0.68 | 0.95 | 0.00 | 0.00 | 0.00 | 0.00 | 0.95 | 0.00 |
| BUV496 | 2.01 | 1.38 | 0.00 | | 1.91 | 0.90 | 0.20 | 0.33 | 0.26 | 0.68 | 1.59 | 0.00 | 1.81 | 5.72 | 4.95 | 1.66 | 0.77 | 0.44 | 0.36 | 0.00 | 0.37 | 1.18 | 1.65 |
| BUV563 | 1.89 | 1.35 | 0.00 | 0.00 | | 1.58 | 0.83 | 0.53 | 0.51 | 1.28 | 1.57 | 0.00 | 1.32 | 3.83 | 3.29 | 1.27 | 0.58 | 0.48 | 0.35 | 0.48 | 0.23 | 0.35 | 1.22 |
| BUV615 | 1.77 | 1.29 | 0.00 | 0.00 | 1.91 | | 2.02 | 1.56 | 1.41 | 0.00 | 0.00 | 0.00 | 0.53 | 1.37 | 0.00 | 0.00 | 1.25 | 1.15 | 0.72 | 0.72 | 0.80 | 0.25 | 0.27 |
| BUV661 | 1.75 | 1.14 | 0.00 | 0.00 | 0.00 | 0.52 | | 2.72 | 2.37 | 0.61 | 0.69 | 0.45 | 0.00 | 0.00 | 0.00 | 0.12 | 0.41 | 1.95 | 1.09 | 1.19 | 1.20 | 0.00 | 0.00 |
| BUV737 | 1.64 | 1.22 | 0.00 | 0.00 | 0.00 | 0.18 | 0.66 | | 4.70 | 0.00 | 0.00 | 0.00 | 0.18 | 1.10 | 0.00 | 0.00 | 0.00 | 0.30 | 1.34 | 1.68 | 1.91 | 0.00 | 0.00 |
| BUV805 | 2.92 | 2.26 | 0.00 | 0.00 | 0.00 | 0.00 | 0.21 | 1.31 | | 1.02 | 1.30 | 0.00 | 0.13 | 0.00 | 0.00 | 0.00 | 0.00 | 0.00 | 0.10 | 0.52 | 1.39 | 0.00 | 0.21 |
| BV421 | 0.63 | 0.68 | 0.72 | 1.27 | 0.92 | 0.24 | 0.00 | 0.00 | 0.00 | | 10.19 | 4.40 | 2.01 | 4.56 | 3.24 | 1.13 | 0.43 | 0.30 | 0.00 | 0.38 | 0.00 | 0.00 | 0.00 |
| Super Bright 436 | 0.00 | 0.09 | 0.00 | 1.22 | 0.81 | 0.16 | 0.20 | 0.00 | 0.00 | 6.07 | | 4.09 | 2.00 | 4.59 | 3.62 | 1.29 | 0.52 | 0.00 | 0.19 | 0.41 | 0.55 | 0.00 | 0.52 |
| eFluor 450 | 0.00 | 0.00 | 0.00 | 0.81 | 0.95 | 0.00 | 0.37 | 0.12 | 0.25 | 6.72 | 11.34 | | 2.48 | 6.12 | 5.08 | 1.70 | 0.54 | 0.00 | 0.00 | 0.00 | 0.00 | 0.43 | 0.00 |
| BV480 | 0.31 | 0.53 | 0.00 | 5.81 | 2.30 | 0.74 | 0.29 | 0.27 | 0.07 | 3.00 | 5.01 | 2.49 | | 10.69 | 8.98 | 3.48 | 1.65 | 0.74 | 0.37 | 0.00 | 0.21 | 1.65 | 1.88 |
| BV510 | 0.00 | 0.39 | 0.00 | 5.44 | 3.38 | 1.22 | 0.77 | 0.70 | 0.70 | 1.81 | 2.95 | 1.47 | 2.31 | | 8.34 | 3.84 | 2.10 | 1.01 | 0.31 | 0.84 | 0.90 | 0.42 | 0.00 |
| Spark Violet 538 | 0.64 | 0.26 | 0.00 | 0.00 | 1.96 | 0.99 | 0.47 | 0.15 | 0.44 | 0.00 | 0.00 | 0.00 | 1.30 | 6.51 | | 4.02 | 2.39 | 1.36 | 0.92 | 0.97 | 1.16 | 0.00 | 1.04 |
| BV570 | 0.56 | 0.34 | 0.00 | 0.00 | 2.41 | 1.08 | 0.62 | 0.58 | 0.52 | 2.92 | 4.44 | 1.67 | 0.96 | 2.59 | 2.26 | | 2.17 | 1.22 | 0.54 | 0.80 | 0.91 | 0.00 | 1.42 |
| BV605 | 0.00 | 0.00 | 0.00 | 0.00 | 1.45 | 2.20 | 1.16 | 1.03 | 0.87 | 1.74 | 2.36 | 0.77 | 0.42 | 0.00 | 1.18 | 1.22 | | 1.98 | 1.06 | 1.40 | 1.50 | 0.00 | 0.57 |
| BV650 | 0.00 | 0.11 | 0.00 | 0.00 | 0.00 | 0.48 | 1.92 | 1.19 | 1.03 | 1.81 | 2.82 | 0.99 | 0.75 | 2.39 | 2.01 | 0.86 | 0.94 | | 1.60 | 1.77 | 1.95 | 0.18 | 0.70 |
| BV711 | 0.00 | 0.00 | 0.00 | 0.00 | 0.00 | 0.20 | 0.60 | 2.24 | 2.04 | 1.88 | 2.82 | 1.12 | 0.45 | 0.00 | 0.00 | 0.33 | 0.00 | 0.66 | | 3.23 | 3.52 | 0.00 | 0.00 |
| BV750 | 0.31 | 0.00 | 0.00 | 0.00 | 0.00 | 0.09 | 0.13 | 2.75 | 2.45 | 1.32 | 1.91 | 0.00 | 0.45 | 1.19 | 1.21 | 0.00 | 0.00 | 0.34 | 1.25 | | 4.65 | 0.00 | 0.00 |
| BV785 | 0.25 | 0.30 | 0.00 | 0.00 | 0.00 | 0.00 | 0.07 | 1.19 | 2.73 | 1.66 | 2.83 | 0.99 | 0.44 | 0.76 | 1.00 | 0.00 | 0.00 | 0.00 | 0.52 | 2.15 | | 0.00 | 0.00 |
| BB515 | 0.00 | 0.00 | 0.00 | 0.67 | 0.54 | 0.15 | 0.14 | 0.00 | 0.00 | 0.48 | 0.58 | 0.28 | 0.38 | 2.30 | 3.21 | 1.49 | 0.30 | 0.00 | 0.16 | 0.13 | 0.18 | | 8.08 |
| cFluor B532 | 0.00 | 0.24 | 0.00 | 0.00 | 0.10 | 0.14 | 0.23 | 0.06 | 0.00 | 0.00 | 1.03 | 0.47 | 0.34 | 2.44 | 3.65 | 1.62 | 0.40 | 0.00 | 0.22 | 0.16 | 0.32 | 3.92 | |
| Spark Blue 550 | 0.00 | 0.12 | 0.00 | 0.00 | 0.72 | 0.43 | 0.46 | 0.25 | 0.00 | 0.94 | 1.10 | 0.44 | 0.66 | 3.61 | 4.60 | 2.50 | 1.02 | 0.95 | 0.55 | 0.55 | 0.82 | 1.91 | 4.47 |
| PerCP | 0.82 | 0.00 | 0.00 | 0.00 | 0.00 | 0.67 | 1.48 | 1.71 | 1.10 | 0.68 | 2.31 | 0.00 | 1.36 | 2.65 | 0.00 | 0.00 | 1.07 | 1.71 | 0.00 | 0.00 | 1.39 | 1.45 | 2.29 |
| PerCP/Cy5.5 | 0.00 | 0.73 | 0.00 | 0.00 | 0.00 | 0.41 | 1.36 | 2.01 | 1.75 | 0.00 | 0.00 | 0.00 | 0.00 | 0.62 | 1.61 | 0.69 | 0.59 | 2.50 | 2.66 | 2.42 | 3.11 | 1.01 | 1.22 |
| PerCP-Vio700 | 0.86 | 0.69 | 0.00 | 0.00 | 0.00 | 0.25 | 1.09 | 1.69 | 1.79 | 0.64 | 2.47 | 0.41 | 0.00 | 0.00 | 0.71 | 0.79 | 0.00 | 2.01 | 3.53 | 2.80 | 3.63 | 0.00 | 0.00 |
| Real Blue 780 | 0.00 | 0.21 | 0.00 | 0.00 | 0.00 | 0.14 | 0.14 | 0.63 | 1.57 | 0.14 | 0.00 | 0.00 | 0.22 | 0.35 | 0.68 | 0.36 | 0.09 | 0.12 | 0.48 | 1.62 | 3.55 | 0.58 | 1.01 |
| PerCP/Fire 806 | 0.32 | 0.22 | 0.00 | 0.00 | 0.00 | 0.00 | 0.31 | 0.38 | 1.73 | 0.14 | 0.00 | 0.00 | 0.00 | 0.77 | 0.49 | 0.00 | 0.00 | 0.51 | 0.16 | 1.10 | 2.92 | 0.00 | 0.00 |
| PE | 0.00 | 0.00 | 0.00 | 0.00 | 1.23 | 0.56 | 0.36 | 0.12 | 0.09 | 0.47 | 0.59 | 0.20 | 0.32 | 0.79 | 1.23 | 2.94 | 1.10 | 0.69 | 0.43 | 0.24 | 0.26 | 0.51 | 1.06 |
| cFluor YG584 | 0.21 | 0.16 | 0.00 | 0.00 | 0.28 | 0.42 | 0.20 | 0.00 | 0.07 | 0.00 | 0.00 | 0.00 | 0.00 | 0.00 | 0.44 | 0.86 | 0.72 | 0.59 | 0.34 | 0.38 | 0.41 | 0.29 | 0.94 |
| PE/Dazzle594 | 0.24 | 0.00 | 0.00 | 0.00 | 0.42 | 0.69 | 0.74 | 0.34 | 0.26 | 0.20 | 0.00 | 0.00 | 0.26 | 0.90 | 1.05 | 0.35 | 1.77 | 1.38 | 0.87 | 0.56 | 0.70 | 0.56 | 1.03 |
| cFluor YG610 | 0.79 | 0.44 | 0.00 | 0.00 | 0.00 | 0.56 | 0.42 | 0.24 | 0.12 | 0.00 | 0.00 | 0.00 | 0.00 | 2.75 | 2.74 | 0.60 | 0.94 | 1.20 | 0.60 | 0.00 | 0.68 | 0.00 | 1.42 |
| PE/Fire 640 | 0.00 | 0.00 | 0.00 | 0.00 | 0.00 | 0.30 | 0.96 | 0.66 | 0.52 | 0.56 | 0.00 | 0.00 | 0.27 | 0.82 | 1.07 | 0.40 | 0.59 | 2.09 | 1.69 | 1.04 | 1.25 | 0.22 | 0.24 |
| PE/Cy5 | 0.05 | 0.00 | 0.00 | 0.00 | 0.13 | 0.21 | 1.25 | 0.69 | 0.53 | 0.00 | 0.00 | 0.10 | 0.23 | 0.23 | 0.42 | 0.29 | 0.71 | 2.67 | 1.90 | 0.99 | 1.27 | 0.30 | 0.77 |
| cFluor BYG710 | 0.00 | 0.04 | 0.00 | 0.00 | 0.25 | 0.22 | 0.64 | 0.72 | 0.64 | 0.00 | 0.41 | 0.16 | 0.00 | 0.00 | 0.16 | 0.57 | 0.46 | 1.14 | 2.24 | 1.14 | 1.54 | 0.23 | 0.61 |
| cFluor BYG750 | 0.00 | 0.00 | 0.00 | 0.00 | 0.33 | 0.20 | 0.37 | 1.21 | 1.09 | 0.46 | 0.44 | 0.00 | 0.13 | 0.00 | 0.46 | 0.33 | 0.26 | 0.63 | 1.63 | 2.01 | 2.73 | 0.09 | 0.00 |
| PE/Cy7 | 0.00 | 0.09 | 0.00 | 0.00 | 0.00 | 0.00 | 0.19 | 0.56 | 1.11 | 0.31 | 0.00 | 0.00 | 0.00 | 0.00 | 0.00 | 0.17 | 0.00 | 0.25 | 0.47 | 1.11 | 3.11 | 0.00 | 0.32 |
| PE/Fire 810 | 0.08 | 0.17 | 0.00 | 0.00 | 0.00 | 0.22 | 0.26 | 0.27 | 1.53 | 0.39 | 0.00 | 0.00 | 0.40 | 0.00 | 0.13 | 0.71 | 0.31 | 0.42 | 0.44 | 0.92 | 2.83 | 0.19 | 0.49 |
| APC | 0.00 | 0.00 | 0.00 | 0.00 | 0.00 | 0.00 | 0.89 | 0.82 | 0.67 | 0.00 | 0.00 | 0.00 | 0.00 | 0.00 | 0.00 | 0.35 | 0.37 | 2.04 | 0.92 | 0.77 | 1.03 | 0.00 | 0.43 |
| Alexa Fluor 647 | 0.20 | 0.00 | 0.00 | 0.00 | 0.00 | 0.00 | 0.35 | 0.34 | 0.14 | 0.26 | 0.55 | 0.00 | 0.00 | 0.00 | 0.00 | 0.00 | 0.31 | 1.46 | 0.49 | 0.38 | 0.30 | 0.00 | 0.00 |
| Spark NIR 685 | 0.00 | 0.31 | 0.00 | 0.00 | 0.00 | 0.00 | 0.47 | 0.57 | 0.48 | 0.00 | 0.00 | 0.00 | 0.00 | 0.00 | 0.00 | 0.00 | 0.09 | 1.37 | 0.72 | 0.44 | 0.72 | 0.00 | 0.00 |
| cFluor R720 | 0.15 | 0.00 | 0.00 | 0.00 | 0.00 | 0.00 | 0.34 | 1.00 | 1.04 | 0.68 | 0.00 | 0.00 | 0.00 | 0.00 | 0.00 | 0.00 | 0.00 | 0.61 | 1.20 | 1.28 | 1.35 | 0.00 | 0.00 |
| APC-H7 | 0.00 | 0.00 | 0.00 | 0.00 | 0.00 | 0.00 | 0.00 | 0.50 | 1.40 | 0.00 | 0.00 | 0.00 | 0.39 | 0.80 | 0.00 | 0.00 | 0.00 | 0.33 | 0.00 | 0.89 | 1.83 | 0.46 | 0.00 |
| APC/Fire 810 | 0.00 | 0.00 | 0.00 | 0.00 | 0.00 | 0.00 | 0.00 | 0.37 | 1.79 | 0.00 | 0.00 | 0.00 | 0.00 | 0.23 | 0.00 | 0.00 | 0.00 | 0.56 | 0.43 | 0.69 | 1.46 | 0.00 | 0.00 |
| Sum | 32.54 | 18.32 | 3.59 | 18.42 | 22.01 | 16.92 | 23.16 | 31.45 | 38.41 | 41.67 | 66.98 | 25.07 | 24.64 | 76.91 | 70.88 | 36.61 | 25.79 | 36.58 | 32.65 | 37.48 | 57.18 | 17.14 | 33.86 |

| CD4 SSM | Spark Blue 550 | PerCP | PerCP/Cy5.5 | PerCP-Vio700 | Real Blue 780 | PerCP/Fire 806 | PE | cFluor YG584 | PE/Dazzle594 | cFluor YG610 | PE/Fire 640 | PE/Cy5 | cFluor BYG710 | cFluor BYG750 | PE/Cy7 | PE/Fire 810 | APC | Alexa Fluor 647 | Spark NIR 685 | cFluor R720 | APC-H7 | APC/Fire 810 | Sum |
|---|---|---|---|---|---|---|---|---|---|---|---|---|---|---|---|---|---|---|---|---|---|---|---|
| Spark UV387 | 0.00 | 0.60 | 1.44 | 1.15 | 0.00 | 0.27 | 0.00 | 0.00 | 1.38 | 1.14 | 1.22 | 0.99 | 0.00 | 0.54 | 0.32 | 0.00 | 0.00 | 0.00 | 0.31 | 0.00 | 0.47 | 0.00 | 25.69 |
| BUV395 | 0.00 | 0.00 | 0.00 | 0.00 | 0.26 | 0.00 | 0.74 | 0.00 | 0.00 | 0.00 | 0.00 | 0.00 | 0.28 | 0.00 | 0.22 | 0.65 | 1.14 | 1.79 | 1.39 | 0.00 | 0.25 | 0.00 | 14.11 |
| LIVE/DEAD Blue | 0.00 | 0.00 | 0.00 | 0.52 | 0.00 | 0.00 | 0.00 | 0.00 | 0.47 | 0.75 | 0.00 | 0.00 | 0.00 | 0.00 | 0.58 | 0.44 | 0.00 | 0.00 | 0.00 | 0.00 | 0.00 | 0.00 | 33.77 |
| BUV496 | 0.74 | 0.00 | 0.00 | 0.24 | 0.11 | 0.00 | 0.60 | 0.21 | 0.37 | 0.59 | 0.00 | 0.00 | 0.00 | 0.00 | 0.00 | 0.00 | 0.47 | 0.73 | 0.53 | 0.00 | 0.00 | 0.25 | 30.74 |
| BUV563 | 1.46 | 1.21 | 1.17 | 1.00 | 0.19 | 0.35 | 2.73 | 3.15 | 1.69 | 2.58 | 2.34 | 2.51 | 0.89 | 0.32 | 0.16 | 0.00 | 1.01 | 1.15 | 0.76 | 0.00 | 0.00 | 0.00 | 45.74 |
| BUV615 | 0.61 | 2.39 | 2.41 | 1.81 | 0.79 | 0.87 | 1.06 | 1.80 | 1.96 | 4.98 | 5.66 | 5.95 | 2.25 | 0.89 | 1.17 | 0.75 | 1.63 | 1.92 | 1.24 | 0.67 | 0.22 | 0.00 | 56.27 |
| BUV661 | 0.00 | 2.81 | 2.91 | 1.95 | 0.81 | 1.19 | 0.00 | 0.27 | 1.72 | 1.36 | 5.88 | 6.68 | 2.40 | 1.02 | 1.29 | 0.78 | 5.10 | 7.44 | 5.30 | 2.01 | 1.44 | 1.18 | 66.82 |
| BUV737 | 0.00 | 0.00 | 2.88 | 3.06 | 1.66 | 2.26 | 0.56 | 0.00 | 0.00 | 0.21 | 1.84 | 1.84 | 1.52 | 1.26 | 1.40 | 0.76 | 1.39 | 2.11 | 2.34 | 2.61 | 1.96 | 1.62 | 42.94 |
| BUV805 | 0.00 | 0.20 | 0.92 | 0.86 | 0.92 | 1.52 | 0.24 | 0.51 | 0.55 | 0.19 | 0.84 | 0.32 | 0.00 | 0.00 | 0.87 | 0.80 | 0.43 | 0.00 | 0.00 | 0.45 | 1.43 | 1.46 | 19.52 |
| BV421 | 0.00 | 0.53 | 0.62 | 0.00 | 0.00 | 0.27 | 0.12 | 0.23 | 0.00 | 0.00 | 0.50 | 0.00 | 0.00 | 0.00 | 0.00 | 0.00 | 0.46 | 0.00 | 0.62 | 0.14 | 0.00 | 0.00 | 33.96 |
| Super Bright 436 | 0.00 | 0.48 | 0.66 | 0.42 | 0.00 | 0.00 | 0.26 | 0.00 | 0.00 | 0.00 | 0.00 | 0.93 | 0.00 | 0.00 | 0.00 | 0.00 | 0.00 | 0.00 | 0.00 | 0.25 | 0.15 | 0.15 | 29.46 |
| eFluor 450 | 0.00 | 0.00 | 0.00 | 0.00 | 0.00 | 0.00 | 0.00 | 0.00 | 0.00 | 0.00 | 0.00 | 0.00 | 0.00 | 0.30 | 0.00 | 0.00 | 0.00 | 0.67 | 0.00 | 0.00 | 0.00 | 0.31 | 37.89 |
| BV480 | 1.02 | 0.47 | 0.50 | 0.00 | 0.48 | 0.25 | 0.87 | 1.11 | 0.60 | 0.92 | 0.58 | 0.87 | 0.00 | 0.00 | 0.13 | 0.41 | 0.00 | 0.41 | 0.00 | 0.00 | 0.35 | 0.18 | 59.13 |
| BV510 | 0.00 | 0.42 | 0.00 | 0.00 | 0.25 | 0.66 | 0.85 | 0.80 | 0.00 | 1.06 | 0.00 | 0.00 | 0.69 | 0.62 | 0.32 | 0.48 | 0.57 | 0.47 | 0.94 | 0.53 | 0.00 | 0.34 | 47.58 |
| Spark Violet 538 | 0.35 | 0.00 | 0.58 | 0.81 | 0.00 | 0.92 | 0.40 | 0.87 | 0.82 | 0.85 | 1.24 | 1.57 | 0.00 | 0.00 | 0.00 | 0.00 | 0.00 | 0.71 | 0.00 | 0.48 | 0.20 | 0.21 | 33.76 |
| BV570 | 0.65 | 1.17 | 1.08 | 0.56 | 0.47 | 0.99 | 2.02 | 2.44 | 1.84 | 2.25 | 3.37 | 3.34 | 1.16 | 0.00 | 0.67 | 0.53 | 1.26 | 0.00 | 0.00 | 0.00 | 0.00 | 0.00 | 51.27 |
| BV605 | 0.72 | 2.21 | 2.41 | 1.49 | 0.76 | 1.04 | 1.38 | 1.68 | 1.98 | 2.99 | 4.40 | 4.63 | 1.98 | 0.87 | 1.15 | 0.86 | 1.52 | 1.65 | 1.13 | 0.62 | 0.00 | 0.00 | 56.38 |
| BV650 | 0.00 | 2.01 | 2.35 | 1.62 | 0.93 | 1.22 | 0.53 | 0.15 | 1.20 | 0.83 | 4.41 | 4.75 | 1.86 | 0.66 | 0.83 | 0.75 | 2.31 | 3.65 | 2.41 | 1.00 | 0.76 | 0.58 | 57.71 |
| BV711 | 0.00 | 2.08 | 2.93 | 2.15 | 1.61 | 2.55 | 0.74 | 0.00 | 0.54 | 0.00 | 2.84 | 3.41 | 2.07 | 0.95 | 1.39 | 1.29 | 1.81 | 3.28 | 3.14 | 2.22 | 1.58 | 1.15 | 55.70 |
| BV750 | 0.00 | 0.15 | 1.34 | 1.22 | 1.79 | 3.29 | 0.50 | 0.00 | 0.00 | 0.00 | 1.05 | 0.78 | 0.61 | 0.64 | 1.14 | 1.32 | 0.00 | 0.32 | 0.00 | 1.12 | 1.29 | 0.99 | 34.28 |
| BV785 | 0.00 | 0.30 | 0.92 | 0.66 | 2.18 | 3.90 | 0.71 | 0.00 | 0.00 | 0.00 | 1.17 | 0.64 | 0.37 | 0.43 | 1.26 | 1.87 | 0.00 | 0.00 | 0.00 | 0.45 | 1.50 | 1.06 | 31.02 |
| BB515 | 4.30 | 0.57 | 0.70 | 0.49 | 0.21 | 0.00 | 2.81 | 2.35 | 1.36 | 1.36 | 0.95 | 0.87 | 0.23 | 0.18 | 0.00 | 0.00 | 0.35 | 0.69 | 0.43 | 0.00 | 0.00 | 0.12 | 36.91 |
| cFluor B532 | 4.66 | 1.09 | 1.00 | 0.28 | 0.44 | 0.19 | 3.20 | 2.74 | 1.51 | 1.61 | 1.20 | 1.41 | 0.26 | 0.00 | 0.16 | 0.16 | 0.78 | 0.46 | 0.00 | 0.00 | 0.00 | 0.29 | 36.49 |
| Spark Blue 550 | | 2.29 | 2.48 | 1.56 | 0.79 | 0.98 | 4.18 | 3.76 | 2.66 | 2.55 | 2.84 | 2.82 | 0.95 | 0.35 | 0.54 | 0.37 | 1.66 | 2.51 | 1.64 | 0.38 | 0.17 | 0.00 | 61.56 |
| PerCP | 0.86 | | 5.75 | 0.00 | 0.00 | 0.00 | 0.00 | 0.00 | 0.00 | 0.00 | 10.17 | 11.72 | 0.00 | 1.26 | 1.40 | 0.56 | 1.95 | 3.64 | 2.44 | 0.95 | 0.80 | 0.00 | 61.36 |
| PerCP/Cy5.5 | 0.00 | 8.43 | | 6.33 | 2.99 | 3.67 | 0.79 | 0.00 | 2.06 | 0.00 | 9.84 | 11.38 | 4.78 | 2.32 | 2.66 | 1.79 | 3.77 | 6.97 | 5.78 | 2.04 | 1.66 | 1.45 | 99.97 |
| PerCP-Vio700 | 0.00 | 7.75 | 10.66 | | 3.78 | 4.62 | 1.55 | 0.34 | 1.22 | 0.64 | 8.14 | 9.14 | 5.27 | 2.88 | 3.33 | 2.43 | 3.20 | 5.34 | 5.28 | 2.21 | 1.66 | 1.45 | 101.93 |
| Real Blue 780 | 0.66 | 1.08 | 1.21 | 1.24 | | 7.98 | 0.71 | 0.51 | 0.13 | 0.46 | 0.95 | 1.08 | 1.07 | 1.16 | 1.92 | 0.83 | 0.40 | 0.58 | 0.47 | 0.32 | 1.55 | 1.72 | 36.21 |
| PerCP/Fire 806 | 0.00 | 1.75 | 1.58 | 0.91 | 2.59 | | 0.30 | 0.00 | 0.00 | 0.00 | 1.55 | 2.04 | 0.95 | 0.74 | 1.90 | 2.52 | 0.68 | 0.92 | 0.00 | 0.14 | 1.04 | 1.43 | 28.35 |
| PE | 1.03 | 1.85 | 1.74 | 1.00 | 0.44 | 0.38 | | 7.02 | 2.88 | 4.18 | 3.11 | 3.11 | 1.09 | 0.41 | 0.47 | 0.30 | 1.11 | 1.64 | 1.02 | 0.28 | 0.09 | 0.19 | 46.34 |
| cFluor YG584 | 0.70 | 1.54 | 1.44 | 0.94 | 0.39 | 0.62 | 1.81 | | 1.76 | 2.98 | 3.36 | 3.65 | 1.50 | 0.52 | 0.59 | 0.41 | 0.95 | 1.47 | 0.77 | 0.15 | 0.00 | 0.00 | 31.63 |
| PE/Dazzle594 | 0.98 | 4.03 | 3.97 | 2.46 | 0.91 | 1.09 | 2.44 | 2.73 | | 5.64 | 6.98 | 7.42 | 2.44 | 1.07 | 1.32 | 0.81 | 2.25 | 3.38 | 2.13 | 0.49 | 0.28 | 0.32 | 64.92 |
| cFluor YG610 | 0.73 | 2.98 | 2.69 | 1.95 | 0.76 | 1.05 | 1.30 | 2.10 | 2.34 | | 6.55 | 6.70 | 2.43 | 0.97 | 1.21 | 0.93 | 1.15 | 1.63 | 1.32 | 0.33 | 0.14 | 0.15 | 51.97 |
| PE/Fire 640 | 0.66 | 6.25 | 6.45 | 4.34 | 1.85 | 2.24 | 1.59 | 1.64 | 2.77 | 2.74 | | 10.25 | 4.59 | 2.03 | 2.30 | 1.62 | 4.33 | 6.68 | 4.30 | 1.26 | 0.80 | 0.67 | 81.37 |
| PE/Cy5 | 0.58 | 8.15 | 7.34 | 4.53 | 1.79 | 2.04 | 1.36 | 1.25 | 2.71 | 1.91 | 15.16 | | 4.80 | 2.21 | 2.32 | 1.51 | 4.81 | 9.38 | 5.21 | 1.39 | 0.87 | 0.65 | 92.04 |
| cFluor BYG710 | 0.46 | 5.60 | 6.83 | 5.56 | 2.36 | 2.47 | 2.55 | 2.16 | 1.73 | 1.59 | 5.96 | 6.78 | | 2.45 | 2.84 | 1.85 | 1.81 | 3.11 | 3.18 | 1.25 | 0.83 | 0.64 | 72.51 |
| cFluor BYG750 | 0.35 | 3.64 | 4.89 | 4.12 | 3.67 | 4.62 | 2.93 | 2.35 | 1.44 | 1.72 | 3.87 | 4.46 | 4.15 | | 4.91 | 3.76 | 1.17 | 1.66 | 1.84 | 1.01 | 1.35 | 1.19 | 70.29 |
| PE/Cy7 | 0.30 | 1.26 | 1.68 | 1.27 | 4.67 | 5.40 | 1.52 | 0.48 | 0.40 | 0.44 | 1.14 | 1.25 | 1.40 | 2.16 | | 4.84 | 0.33 | 0.39 | 0.66 | 0.26 | 1.76 | 1.24 | 39.30 |
| PE/Fire 810 | 0.00 | 1.62 | 1.80 | 1.19 | 2.57 | 6.20 | 3.65 | 2.63 | 1.49 | 1.80 | 2.31 | 2.27 | 1.46 | 1.09 | 3.73 | | 0.55 | 1.18 | 0.81 | 0.18 | 0.88 | 1.31 | 47.10 |
| APC | 0.00 | 2.65 | 2.50 | 1.58 | 0.76 | 0.92 | 0.55 | 0.87 | 1.51 | 1.35 | 5.71 | 6.55 | 2.31 | 1.12 | 1.36 | 0.94 | | 7.95 | 5.16 | 1.86 | 1.50 | 1.08 | 55.44 |
| Alexa Fluor 647 | 0.00 | 1.72 | 1.45 | 1.16 | 0.30 | 0.00 | 0.00 | 0.00 | 1.00 | 0.36 | 4.54 | 5.12 | 1.53 | 0.67 | 0.85 | 0.56 | 4.82 | | 5.92 | 1.87 | 1.63 | 1.12 | 38.07 |
| Spark NIR 685 | 0.00 | 2.03 | 1.86 | 1.41 | 0.65 | 0.79 | 0.12 | 0.23 | 0.78 | 0.48 | 4.00 | 4.46 | 1.99 | 0.76 | 1.06 | 0.72 | 4.34 | 8.55 | | 2.14 | 2.08 | 1.33 | 43.62 |
| cFluor R720 | 0.00 | 1.41 | 1.50 | 1.04 | 0.90 | 1.24 | 0.44 | 0.00 | 0.00 | 0.00 | 1.17 | 1.75 | 1.94 | 0.76 | 1.34 | 1.00 | 2.21 | 4.53 | 4.20 | | 2.97 | 2.10 | 35.89 |
| APC-H7 | 0.00 | 0.56 | 0.73 | 0.14 | 1.52 | 2.15 | 0.93 | 0.24 | 0.00 | 0.00 | 0.80 | 0.00 | 0.58 | 0.81 | 2.54 | 2.00 | 0.57 | 0.98 | 0.93 | 0.45 | | 3.20 | 22.54 |
| APC/Fire 810 | 0.00 | 0.82 | 0.80 | 0.37 | 1.31 | 2.67 | 0.32 | 0.00 | 0.00 | 0.00 | 1.33 | 1.62 | 0.69 | 0.40 | 1.61 | 2.52 | 1.65 | 2.12 | 1.51 | 0.60 | 2.58 | | 28.45 |
| Sum | 21.85 | 84.69 | 93.92 | 61.89 | 47.55 | 69.94 | 49.03 | 46.65 | 43.48 | 51.38 | 136.41 | 142.48 | 61.82 | 33.88 | 51.35 | 41.60 | 62.30 | 99.95 | 73.29 | 31.52 | 33.19 | 31.14 | 4187.77 |

Spillover Spreading Matrix Color Coding: 0 3 9

**4E**

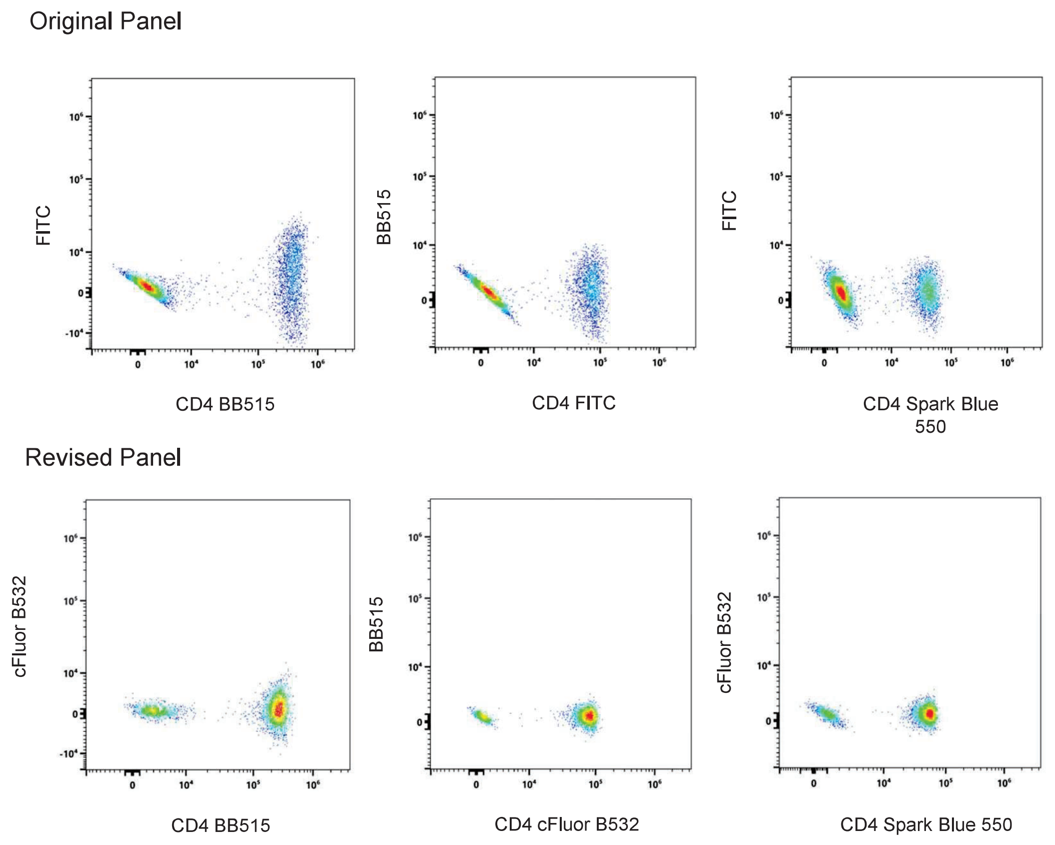

**Online Figure 4. Characterization of Performance of 45-color OMIP**

**(A)** Spectral signatures of the 45 fluorochromes used in this OMIP were overlaid using the Cytek Cloud. All signatures are normalized to peak channel for direct comparison. Fluorochromes that are different to the ones used in OMIP-069 are highlighted in green. **(B)** Similarity index matrix. The chart displays the numerical value for each pair of fluorochromes selected for the 45-color assay. The highest similarity index value is 0.95. On the right side of the matrix, the complexity index, is displayed. **(C)** Stain indices calculated for each of the fluorochromes in the panel, ranked from low to high. The colors in the bars represent the primary excitation laser line: 355 nm (grey), 405 nm (violet), 488 nm (blue), 561 nm (green) and 635 nm (red). Arrows indicate the new fluorochromes that are different or additional to the ones in OMIP-069**.** **(D)** The spillover-spread of the 45-fluorochome combination. The spillover spread matrix (SSM) was calculated using CD4 stained cells and the viability dye. Spillover values are color-coded as follows: white: <3, shades of orange: 3-9, and red: >9. **(E)** Comparison of the spread between BB515 and FITC or BB515 and cFluor B532 using CD4 stained cells. Differences in the spread from Spark Blue 550 to FITC or cFluor are also shown.

Based on brightness, the use of these fluorochrome combinations on another spectral cytometer would require characterization of that instrument and the generation of a spillover spread matrix for that particular instrument, as multiple instrument design factors can impact this measurement.

#### 3. Panel design strategy

As mentioned, OMIP-069 was used as a starting point from the biology/marker selection perspective and for the fluorochrome choices.

In terms of biology, to expand the characterization of different T cell subsets, thirty-seven markers were kept from the original panel and 8 new markers were added. The new markers added were: CD11b, TIM-3, DNAM-1, CD103, TIGIT, CD161, GPR56, and KLRG1. The three markers removed from OMIP-069 were used for subsetting NK and dendritic cells (DC) populations and were CD159a and CD159c (NK cells) and CD141 (DCs).

The strategy for fluorochrome assignment was as follows:

1. Original markers in OMIP-069 needing fluorochrome replacement: As explained in section 1, seven (7) fluorochrome replacements to the original fluorochromes used in OMIP-069 were identified based on performance or reagent availability. For six reagents, we identified a direct replacement by having the same marker conjugated to a fluorochrome with similar or superior performance. Those replacements were: CD20 Spark Violet 538, CD127 cFluor R720, CD25 cFluor BYG710, TCRγδ PerCP-Vio 700, CD24 cFluor YG610, and CD57 cFluor B532. The same clone was available for CD24 (SN3) and CD57 (HNK-1), whereas for CD20,

CD25, CD127, and TCRγδ we had to identify clones with similar performance **(Online Figure 1)**. A direct comparison of the newly added markers, comparing new fluorochromes against PE, a known bright well characterized fluorochrome, is shown in **Online Figure 5.**

2. For those users having issues with HLA-DR PE/Fire 810 stability and wishing to run OMIP-069, we decided to move HLA-DR to BV480 and to assign IgD to cFluor BYG750. This swap was made based on antigen level of expression, fluorochrome brightness and co-expression with markers already assigned to fluorochromes having spillover spread from or into BV480 and cFluor BYG750.
3. Of the newly identified markers, CD11b was the only available in Spark UV387.
4. For all the remaining new markers (TIM-3, DNAM-1, CD103, TIGIT, CD161, and KLRG1) clones were selected based on internal historical data, literature, reagent availability, and information available from reagent manufacturers. For the new panel to be easily adopted, it was critical that reagents were commercially available. Of importance, all these markers were classified as tertiary and therefore needed to be assigned to bright fluorochromes. A list of the reagents tested throughout the panel's development is in **Online Table 2**. Our strategy to qualify reagents to be incorporated in the panel followed the following stepwise procedure. First, a full antibody titration and comparison at optimal titer to a reference reagent (often conjugated to PE or other similarly bright fluorochrome) was done. If significant differences were observed in resolution or non-specific binding, the reagent under testing was not considered for further evaluation. For those reagents showing adequate performance based on this first step, testing using the full panel, or a subset of markers was performed (**Online Figure 3C**). After this careful evaluation, the following reagents were selected: TIM-3 BB515, DNAM-1 Real Blue 780, GPR56 PE, TIGIT PE/Fire 810, CD161 APC, and KLRG1 PE/Fire 810.

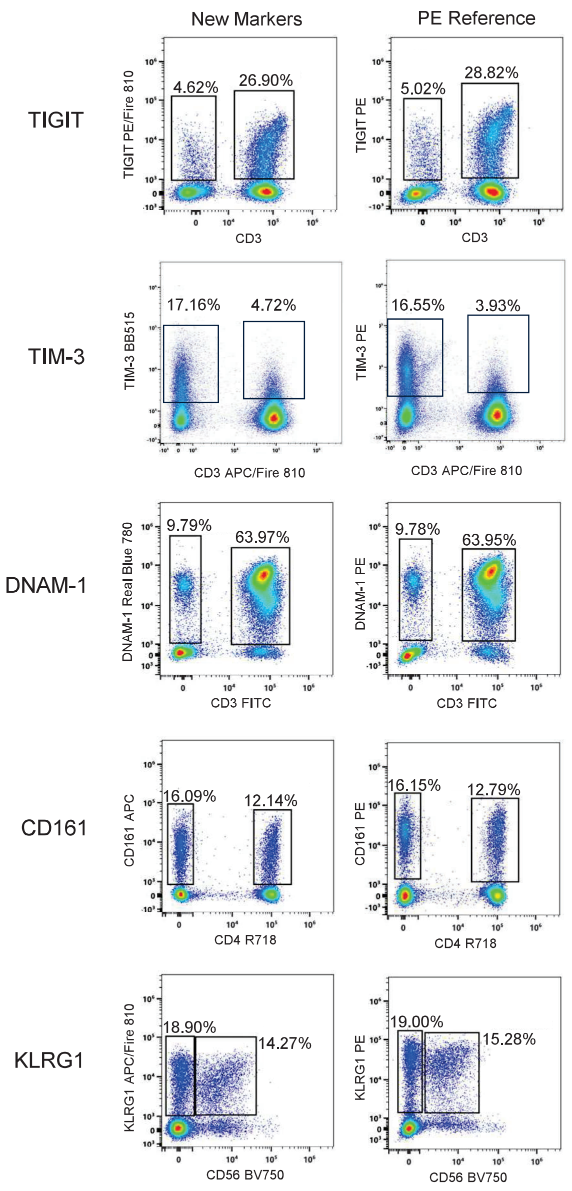

**Online Figure 5. Evaluation of New Reagent Performance.** Identification of optimal reagents for the panel was done by comparison of the same clone conjugated to the assigned fluorochrome vs a PE conjugate. Titrations were done in parallel for both reagents. Evaluation of TIGIT PE/Fire 810, TIM-3 BB515, DNAM-1 Real Blue 780, CD161 APC and KRLG-1 APC/Fire 810 are presented. TIGIT, TIM-3 and DNAM-1 had equivalent performance compared to their PE counterpart at the optimal titer. CD161 APC and KLRG1 APC/Fire 810 positive populations were dimmer than when using PE conjugates, but still resolved. The percentage of positive cells was preserved for all comparisons. As shown in the figure, co staining with CD3, CD4, or CD56 was used to facilitate identification of the populations of interest.

5. Of the markers needing fluorochrome assignment, none worked adequately in PerCP/Fire 806 (**Online Table 2 and Online Figure 3C**), so we chose to move CD4 to PerCP/Fire 806. cFluor YG584 became available and based on level of expression, co-expression, and reagent availability, we assigned it to CD56. BUV737 became available and was assigned to CD38, freeing up APC/Fire 810 used for KLRG1.
6. Based on the results of this testing, panel design was finalized. New assignments outlined in green are shown in **Online Figure 6** and the full list of reagents is presented in **Online Table 3**, with new conjugates highlighted in green.

### 4. 45-color assay optimization

#### a. Antibody titrations

Titrations were done for the nineteen reagents not part of the original OMIP-069 panel and eleven for those reagents changed in titers compared to OMIP-069. Titrations were done using previously frozen PBMCs, $2x10^5$ cells were used per titration point and True-Stain Monocyte Blocker™ (BioLegend, catalog #426101) was added to the cells at a dilution of five µl per 100 µl of cell suspension, per the manufacturer recommendation. All titrations we made in 96-well plates. The cells were resuspended in a final volume of 250 µL to mimic the final staining volume of the multicolor sample. Of note, after full optimization the final volume of our cell suspension was 292 µL. Because of the volume being close to the one used for the initial titrations, we did not consider additional titrations were needed. The concentrations tested (expressed in ng/test, where a test ranged from 50 to 500 µL of cell suspension) varied across reagents. Moreover, to evaluate non-specific binding and the titration in specific cellular subsets, in some of the titrations an additional marker was included: CD19 for CD24 and IgD; CD3 for TIM-3, DNAM-1, and CD103; CD56 for KLRG1 and GPR56; and CD4 for CD161 and CD25. Titration was evaluated using concatenated

| UV 355 | | V 405 | | B 488 | | YG 561 | | R 640 | |
|---|---|---|---|---|---|---|---|---|---|
| Marker | Fluor | Marker | Fluor | Marker | Fluor | Marker | Fluor | Marker | Fluor |
| CD11b | Spark UV 387 | | | | | | | | |
| CD45RA | BUV395 | | | | | | | | |
| | | CD197 (CCR7) | BV421 | | | | | | |
| | | CD123 | Super Bright 436 | | | | | | |
| | | CD11c | eFluor 450 | | | | | | |
| Viability | LIVE DEAD Blue | | | | | | | | |
| CD16 | BUV496 | MHC Class II (HLA-DR) | BV480 | CD366 (TIM3) | BB515 | | | | |
| | | | | CD57 | cFluor B532 | | | | |
| | | CD20 | Spark Violet 538 | CD14 | Spark Blue 550 | | | | |
| | | CD3 | BV510 | | | | | | |
| CD195 (CCR5) | BUV563 | IgM | BV570 | | | CD56 | cFluor YG584 | | |
| | | | | | | GPR56 | PE | | |
| CD314 | BUV615 | IgG | BV605 | | | CD24 | cFluor YG610 | | |
| | | | | | | CD337 | PE-Dazzle 594 | | |
| CD39 | BUV661 | CD28 | BV650 | | | CD103 | PE-Fire 640 | CD161 | APC |
| | | | | CD45 | PerCP | CD95 (FAS) | PE-Cy5 | CD1C | Alexa Fluor 647 |
| | | | | CD2 | PerCP-Cy5.5 | | | CD19 | Spark NIR 685 |
| | | CD196 (CCR6) | BV711 | TCR gd | PerCP-Vio 700 | CD25 | cFluor BYG710 | CD127 | cFluor R720 |
| CD38 | BUV737 | CD185 (CXCR5) | BV750 | | | IgD | cFluor BYG750 | | |
| | | CD279 (PD-1) | BV785 | | | CD183 (CXCR3) | PE-Cy7 | CD27 | APC-H7 |
| CD8 | BUV805 | | | CD226 | RB780 | TIGIT | PE-Fire 810 | KLRG1 | APC-Fire 810 |
| | | | | CD4 | PerCP-Fire 806 | | | | |

**Online Figure 6. Panel Design**

The final panel design is shown. We organized the reagents according to their primary excitation wavelength (columns) and peak emission wavelength (rows). Markers (8) and fluorochromes (11) that were not present in OMIP-69 are highlighted in green boxes. The panel optical layout was built using the Cytek Cloud. Fluorochrome brightness, compared to PE, is displayed in colored bars under the fluorochrome names. Brightness bar color indicates primary exciting laser (dark purple-UV, light purple-V, blue-B, green-YG, and red-R). The blue bars displayed under the marker names, show relative antigen density, defined manually in the Cytek Cloud.

| Specificity | Fluorochrome | Clone | Catalog # | Vendor | Titer (ng/300 µL) |
|---|---|---|---|---|---|
| Viability | Live/Dead Blue | - | L34962 | Thermo Fisher | 10 µL of 1:40 |
| CD45 | PerCP | 2D1 | 368505 | BioLegend | 250 |
| CD3 | BV510 | SK7 | 344827 | BioLegend | 400 |
| **CD4** | **PerCP/Fire 806** | **SK3** | **344693** | **BioLegend** | **100** |
| CD8 | BUV805 | SK1 | 612889 | BD Biosciences | 250 |
| **CD25** | **cFluor BYG710** | **4E3** | **R7-20660** | **Cytek Biosciences** | **500** |
| PD-1 | BV785 | EH12.2H7 | 329929 | BioLegend | 250 |
| **TCRγδ** | **PerCP-Vio700** | **REA591** | **130-114-040** | **Miltenyi** | **800** |
| CD14 | Spark Blue 550 | 63D3 | 367147 | BioLegend | 1000 |
| CD16 | BUV496 | 3G8 | 612945 | BD Biosciences | 250 |
| **CD11b** | **Spark UV 387** | **ICRF44** | **301365** | **BioLegend** | **1000** |
| CD11c | eFluor 450 | 3.9 | 48-0116-41 | | 1000 |
| CD19 | Spark NIR 685 | HIB19 | 302269 | BioLegend | 125 |
| **CD20** | **Spark Violet 538** | **2H7** | **302373** | **Biolegend** | **500** |
| **CD24** | **cFluor YG610** | **SN3** | **R7-20658** | **Cytek Biosciences** | **1000** |
| CD39 | BUV661 | TU66 | 569788 | BD Biosciences | 1000 |
| **IgD** | **cFluor BYG750** | **IgD26** | **R7-20662** | **Cytek Biosciences** | **32** |
| IgG | BV605 | G18-145 | 563246 | BD Biosciences | 250 |
| IgM | BV570 | MHM-88 | 314517 | BioLegend | 250 |
| CD1c | Alexa Fluor 647 | L161 | 331510 | BioLegend | 500 |
| CD123 | Super Bright 436 | 6H6 | 62-1239-42 | Thermo Fisher | 125 |
| CD2 | PerCP/Cy5.5 | TS1/8 | 309225 | BioLegend | 250 |
| **CD56** | **cFluor YG584** | **5.1H11** | **R7-20786** | **Cytek Biosciences** | **500** |
| CCR7 | BV421 | G043H7 | 353207 | BioLegend | 700 |
| CD27 | APC-H7 | M-T271 | 560223 | BD Biosciences | 250 |
| CD45RA | BUV395 | 5H9 | 569489 | BD Biosciences | 250 |
| CD95 | PE/Cy5 | DX2 | 305610 | BioLegend | 125 |
| **CD127** | **cFluor R720** | **A019D5** | **R7-20664** | **Cytek Biosciences** | **500** |
| CD337 | PE/Dazzle 594 | P30-15 | 325231 | BioLegend | 250 |
| CCR6 | BV711 | G034E3 | 353435 | BioLegend | 500 |
| CCR5 | BUV563 | 2D7/CCR5 | 741401 | BD Biosciences | 500 |
| CXCR5 | BV750 | RF8B2 | 569501 | BD Biosciences | 500 |
| CXCR3 | PE/Cy7 | G025H7 | 353719 | BioLegend | 1000 |
| **HLA-DR** | **BV480** | **L203** | **752499** | **BD Biosciences** | **250** |
| **CD38** | **BUV737** | **HB7** | **612825** | **BD Biosciences** | **500** |
| **CD57** | **cFluor B532** | **HNK-1** | **R7-20656** | **Cytek Biosciences** | **8** |
| CD28 | BV650 | CD28.2 | 302945 | BioLegend | 500 |
| CD314 | BUV615 | 1D11 | 751232 | BD Biosciences | 1000 |
| **TIM-3** | **BB515** | **7D3** | **565569** | **BD Biosciences** | **250** |
| **TIGIT** | **PE/Fire 810** | **A15153G** | **372745** | **BioLegend** | **500** |
| **GPR56** | **PE** | **CG4** | **358203** | **BioLegend** | **125** |
| **DNAM-1** | **Real Blue 780** | **DX11** | **755549** | **BD Biosciences** | **1000** |
| **CD103** | **PE/Fire 640** | **Ber-ACT8** | **350243** | **BioLegend** | **250** |
| **CD161** | **APC** | **HP-3G10** | **17-1619-42** | **Thermo Fisher** | **500** |
| **KLRG1** | **APC/Fire 810** | **SA231A2** | **367731** | **BioLegend** | **500** |

**Online Table 3. Final Selection of Reagents Used in OMIP-xxx**
List of reagents used in the final panel with antibody clone, manufacturer information, and final concentrations used in the staining protocol provided. Reagents that are different or additions to the ones in OMIP-069 are highlighted in bold/green.

files, calculating stain index, evaluating binding to monocytes, spread and brightness of the negative population, and percent positive cells to determine the optimal titer. In addition, each marker was plotted vs SSC to evaluate staining pattern and non-specific binding. Results of the titration experiments are presented in **Online Figure 7A and B.** Moreover, reagents used in OMIP-069, for which the titer changed after full panel optimization are presented in **Online Figure 7C**. The final titers for all reagents, expressed in ng/total staining volume (300µL), are summarized in **Online Table 3.**

#### b. Reference control optimization

To accurately assess multicolor panel performance, it is necessary to define the optimal reference controls that will lead to accurate unmixing. Optimization of the controls entails establishing if beads or cells lead to accurate unmixing for each fluorochrome, as differences in the spectrum of conjugated antibodies might differ when binding cells vs. beads (1). For this purpose, beads (UltraComp eBeads Plus Compensation Beads: Thermo Fisher, catalog #01-3333-41) and cells were stained in parallel with the 44 fluorochromes selected for this panel (for the viability dye, cells were used as control). Unmixing accuracy was evaluated by statistically assessing the medians of the negative and positive populations of the overlapping channel in single stained cells unmixed with beads as controls. Data were unmixed using the Ordinary Least Squares algorithm available in SpectroFlo® software (Cytek Biosciences, Fremont, CA). For fifteen (15) fluorochromes, we found that beads led to unmixing inaccuracies and hence cells were used. The fifteen fluorochromes that required cell reference controls were: BUV496, BUV563, BUV661, BUV737, BV480, BV570, BV750, Spark Blue 550, PerCP-Cy5.5, Real Blue 780, PE, cFluor YG584, PE/Fire 640, cFluor BYG750, and LIVE/DEAD Blue Fixable Viability Dye **(Online Table 4 and Figure 8)**. These findings agree with the recommendation provided in OMIP-069, with some exceptions. For BUV805, PE/Cy5, APC-H7, cells are no longer mandatory, whereas  for BUV737, BV750, PerCP/Cy5.5, PE, cFluor YG584

7A

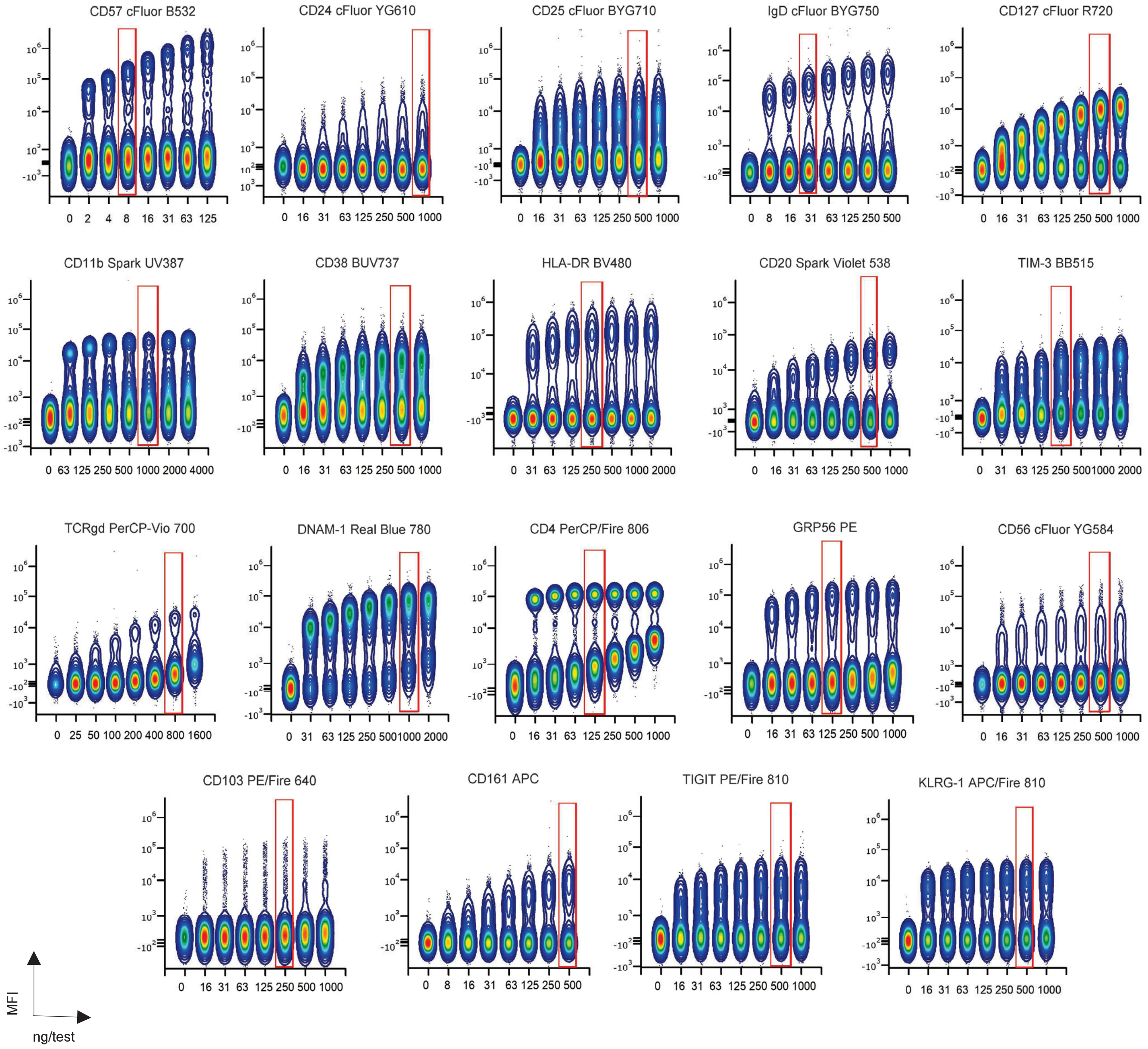
CD57 cFluor B532
CD24 cFluor YG610
CD25 cFluor BYG710
IgD cFluor BYG750
CD127 cFluor R720
CD11b Spark UV387
CD38 BUV737
HLA-DR BV480
CD20 Spark Violet 538
TIM-3 BB515
TCRgd PerCP-Vio 700
DNAM-1 Real Blue 780
CD4 PerCP/Fire 806
GRP56 PE
CD56 cFluor YG584
CD103 PE/Fire 640
CD161 APC
TIGIT PE/Fire 810
KLRG-1 APC/Fire 810
MFI
ng/test

**7B**

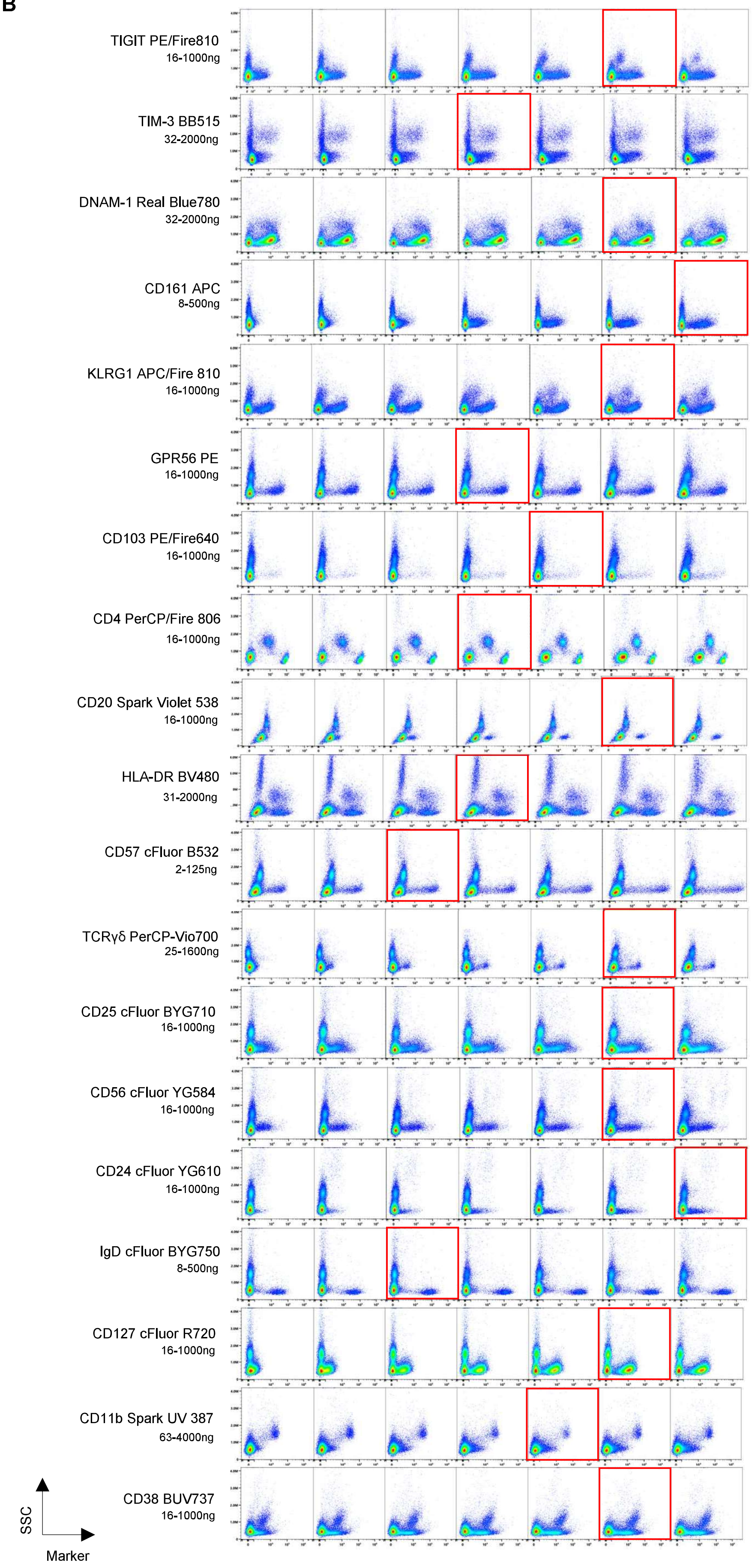
TIGIT PE/Fire810
16-1000ng
TIM-3 BB515
32-2000ng
DNAM-1 Real Blue780
32-2000ng
CD161 APC
8-500ng
KLRG1 APC/Fire 810
16-1000ng
GPR56 PE
16-1000ng
CD103 PE/Fire640
16-1000ng
CD4 PerCP/Fire 806
16-1000ng
CD20 Spark Violet 538
16-1000ng
HLA-DR BV480
31-2000ng
CD57 cFluor B532
2-125ng
TCRγδ PerCP-Vio700
25-1600ng
CD25 cFluor BYG710
16-1000ng
CD56 cFluor YG584
16-1000ng
CD24 cFluor YG610
16-1000ng
IgD cFluor BYG750
8-500ng
CD127 cFluor R720
16-1000ng
CD11b Spark UV 387
63-4000ng
CD38 BUV737
16-1000ng
SSC
Marker

**7C**

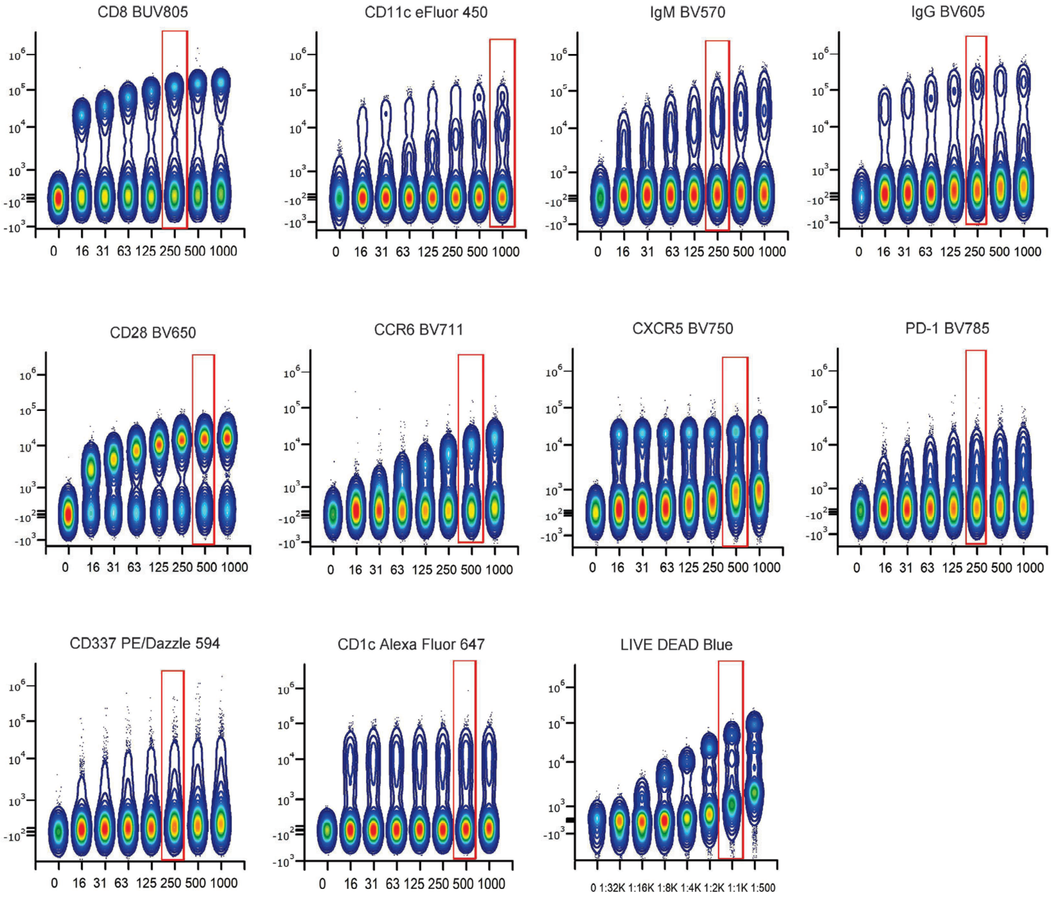

**Online Figure 7. Antibody Titrations**

Optimal concentrations of all antibodies were determined through titration experiments. Two-fold dilutions of antibodies were tested, with concentrations from lowest to highest, depending on the marker. The highest concentration tested for each antibody was 2-fold the one recommended by the manufacturer and from there 7 sequential dilutions were made. **(A)** Titrations for the new reagents are shown. Files were concatenated for analysis using FCS Express version 7 (De Novo Software). Optimal titer is shown in red box. **(B)** Non-specific binding and staining pattern were evaluated by plotting each marker (x-axis) vs. SSC. The range of concentrations tested is specified underneath the reagent name. Final titration results are expressed as ng/test. **(C)** Titrations for reagents whose titer changed, compared to the concentrations used for OMIP-69. Final titration results are expressed as ng per staining volume, and in dilution for LIVE/ DEAD Blue in final staining volume of 250 µL. (see **Online Table 3**).

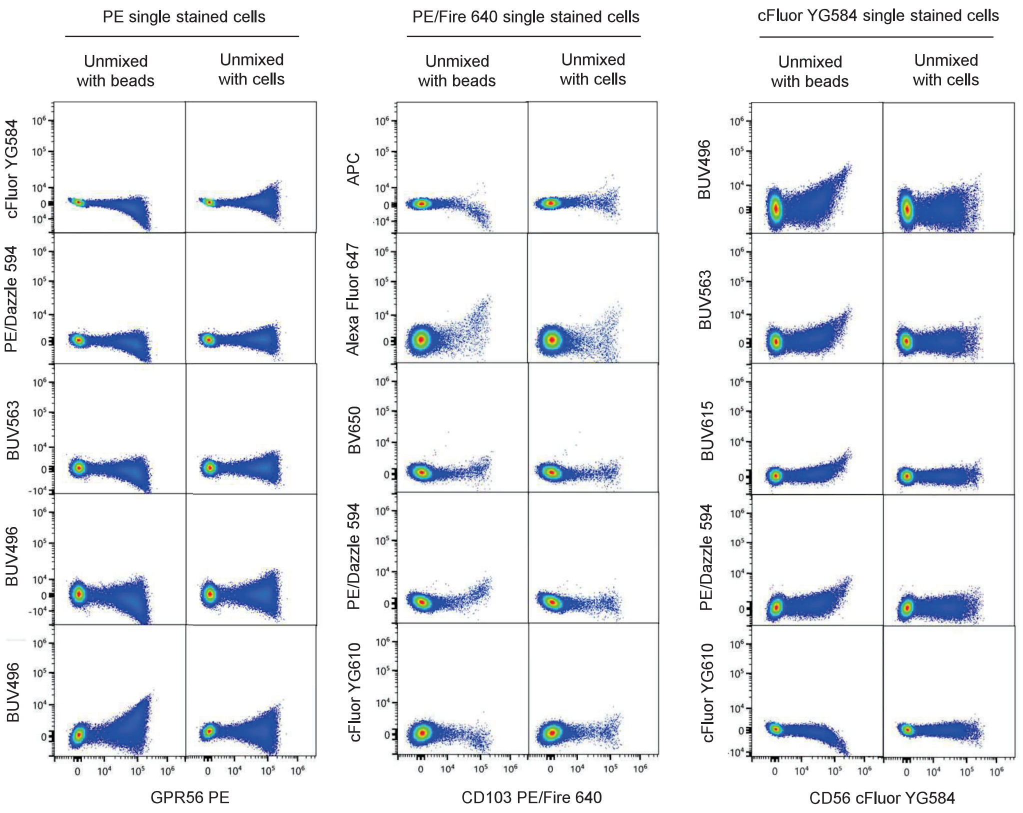

**Online Figure 8. Unmixing accuracy evaluation**

When evaluating unmixing outcome using beads vs. cells as reference controls, several fluorochromes were identified for which the use of beads led to unmixing inaccuracies. Examples of unmixing errors are shown for GPR56 PE, CD103 PE/Fire640, and CD56 cFluor YG584 conjugates bound to beads or cells, and then used to unmix single stained cells. Unmixing inaccuracies were observed as populations tilted in the X-axis. OMIP-069 (1) showed how these unmixing inaccuracies were the result of minor differences in the spectra of fluorochromes when the same antibody was bound to beads vs. cells.

| Antibody | Reference control recommendation | Number of lymphocytes to acquire | Sequential staining step |
|---|---|---|---|
| CD11b Spark UV387 | Cells or Beads | 20,000 | 6 |
| CD45RA BUV395 | Cells or Beads | 10,000 | 6 |
| CD16 BUV496 | Cells | 20,000 | 6 |
| CCR5 BUV563 | Cells | 40,000 | 2 |
| CD314 BUV615 | Cells or Beads | 40,000 | 6 |
| CD39 BUV661 | Cells | 60,000 | 6 |
| CD38 BUV737 | Cells | 40,000 | 6 |
| CD8 BUV805 | Cells or Beads | 10,000 | 6 |
| CCR7 BV421 | Cells or Beads | 20,000 | 5 |
| CD123 Super Bright 436 | Cells or Beads | 60,000 | 3 |
| CD11c eFluor 450 | Cells or Beads | 20,000 | 5 |
| HLA-DR BV480 | Cells | 60,000 | 6 |
| CD3 BV510 | Cells or Beads | 10,000 | 6 |
| CD20 SparkViolet 538 | Cells or Beads | 20,000 | 6 |
| IgM BV570 | Cells | 40,000 | 6 |
| IgG BV605 | Cells or Beads | 60,000 | 3 |
| CD28 BV650 | Cells or Beads | 20,000 | 6 |
| CCR6 BV711 | Cells or Beads | 20,000 | 6 |
| CXCR5 BV750 | Cells | 40,000 | 5 |
| PD-1 BV785 | Cells or Beads | 40,000 | 5 |
| TIM-3 BB515 | Cells or Beads | 80,000 | 6 |
| CD57 cFluor B532 | Cells or Beads | 10,000 | 6 |
| CD14 Spark Blue 550 | Cells | 20,000 | 6 |
| CD45 PerCP | Cells or Beads | 10,000 | 6 |
| CD2 PerCP/Cy5.5 | Cells | 20,000 | 6 |
| TCRgd PerCP-Vio 700 | Cells or Beads | 80,000 | 4 |
| DNAM-1 Real Blue 780 | Cells | 40,000 | 6 |
| CD4 PerCP/Fire 806 | Cells or Beads | 10,000 | 6 |
| GPR56 PE | Cells | 40,000 | 6 |
| CD56 cFluor YG584 | Cells | 60,000 | 6 |
| CD337 PE/Dazzle 594 | Cells or Beads | 40,000 | 6 |
| CD24 cFluor YG610 | Cells or Beads | 20,000 | 6 |
| CD103 PE/Fire 640 | Cells | 80,000 | 6 |
| CD95 PE/Cy5 | Cells or Beads | 10,000 | 6 |
| CD25 cFluor BYG710 | Cells or Beads | 20,000 | 6 |
| IgD cFluor BYG750 | Cells | 40,000 | 6 |
| CXCR3 PE/Cy7 | Cells or Beads | 40,000 | 5 |
| TIGIT PE/Fire 810 | Cells or Beads | 40,000 | 6 |
| CD161 APC | Cells or Beads | 40,000 | 6 |
| CD1c Alexa Fluor 647 | Cells or Beads | 40,000 | 6 |
| CD19 Spark NIR 685 | Cells or Beads | 10,000 | 6 |
| CD127 cFluor R720 | Cells or Beads | 10,000 | 6 |
| CD27 APC-H7 | Cells or Beads | 10,000 | 6 |
| KLRG1 APC/Fire810 | Cells or Beads | 40,000 | 6 |
| LIVE/DEAD Blue Fixable Viability Dye | Cells | 40,000 | 1 |

**Online Table 4. Reference Control Selection.**
The unmixing accuracy outcome was evaluated when using beads vs. cells as reference controls. For each marker, recommendations on optimal particle to be used, as well as number of events to collect, if using cells, is listed. For each antibody, the order of addition is listed in the last column, according to the steps described in the staining protocol (see section 6).

cells are needed. For accurate unmixing with cell reference controls, we established the optimal number of events to acquire for each marker. This information is summarized in **Online Table 4.**

**c. Evaluation of unmixing accuracy in multicolor samples**

To check the unmixing accuracy of the multicolor samples throughout the development of the panel, NxN plot permutations for all fluorochromes, gated on singlets/non-red blood cells (RBCs)/live events were used. When unmixing errors were observed, the magnitude of the error was assessed, and no corrections were made if the permutation where the error was minimal and/or without spillover between the fluorochrome pair. For pairs which needed any correction, we evaluated if adjustments to the number of events collected or the gating position for the positive events made an improvement **(Online Figure 9)**. Corrections were made primarily for aesthetic visualization of populations and did not impact data interpretation, as confirmed by both manual and unsupervised analysis. These corrections did not affect any of the frequencies or clustering but only served to improve the visualization of the data and minimize the need for gate adjustments **(Online Figure 23)**.

**d. Individual marker resolution assessment in single stained vs. multicolor samples**

To assess the resolution of each individual marker in the multicolor sample, we compared it to the resolution in the corresponding single stained controls. This comparison determines: 1) if there is loss of resolution due to a loss in the signal in the multicolor tube, and 2) if there is loss of resolution due to spillover spread affecting the negative population. An initial assessment, following the protocol published in OMIP-069, showed loss of signal for IgG, CD123, CXCR3, CD11c, PD-1, CD161, and CCR7. Poor resolution of those markers was not caused by spillover spreading, confirming appropriate panel design. Modifications to the staining protocol were tested to improve resolution (see Step e and f).

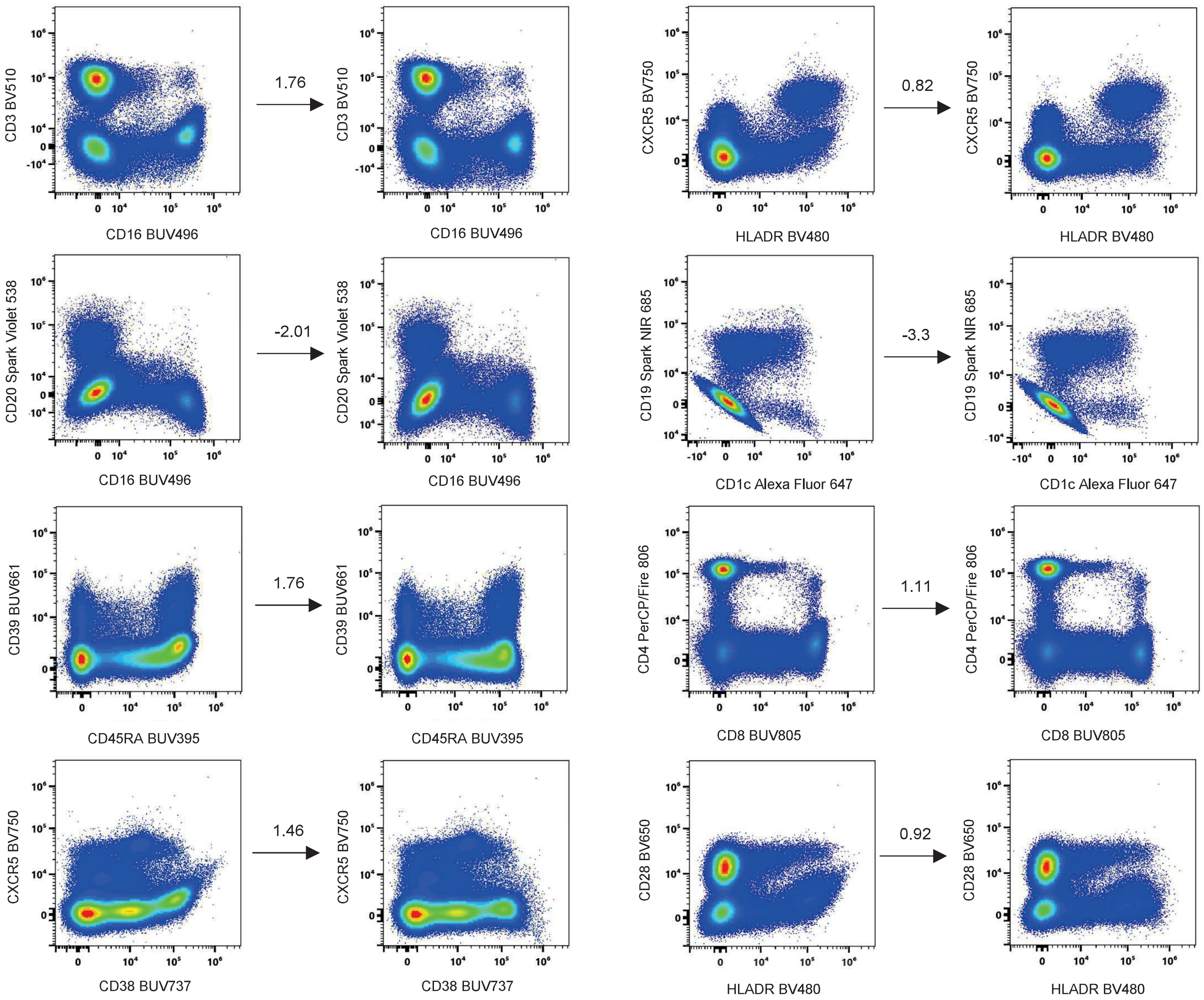

**Online Figure 9. Unmixing accuracy evaluation multicolor samples**

For multicolor samples, NxN plots were used to visually assess unmixing accuracy, gating on singlets, scatter, non-RBCs, live, and non-aggregate events. Examples of unmixing errors are shown. The magnitude of the spillover correction is shown for 8 pairs, as well as the plots before and after manual adjustments. No changes in biological patterns were observed as a result of the manual spillover correction.

#### e. Optimization of the staining protocol

For the markers which exhibited loss of signal in the multicolor tube, only CD161 could be improved by increasing the titer 2-fold. The next step was to determine if the order of addition could improve resolution. Based on OMIP-069, TCR $\gamma\delta$, CXCR5, and CCR5 need to be added sequentially for optimal staining. In this new panel, CD123 and IgG showed significant loss of signal when added in the main cocktail, while resolution for CCR7, CXCR3, PD-1, and CD11c were impacted less dramatically. Since a titer increase did not restore population resolution, sequential staining was tested to define the order of addition of all reagents. The newly introduced TCR$\gamma\delta$ PerCP-Vio 700 antibody is a human recombinant immunoglobulin, which could be related to the loss of IgG signal.

Fluorescence minus one controls (FMO)s were used to understand which antibodies could be blocking binding and the order of conjugate addition. For each FMO, the MFI and % parent of all markers present in the staining were analyzed. For example, considering markers A and B, if in the absence of marker A, marker B's brightness and % positive is higher, then marker B needs to be added before marker A. Using this method, we determined six consecutive steps: first viability then CCR5, followed by CD123 and IgG, then TCR$\gamma\delta$, next CXCR3, CCR7, CXCR5, PD-1 and CD11c, and finally a cocktail of the remaining 35 antibodies. Examples of the impact of sequential staining are shown in **Online Figure 10A**. The order of addition of each antibody is specified in **Online Table 4.** Of note, since there are differences in total volume of the sample throughout the sequential addition of the reagents, we investigated the impact of those differences according to the volume used for the titrations (300 µL). We found that the volume impacted the resolution for those reagents added in sequential step, being higher at the lower volume, but that the final titer did not change **(Online Figure 10B)**. Upon the modifications to titers and the order of addition of the reagents, the resolution of each marker was re-evaluated, by comparing single stained vs multicolor samples **(Online**

**10A**

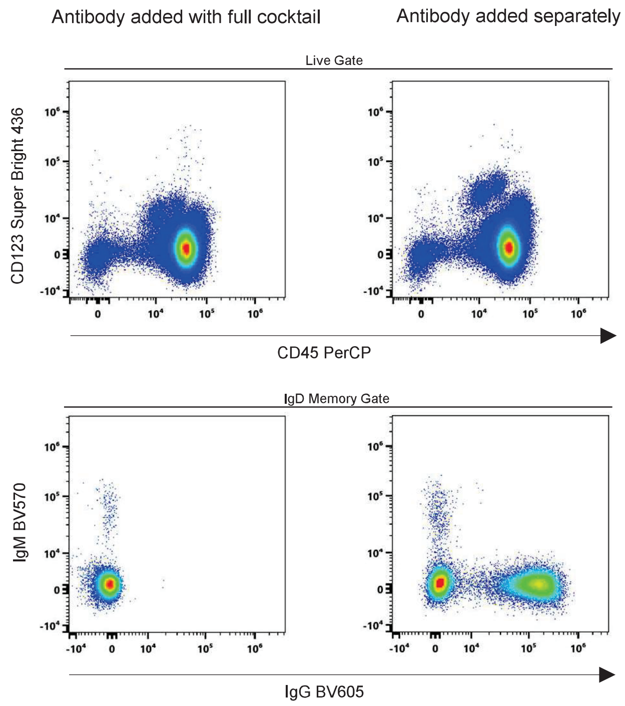

**10B**

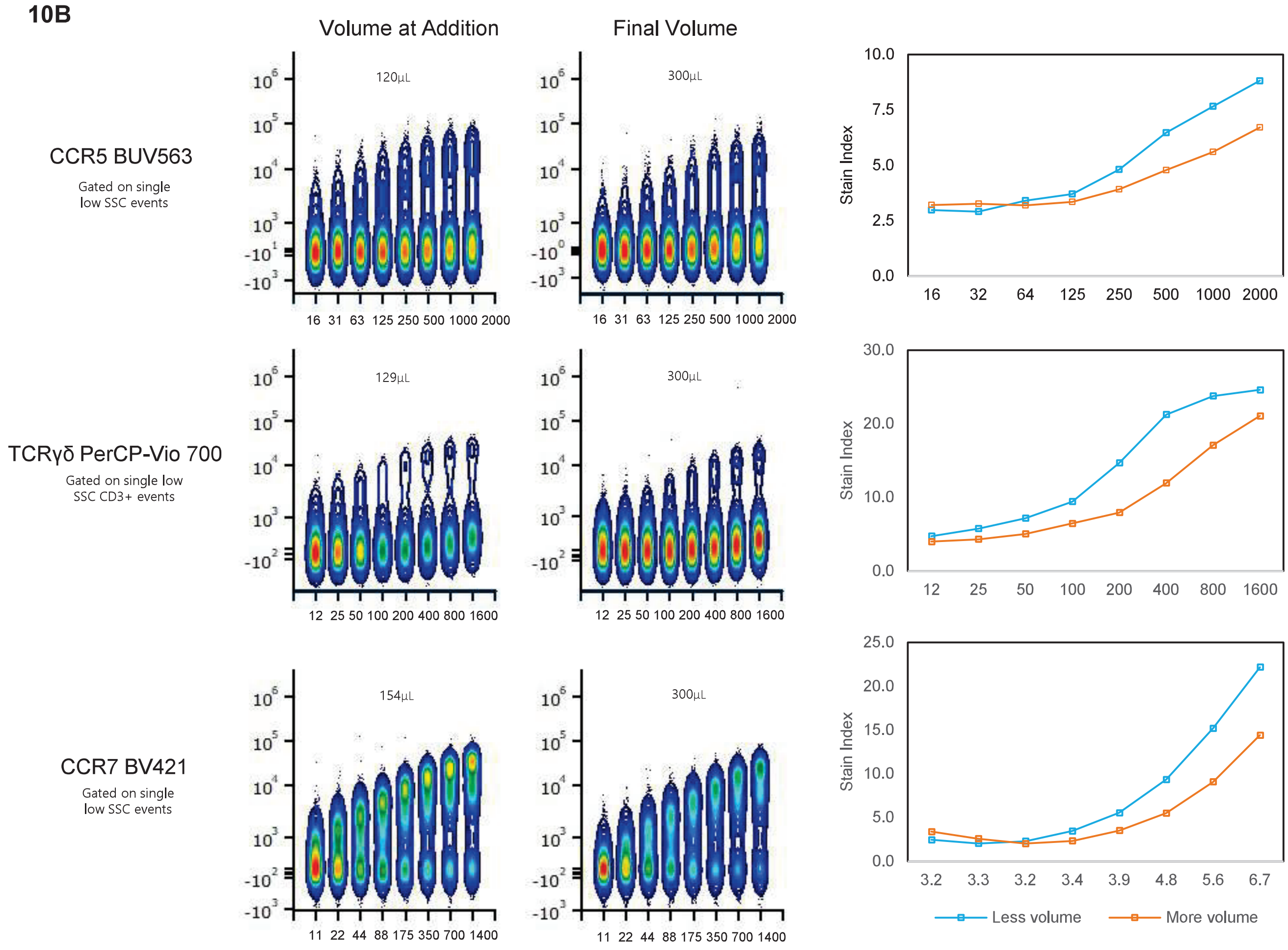

**Online Figure 10. Effects of Sequential Staining on Resolution**

**(A)** Optimization of the staining protocol was performed to improve the resolution of populations of interest. Improvements in staining quality of CD123 and IgG: comparison of resolution when antibody was added as part of the full cocktail (left) or added separately (right). **(B)** Volume impact on titration for reagents added in sequential steps was tested, comparing titers at the volume when the reagent is added vs. final volume (300µL). Concatenated plots are shown for CCR5 BUV563, CCR7 BV421, and TCRγδ PerCP-Vio 700, co-stained with CD3 FITC. Titer remains identical at the different volumes tested, while some differences in resolution are observed, confirming the utility of adding these reagents at earlier stages of the staining protocol.

**Figure 11)**. An elevated level of agreement was achieved, giving confidence that every marker's resolution was optimal, which is required for proper identification of every population of interest.

**f. Assessment of resolution of markers assigned to highly overlapping fluorochromes**

The impact of the spread introduced by using highly overlapping fluorochromes was assessed in the multicolor panel (**Online Figure 12A).** For all challenging fluorochrome combinations, the ability to identify populations was not impacted by overlapping fluorochromes. Based on our SSM calculation, PE/Cy5 had the highest potential spread impact on PE/Fire 640 (15.16). We verified that markers assigned to those fluorochromes were resolving successfully, including the low frequency population co-expressing CD103 and CD95. Optimal resolution was achieved also for other highly overlapping pairs, including BUV395-SparkUV387, Super Bright 436-eFluor 450, PE/Cy5-PerCP-Cy5.5, and BV510-BV480. For co-expressed markers assigned to overlapping fluorochromes, FMO controls were used to guide and/or assess positive and negative delineations. FMOs were used for all the new markers (CD103 PE/Fire 640, CD95 PE/Cy5, CD11b Spark UV 387, and DNAM-1 Real Blue 780), to evaluate whether the spillover received by those fluorochromes could be affecting resolution. The impact of spread in population resolution, evaluated by comparing the multicolor samples with FMOs of the highly overlapping fluorochromes, showed that for all the markers evaluated, identification of positive events was accurate and straightforward **(Online Figure 12B).**

**g. Comparison of resolution of common markers between OMIP-069 and the 45-color panel.**

Since OMIP-069 was used as a starting point, the resolution of the 37 shared markers was compared between the two panels. This comparison was aimed at ensuring that the addition of new fluorochromes and markers did not perturb the resolution of the original panel. Side-by-side

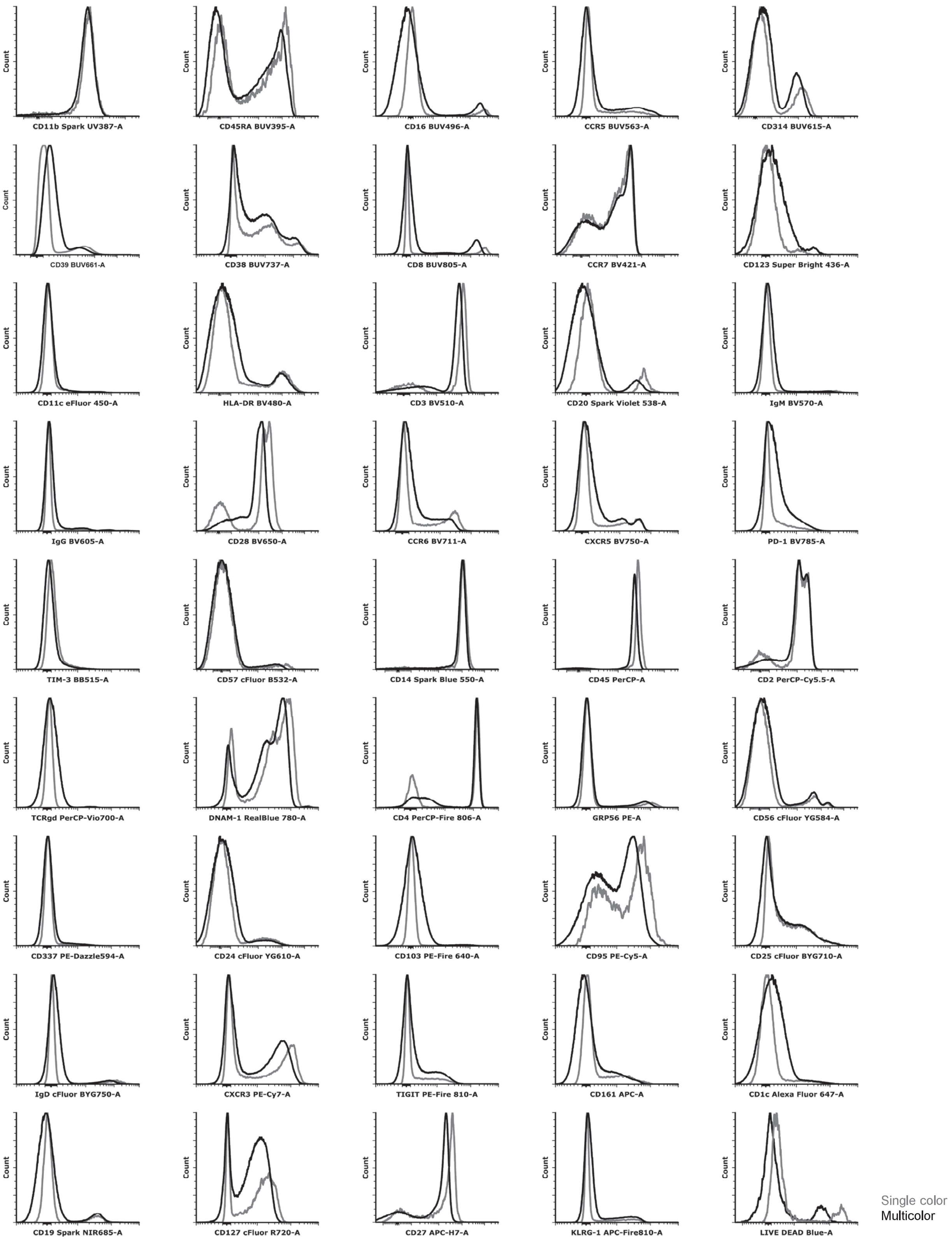

**Online Figure 11. Staining Resolution: Single Stained vs. Multicolor Samples**

As part of the panel optimization, each marker in the multicolor stained samples was evaluated to determine whether optimal resolution was maintained. To assess this, staining patterns were compared between the single stained controls (gray line) and the multicolor sample (black line) within the same experiment. The data in the overlaid histograms was gated on singlets and depending on the marker, lymphocyte or monocyte scatter gates. The data presented was obtained after the final titers were established and the staining protocol was optimized. Excellent level of agreement in resolution was found across all markers.

**12A**

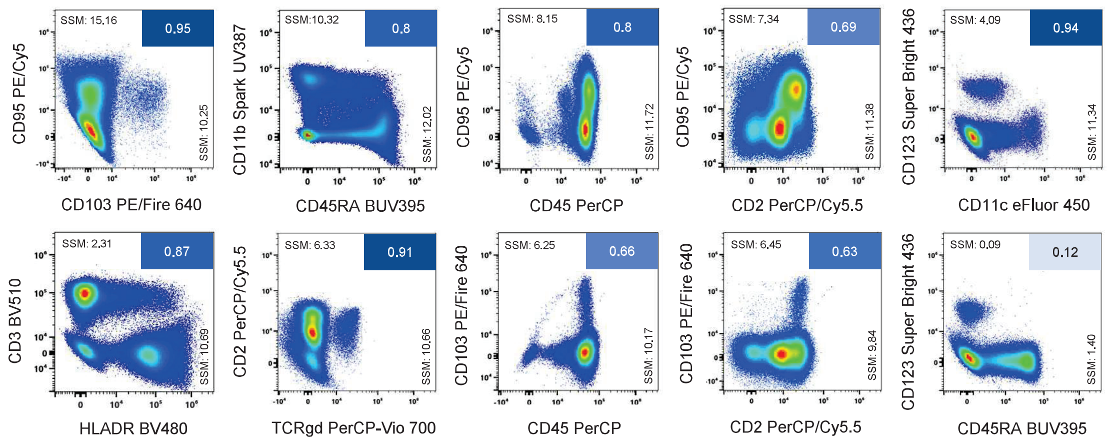

**12B**

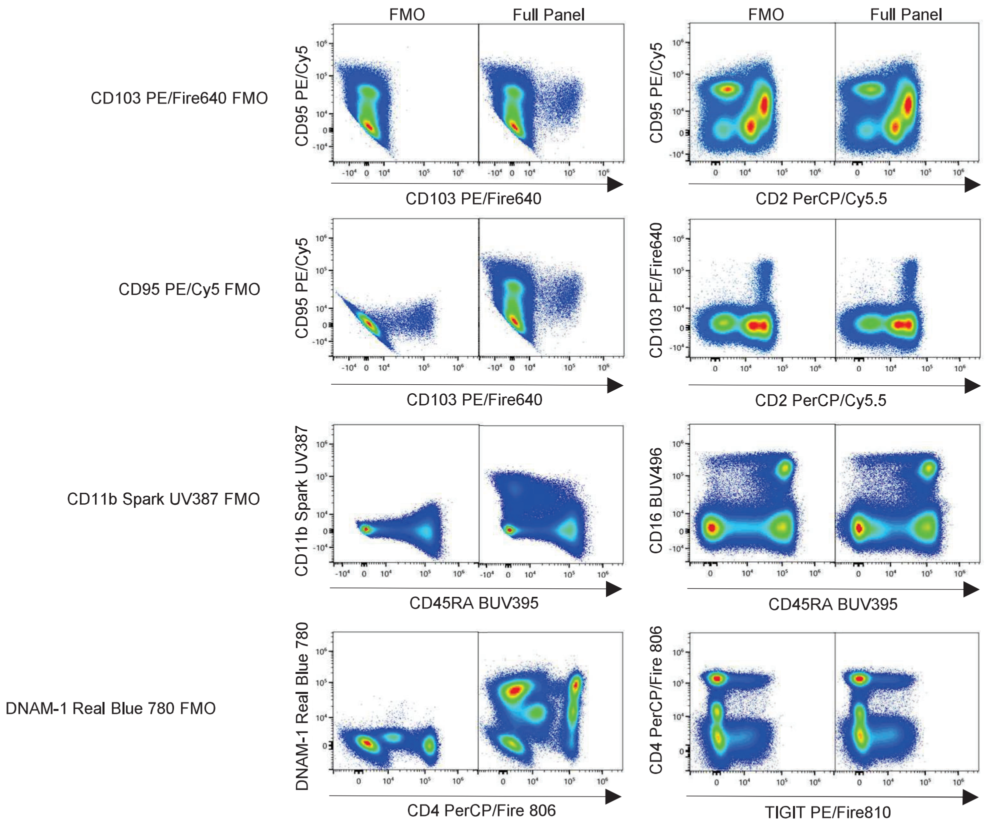

**Online Figure 12. Assessment of Resolution of Highly Overlapping Fluorochromes.**
The impact of spread introduced by using highly overlapping fluorochromes was assessed using the multicolor sample. **(A)** Fluorochrome combinations having the highest spillover spread values are presented. Plots are gated on singlets, live, non-aggregates, FSC/ SSC low/ intermediate, and CD45+, except for the plots displaying CD45 as a parameter in the x-axis, which are gated on live single low scatter events. Spillover spread values are indicated in each plot. Plots are organized from highest to lowest SSM. **(B)** FMOs were used to assess the impact of spread introduced by or into the newly selected fluorochromes: PE/Fire 640 into PE/Cy5 and PerCP Cy5.5; PE/Cy5 into PE/Fire 640; Spark UV 387 into BUV395, DNAM-1 Real Blue 780 into CD4 PerCP/Fire 806 and TIGIT PE/Fire 810. No significant changes in resolution were observed between the FMO controls and the full panel.

comparison is shown in **Online Figure 13.** All populations defined by the common markers in both panels showed equal resolution, confirming that the selection of new fluorochromes, panel design, and assay optimization steps were adequately performed.

### 5. 45-color assay performance

#### a. Manual gating of identified cellular subsets across donors

The entire gating strategy, as in the one published in OMP-069, is presented in **Online Figure 14**. First, cell doublets, red blood cells, dead cells, and CD45 negative events were gated out. Sequential gating plots are displayed for main lineages, including monocytes, DCs, ILCs, NK cells, NKT-like cells, T cells, B cells, and basophils. Each population was further characterized based on the presence or absence of various markers in the panel. Straightforward gating was achieved due to the outstanding resolution of each marker in the panel, from the top of the gating strategy to minor subsets. With the inclusion of other T cell markers, we characterized additional subsets from those described in OMIP-069. For example, mucosa-associated invariant T (MAIT) $CD8^+$ cells, were identified based on CD95, CD161, and KLRG1 co-expression. $CD8^+$ MAIT cells were further characterized based on the presence of CCR5, CCR6, CD127, and the bimodal expression of DNAM-1. $CD4^-CD8^-$ MAIT cells were identified using a similar gating strategy (**Online Figure 15)**.

T stem cell memory T cells (TSCM) are defined as $CD45RA^+CCR7^+CD28^+CD27^+$. In contrast to naive T cells (TN), TSCM express CD95, as an indicator of memory status. The present panel also allows identification of tissue-resident memory (TRM) cells, identified by expression of CD95, CD103, and DNAM-1. Subsets of TRM cells can be identified based on the presence of CCR5, CCR6, CD127, CD25, and CD28, as seen in the **Online Figure 16**. Terminal exhausted (TEX) T

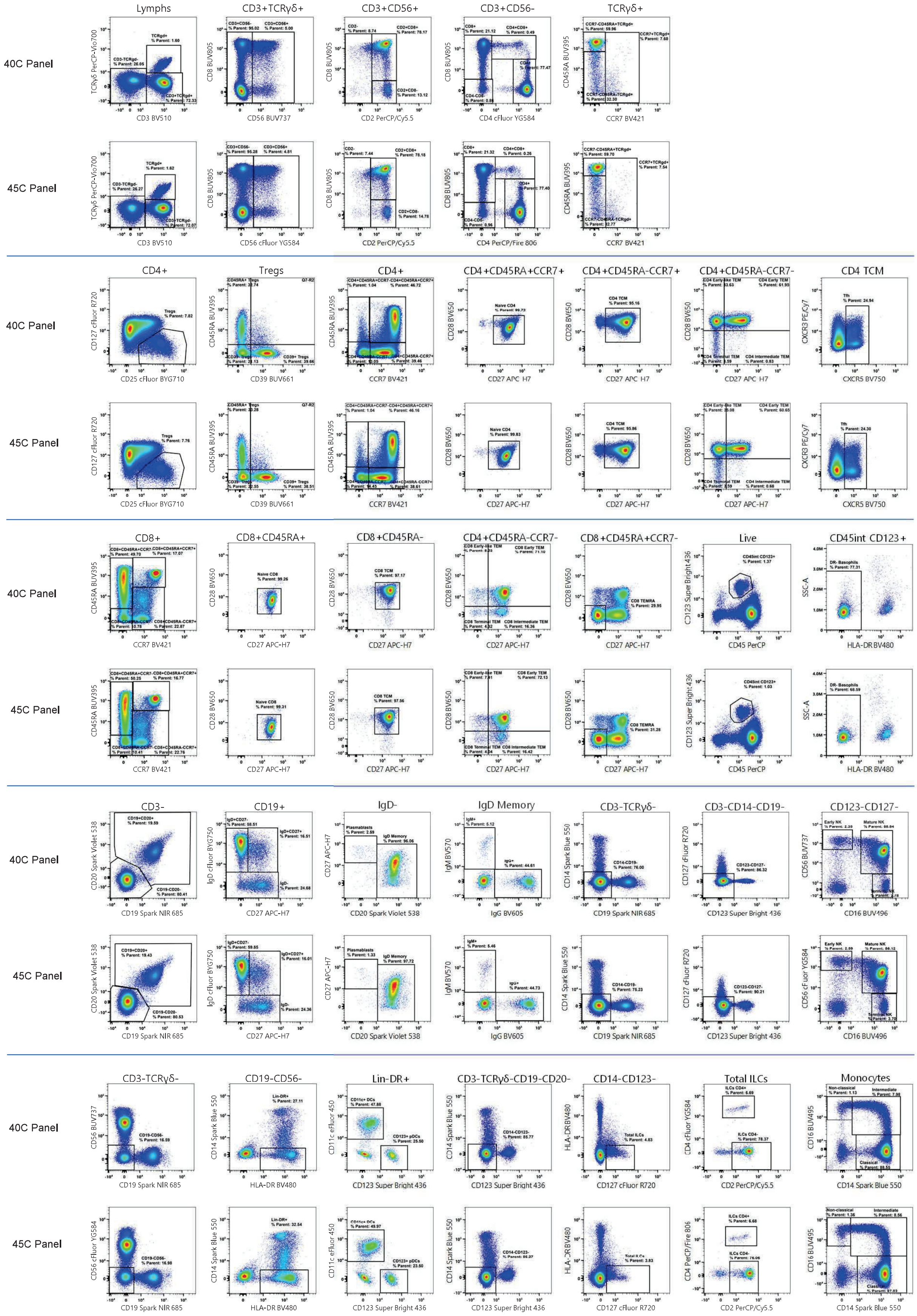

**Online Figure 13. Comparison of Marker Resolution between 40-color OMIP-069 and 45-color OMIP-xxx.** Visual comparison of the resolution of the common markers included in both panels (revised 40-color OMIP-069 and 45-color expansion) was performed. Note that the bi exponential scale was adjusted to be identical between both datasets. No significant differences in population definition or frequency were observed, indicating that the addition of new markers and fluorochromes did not perturb the original panel.

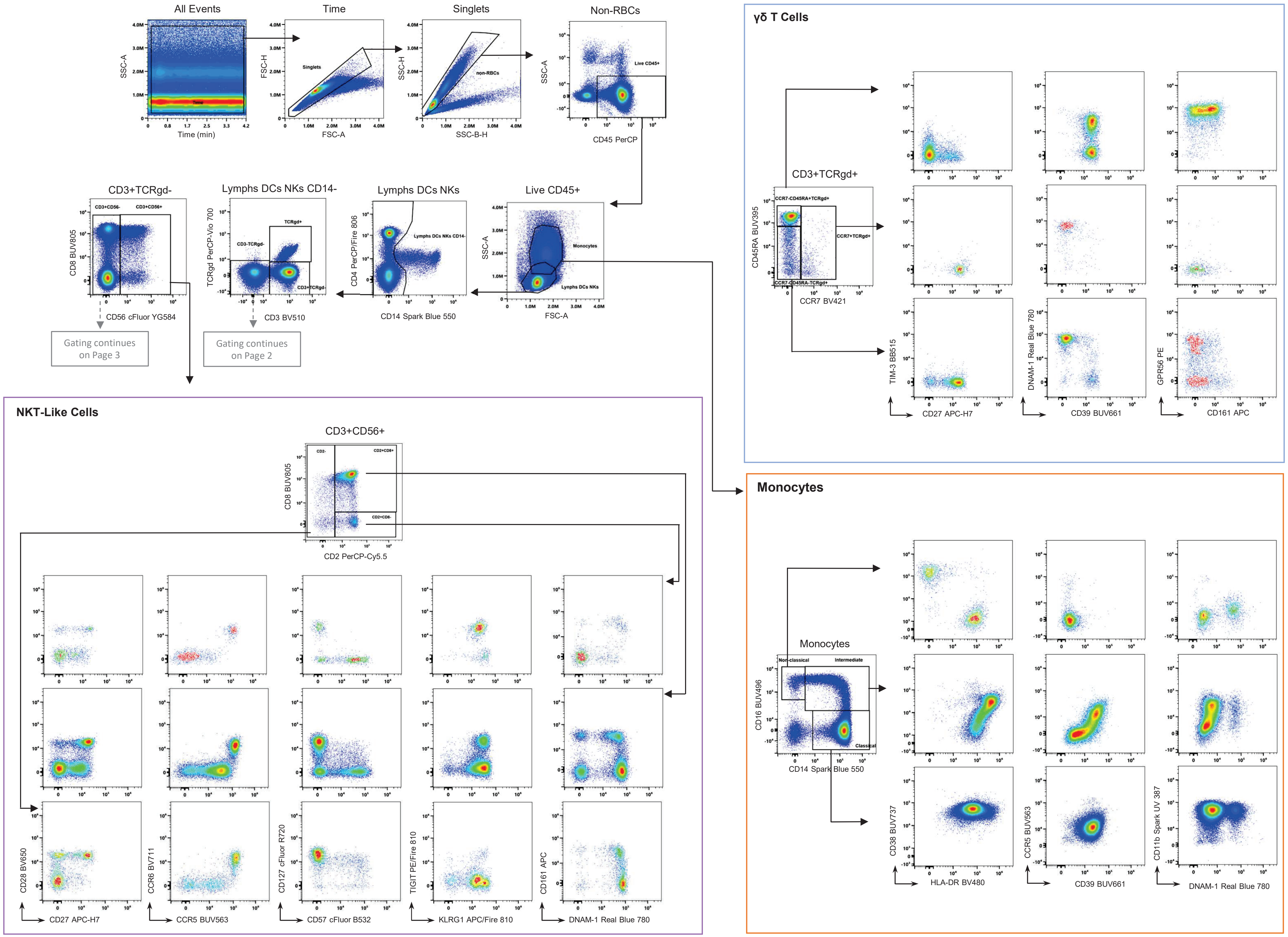
All Events
Time
Singlets
Non-RBCs
SSC-A
Time (min)
FSC-H
FSC-A
SSC-H
SSC-B-H
CD45 PerCP
Live CD45+
CD3+TCRgd-
Lymphs DCs NKs CD14-
Lymphs DCs NKs
CD8 BUV805
CD56 cFluor YG584
TCRgd PerCP-Vio 700
CD3 BV510
CD4 PerCP/Fire 806
CD14 Spark Blue 550
Monocytes
Gating continues on Page 3
Gating continues on Page 2
γδ T Cells
CD3+TCRgd+
CD45RA BUV395
CCR7 BV421
TIM-3 BB515
CD27 APC-H7
DNAM-1 Real Blue 780
CD39 BUV661
GPR56 PE
CD161 APC
NKT-Like Cells
CD3+CD56+
CD8 BUV805
CD2 PerCP-Cy5.5
CD28 BV650
CD27 APC-H7
CCR6 BV711
CCR5 BUV563
CD127 cFluor R720
CD57 cFluor B532
TIGIT PE/Fire 810
KLRG1 APC/Fire 810
CD161 APC
DNAM-1 Real Blue 780
Monocytes
CD16 BUV496
CD14 Spark Blue 550
Non-classical
Intermediate
Classical
CD38 BUV737
HLA-DR BV480
CCR5 BUV563
CD39 BUV661
CD11b Spark UV 387
DNAM-1 Real Blue 780

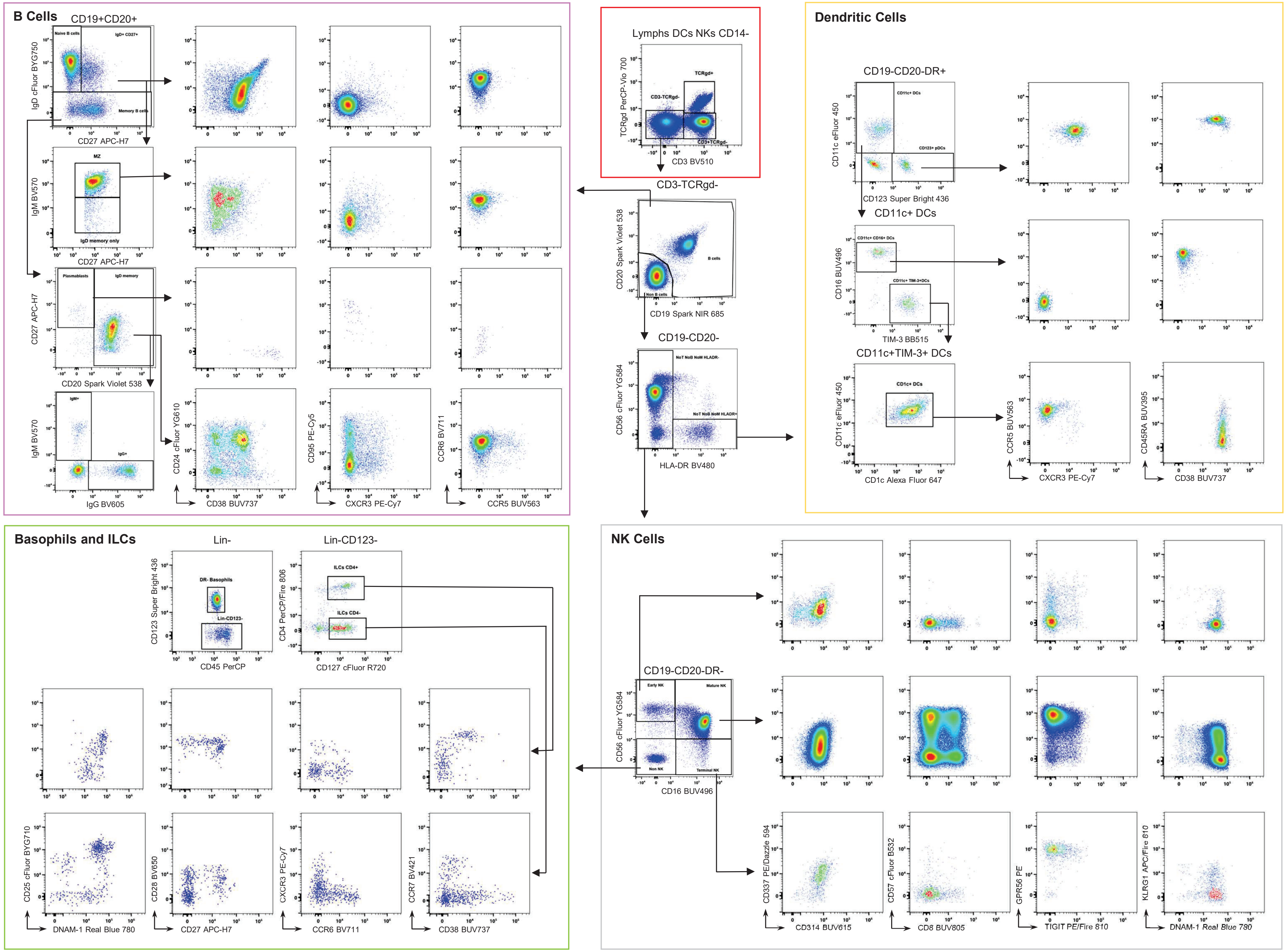

B Cells
CD19+CD20+
IgD cFluor BYG750
CD27 APC-H7
Memory B cells
IgM BV570
MZ
IgD memory only
CD27 APC-H7
Plasmablasts
IgD memory
CD20 Spark Violet 538
IgM+
IgG+
IgG BV605
CD24 cFluor YG610
CD38 BUV737
CD95 PE-Cy5
CXCR3 PE-Cy7
CCR6 BV711
CCR5 BUV563
Lymphs DCs NKs CD14-
TCRgd PerCP-Vio 700
TCRgd+
CD3-TCRgd-
CD3+TCRgd-
CD3 BV510
CD3-TCRgd-
CD20 Spark Violet 538
B cells
Non B cells
CD19 Spark NIR 685
CD19-CD20-
CD56 cFluor YG584
HLA-DR BV480
Dendritic Cells
CD19-CD20-DR+
CD11c eFluor 450
CD11c+ DCs
CD123+ pDCs
CD123 Super Bright 436
CD11c+ DCs
CD16 BUV496
CD11c+ CD16+ DCs
CD11c+ TIM-3+DCs
TIM-3 BB515
CD11c+TIM-3+ DCs
CD11c eFluor 450
CD1c Alexa Fluor 647
CCR5 BUV563
CXCR3 PE-Cy7
CD45RA BUV395
CD38 BUV737
Basophils and ILCs
Lin-
Lin-CD123-
CD123 Super Bright 436
DR- Basophils
Lin-CD123-
CD45 PerCP
CD4 PerCP/Fire 806
ILCs CD4+
ILCs CD4-
CD127 cFluor R720
CD25 cFluor BYG710
DNAM-1 Real Blue 780
CD28 BV650
CD27 APC-H7
CXCR3 PE-Cy7
CCR6 BV711
CCR7 BV421
CD38 BUV737
NK Cells
CD19-CD20-DR-
CD56 cFluor YG584
Early NK
Mature NK
Non NK
Terminal NK
CD16 BUV496
CD337 PE/Dazzle 594
CD314 BUV615
CD57 cFluor B532
CD8 BUV805
GPR56 PE
TIGIT PE/Fire 810
KLRG1 APC/Fire 810
DNAM-1 Real Blue 780

14

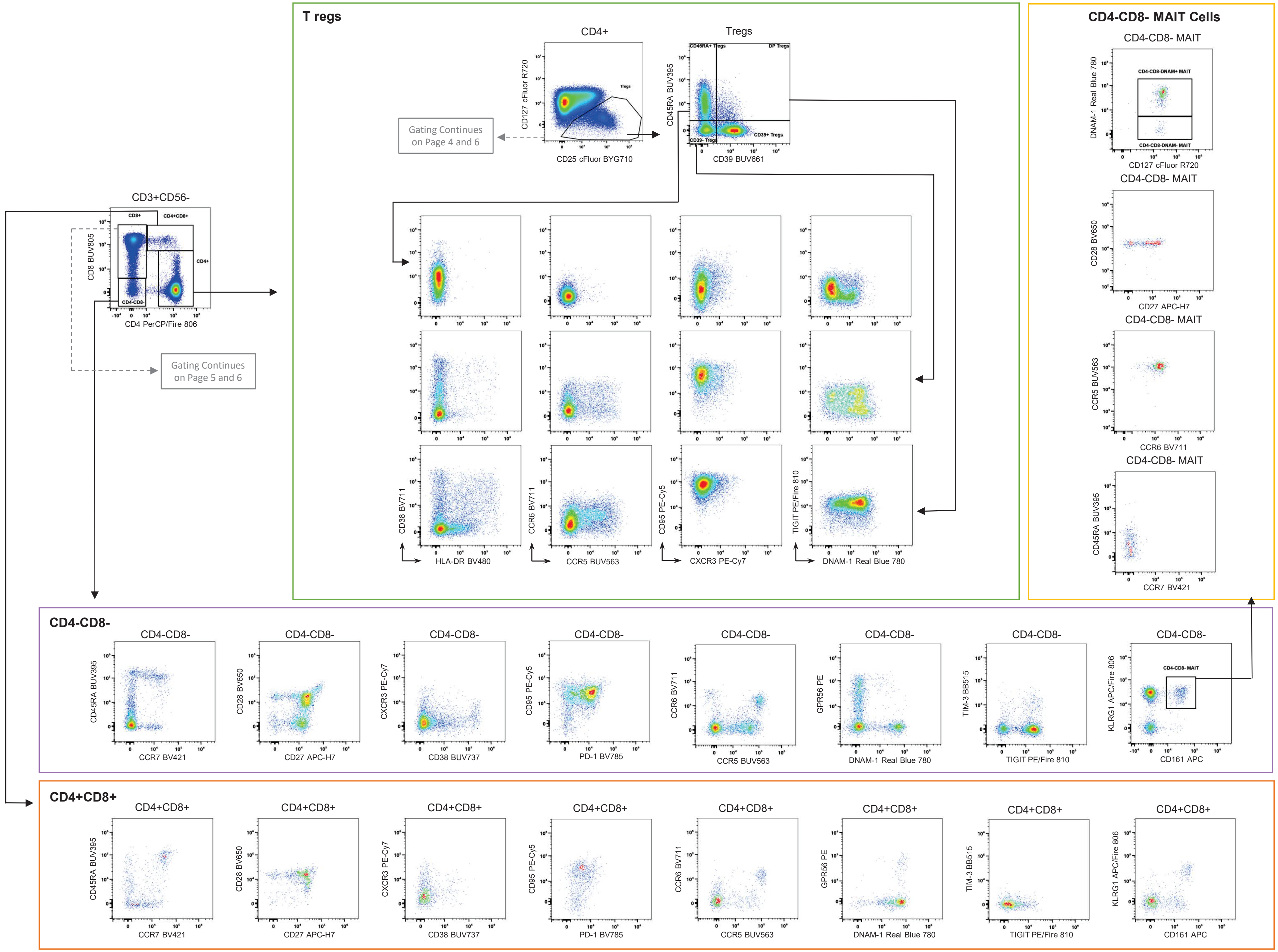
T regs
CD4+
Tregs
CD127 cFluor R720
CD25 cFluor BYG710
Gating Continues on Page 4 and 6
CD45RA BUV395
CD39 BUV661
CD45RA+ Tregs
DP Tregs
CD39- Tregs
CD39+ Tregs
CD38 BV711
HLA-DR BV480
CCR6 BV711
CCR5 BUV563
CD95 PE-Cy5
CXCR3 PE-Cy7
TIGIT PE/Fire 810
DNAM-1 Real Blue 780
CD4-CD8- MAIT Cells
CD4-CD8- MAIT
DNAM-1 Real Blue 780
CD127 cFluor R720
CD4-CD8-DNAM+ MAIT
CD4-CD8-DNAM- MAIT
CD28 BV650
CD27 APC-H7
CCR5 BUV563
CCR6 BV711
CD45RA BUV395
CCR7 BV421
CD3+CD56-
CD8 BUV805
CD4 PerCP/Fire 806
CD8+
CD4+CD8+
CD4+
CD4-CD8-
Gating Continues on Page 5 and 6
CD4-CD8-
CXCR3 PE-Cy7
CD38 BUV737
CD95 PE-Cy5
PD-1 BV785
GPR56 PE
TIM-3 BB515
TIGIT PE/Fire 810
KLRG1 APC/Fire 806
CD161 APC
CD4-CD8- MAIT
CD4+CD8+

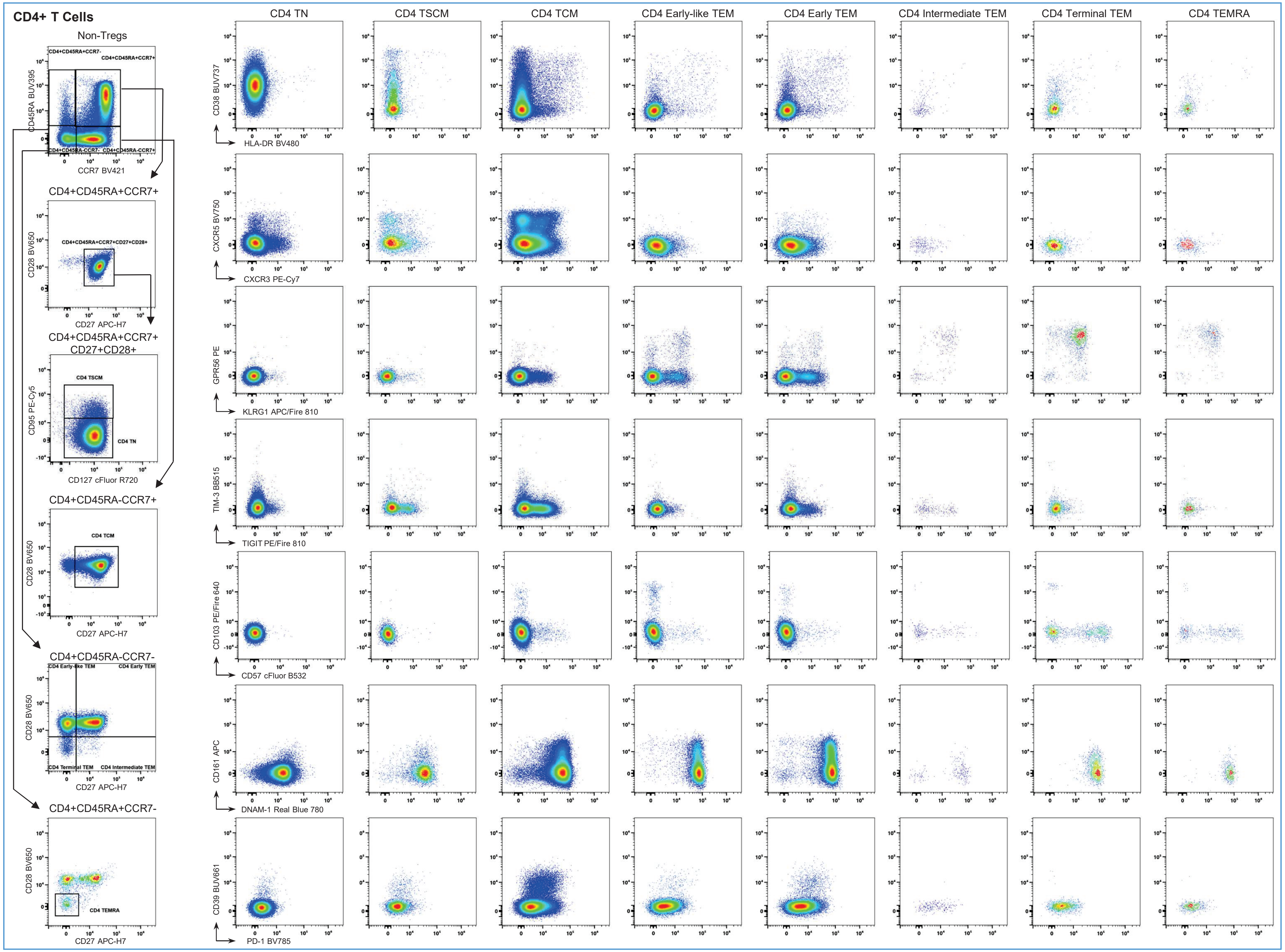

CD4+ T Cells
Non-Tregs
CD4+CD45RA+CCR7-
CD4+CD45RA+CCR7+
CD4+CD45RA-CCR7-
CD4+CD45RA-CCR7+
CD45RA BUV395
CCR7 BV421
CD4+CD45RA+CCR7+
CD4+CD45RA+CCR7+CD27+CD28+
CD28 BV650
CD27 APC-H7
CD4+CD45RA+CCR7+
CD27+CD28+
CD4 TSCM
CD4 TN
CD95 PE-Cy5
CD127 cFluor R720
CD4+CD45RA-CCR7+
CD4 TCM
CD28 BV650
CD27 APC-H7
CD4+CD45RA-CCR7-
CD4 Early-like TEM
CD4 Early TEM
CD4 Terminal TEM
CD4 Intermediate TEM
CD28 BV650
CD27 APC-H7
CD4+CD45RA+CCR7-
CD4 TEMRA
CD28 BV650
CD27 APC-H7
CD4 TN
CD4 TSCM
CD4 TCM
CD4 Early-like TEM
CD4 Early TEM
CD4 Intermediate TEM
CD4 Terminal TEM
CD4 TEMRA
CD38 BUV737
HLA-DR BV480
CXCR5 BV750
CXCR3 PE-Cy7
GPR56 PE
KLRG1 APC/Fire 810
TIM-3 BB515
TIGIT PE/Fire 810
CD103 PE/Fire 640
CD57 cFluor B532
CD161 APC
DNAM-1 Real Blue 780
CD39 BUV661
PD-1 BV785

14

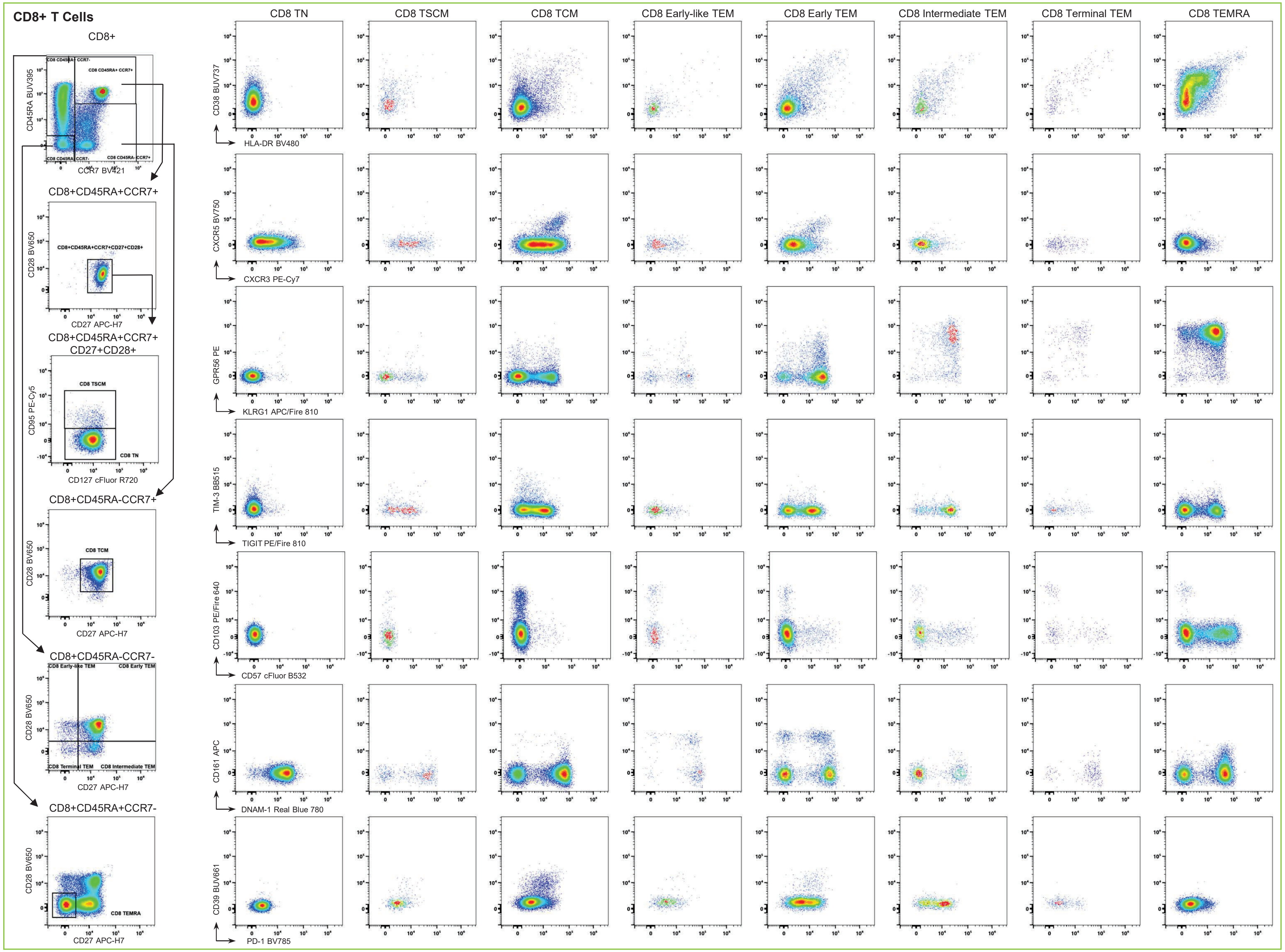
CD8+ T Cells
CD8+
CD45RA BUV395
CCR7 BV421
CD8+CD45RA+CCR7+
CD28 BV650
CD27 APC-H7
CD8+CD45RA+CCR7+ CD27+CD28+
CD95 PE-Cy5
CD127 cFluor R720
CD8 TSCM
CD8 TN
CD8+CD45RA-CCR7+
CD8 TCM
CD8+CD45RA-CCR7-
CD8 Early-like TEM
CD8 Early TEM
CD8 Terminal TEM
CD8 Intermediate TEM
CD8+CD45RA+CCR7-
CD8 TEMRA
CD8 TN
CD8 TSCM
CD8 TCM
CD8 Early-like TEM
CD8 Early TEM
CD8 Intermediate TEM
CD8 Terminal TEM
CD8 TEMRA
CD38 BUV737
HLA-DR BV480
CXCR5 BV750
CXCR3 PE-Cy7
GPR56 PE
KLRG1 APC/Fire 810
TIM-3 BB515
TIGIT PE/Fire 810
CD103 PE/Fire 640
CD57 cFluor B532
CD161 APC
DNAM-1 Real Blue 780
CD39 BUV661
PD-1 BV785

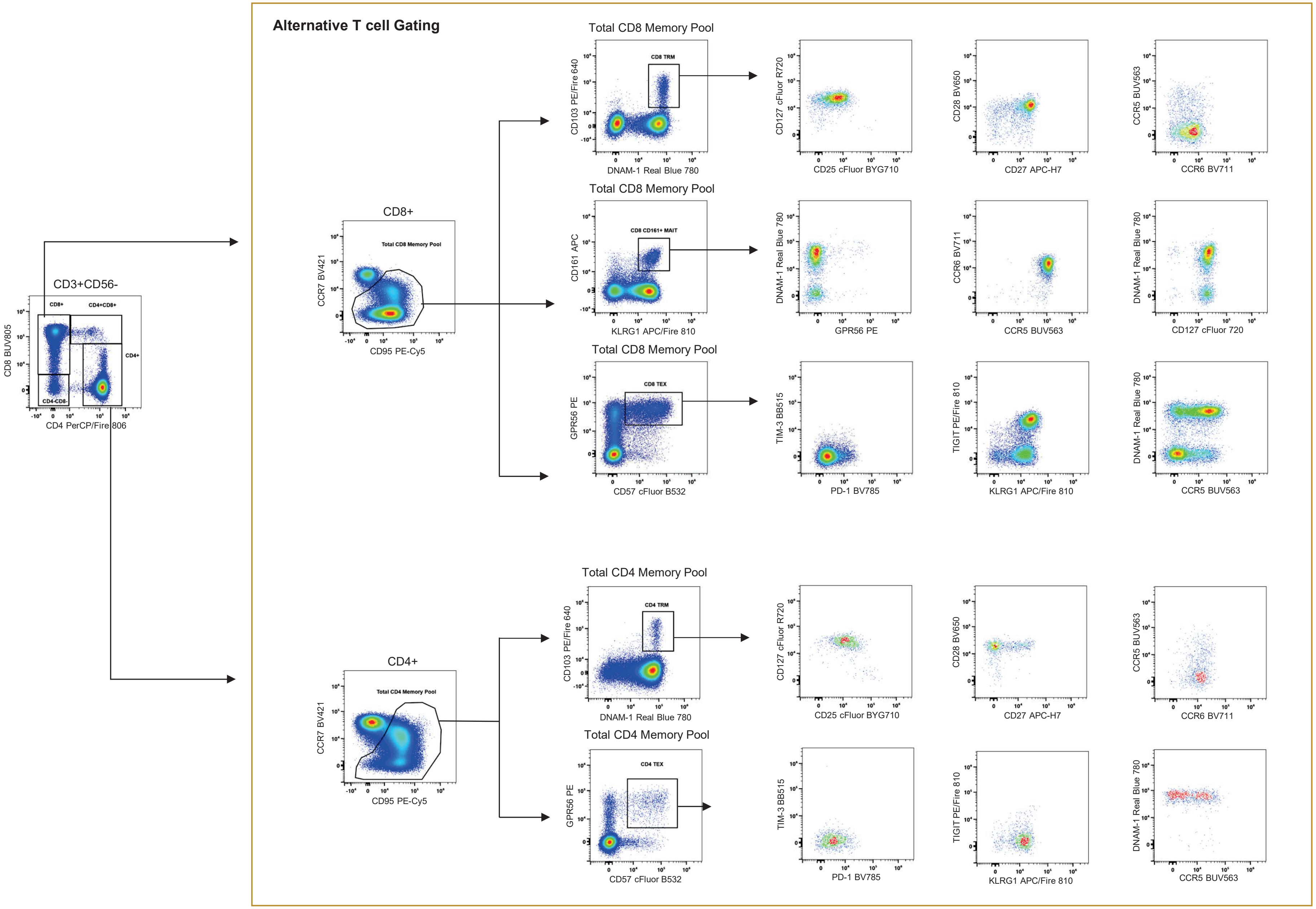

**Online Figure 14. Full Manual Gating Strategy.** Following the gating scheme presented in Figure 1A, the different cellular subsets were further characterized by manual gating. For each main population, the markers that showed different levels of expression across the various subsets of specific populations are presented in sequential gating.

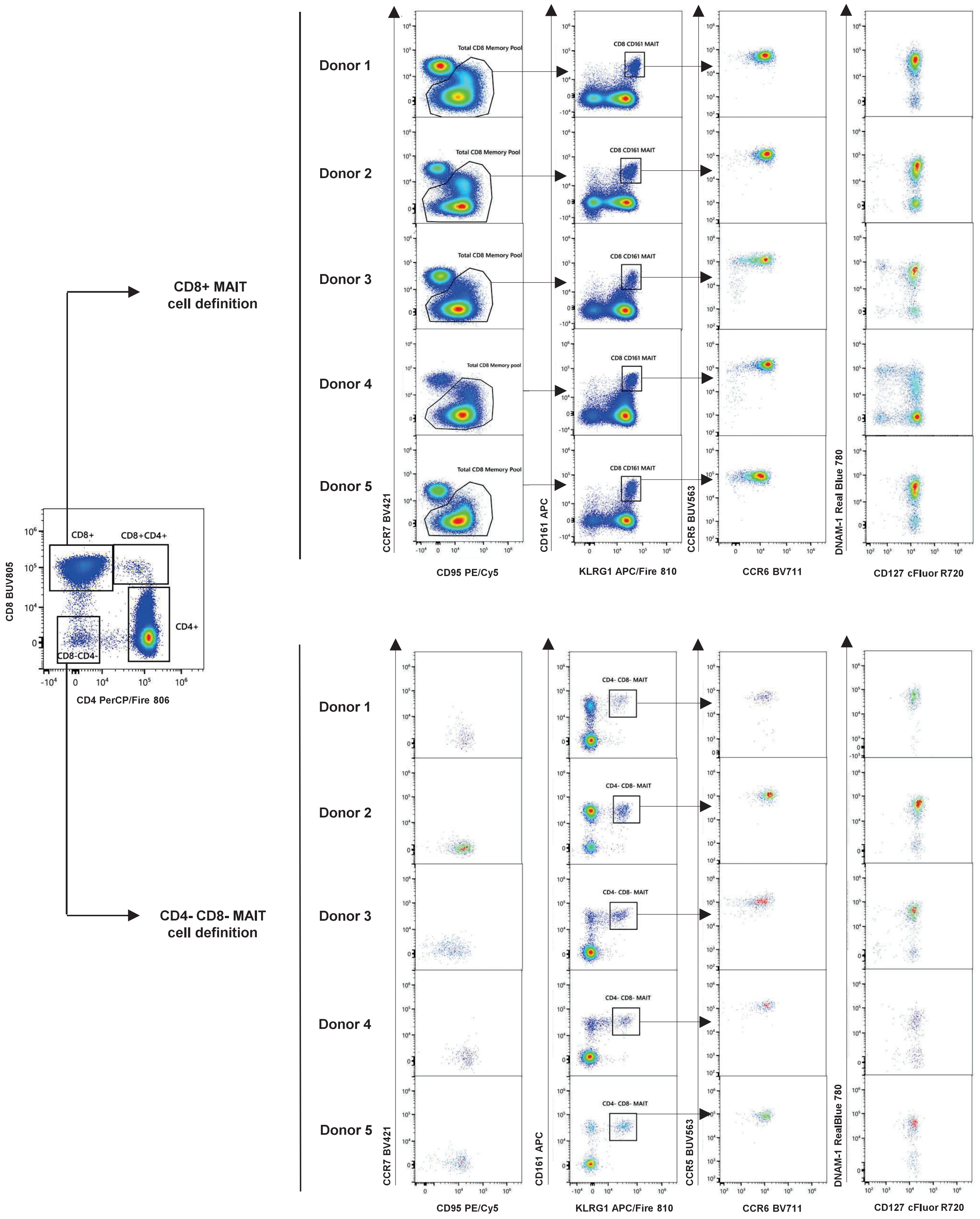

**Online Figure 15. Identification and Characterization of Mucosa-Associated Invariant T Cells (MAIT).** The manual gating strategy to identify MAIT CD8⁺ and CD4- CD8-T cells is presented. Data from 5 donors illustrates variation in the phenotype of these cells. MAIT cells are identified from the total memory pool by the co-expression of CD161 and KLRG1. Expression of CCR5, CCR6, DNAM-1, and CD127 is shown for both CD8+ MAIT and CD4- CD8- MAIT cells.

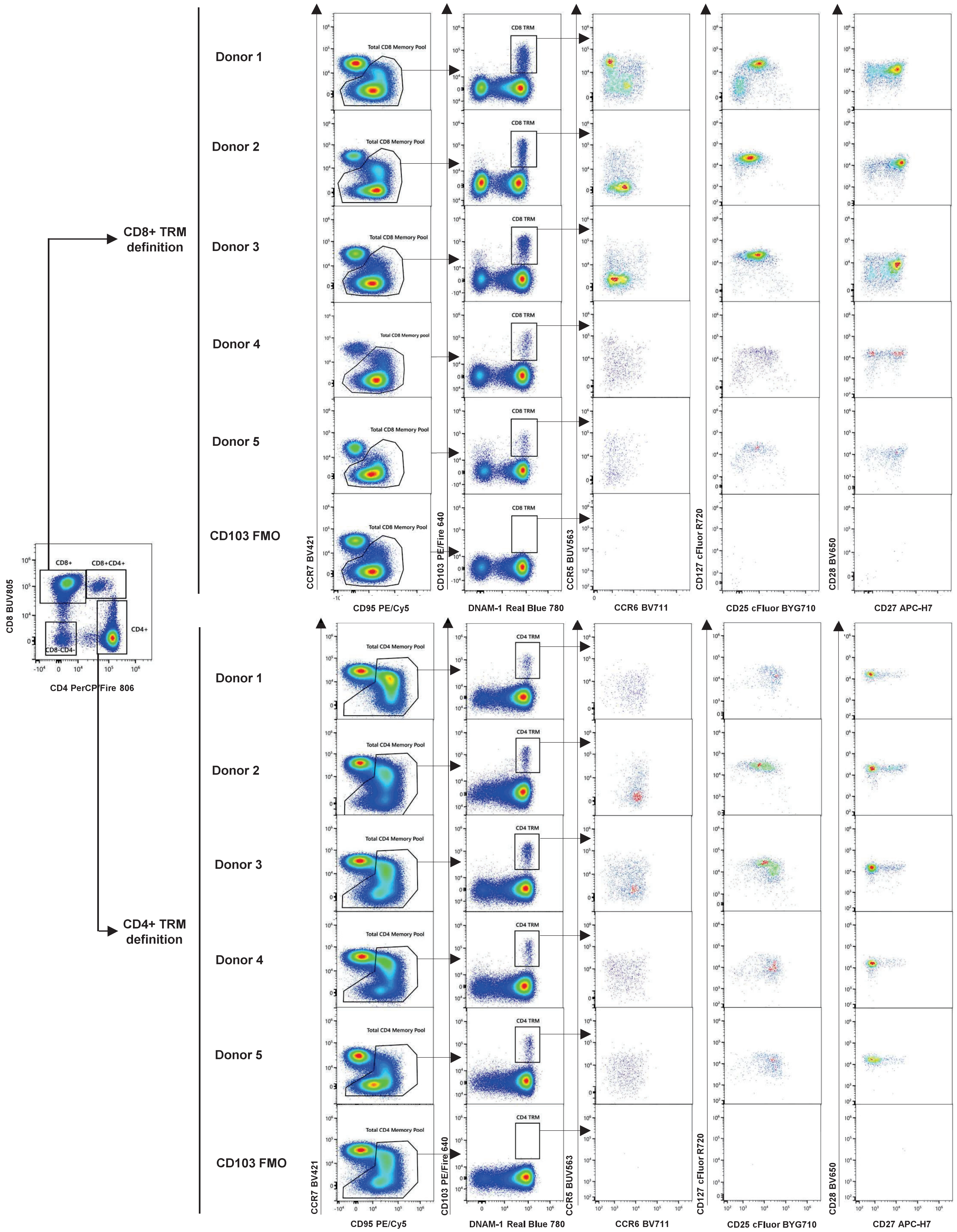

**Online Figure 16. Identification and Characterization of Tissue-Resident Memory T Cells (TRM).**

The manual gating strategy to identify TRM CD8+ and CD4+ T cells is presented. Data from 5 donors illustrates variation in the phenotype of these cells. Since the identification of this subset relies on CD103 PE Fire 640 and this fluorochrome receives significant spread from other fluorochromes, the FMO control is presented. TRM cells are identified from the total memory pool by the co-expression of CD103 and DNAM-1. Expression of CCR5, CCR6, CD25, CD127, CD28, and CD27 is shown for both CD8+ and CD4+ TRM cells.

cells can be identified by co-expression of CD95, CD57, and GPR56, with further sub setting based on the presence of PD-1, KLRG1, DNAM-1, and CCR5 (**Online Figure 17).**

Finally, a deep characterization of TSCM cells is shown in **Online Figure 18**. The panel supports clear delineation of T cell memory subpopulations using the newly incorporated markers. Moreover, insights on cell differentiation can be obtained out of the naïve (TN), stem cell memory (TSCM), effector memory (TEM), and central memory (TCM) $CD4^+$ or $CD8^+$ T cells, based on the expression of markers like TIM-3, PD-1, CD161, CD57, TIGIT, KLRG1, CD103, and DNAM-1. For example, in **Online Figure 19A,** we observed for CD8 T cells that TIM-3 is only expressed in TCM, while CD161 was present only in early effector TEM. TIGIT and CD57 facilitate further exhaustion delineation of 4 subsets within intermediate and CD45RA+ terminal TEM cells. GPR56 was confirmed as a marker terminal effector differentiation, being present only intermediate and terminal effector cells, while CD25 and CCR6 were expressed more in early effector populations. Unique patterns were also observed in the range of memory CD4+ T cell subsets, as shown in **Online Figure 19B.**

#### b. Clustering and dimensionality reduction analysis

Although manual analysis of this 45-color panel allows clear identification of many distinct subsets of cells, algorithms such as FlowSOM (6) and UMAP (7), can be used to explore deeper complex multiparametric datasets, in an unbiased and semi-unsupervised manner (8-10). Data were analyzed using an OMIQ pipeline ((https://www.omiq.ai/), like the one published in OMIP-069 (1). After unmixing and scaling, PeacoQC package was used for QC purposes, to check signal stability prior clustering analysis (11). Main lineages were manually identified, upon removal of dead cells, red blood cells, CD45 negative events, and aggregates. To explore the heterogeneity of the memory pool, FlowSOM and UMAP analysis were completed separately in CD8+ T cells, CD4+ T cells, and NKT-like T cells, with identification of 30 distinct metaclusters per population. First, we will focus the

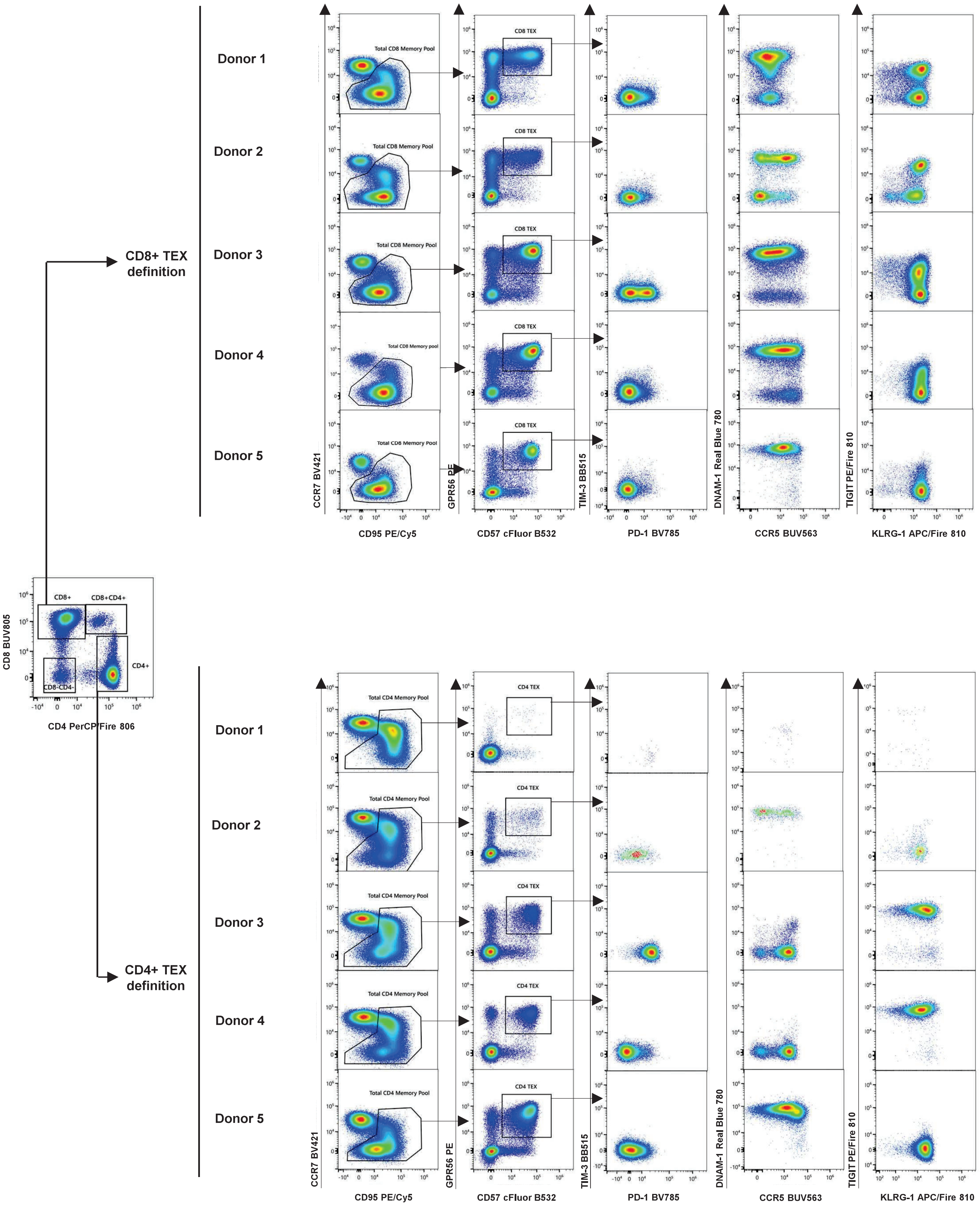

**Online Figure 17. Identification and Characterization of Terminal Exhausted (TEX) T cells.**
The manual gating strategy to identify TEX CD8+ and CD4+ T cells is presented. Data from 5 donors illustrates variation in the phenotype of these cells. TEX cells are identified from the total memory pool by the co-expression of CD57 and GPR56. Expression of TIM-3, PD-1, DNAM-1, CCR5, TIGIT, and KLRG-1 is shown for both CD8+ and CD4+ TEX cells.

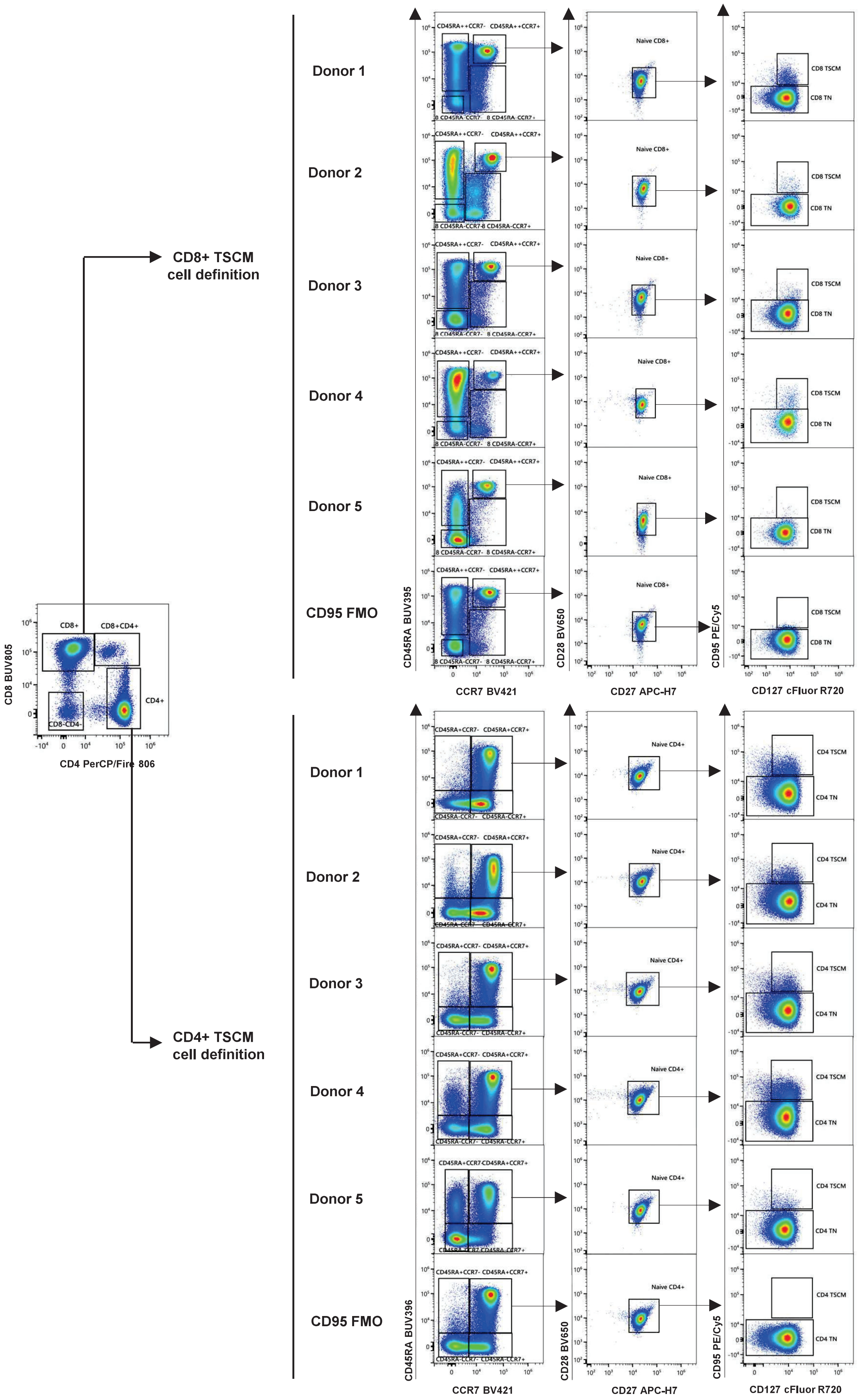

**Online Figure 18. Identification T Stem Cell Memory Cells (TSCM).**

The manual gating strategy to identify TSCM CD8+ and CD4+ T cells is presented. Data from 5 donors illustrates variation in frequency of these populations. Because CD95 is the key marker for identification of these cells and it is expressed at low levels, an FMO control was run to validate the positioning of the gate. TSCM cells are identified from by the co-expression of CD45RA, CCR7, CD27, CD28, and CD95. TN cells have a similar phenotype, except for the lack of CD95 expression.

19A

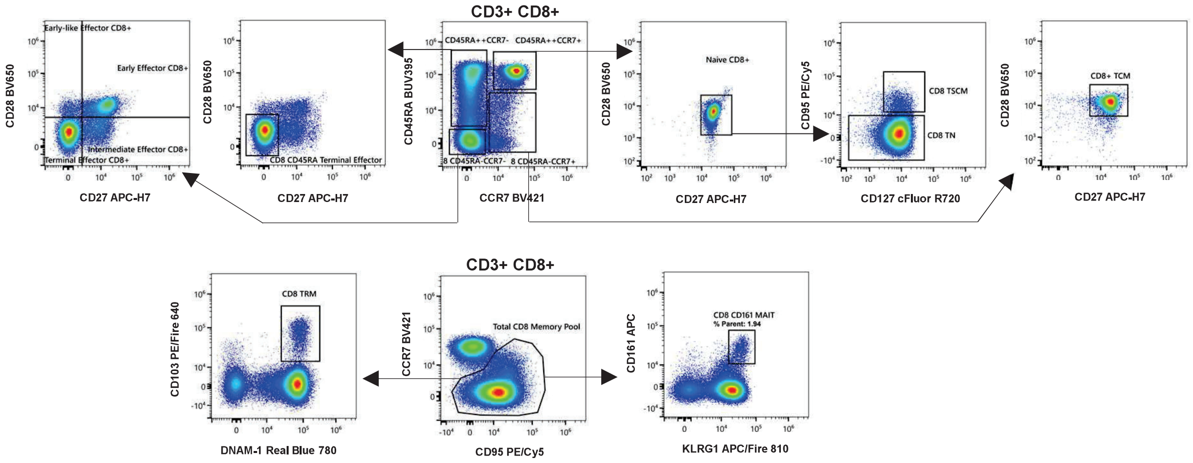

**Exhaustion and Activation in CD8+ T cells**

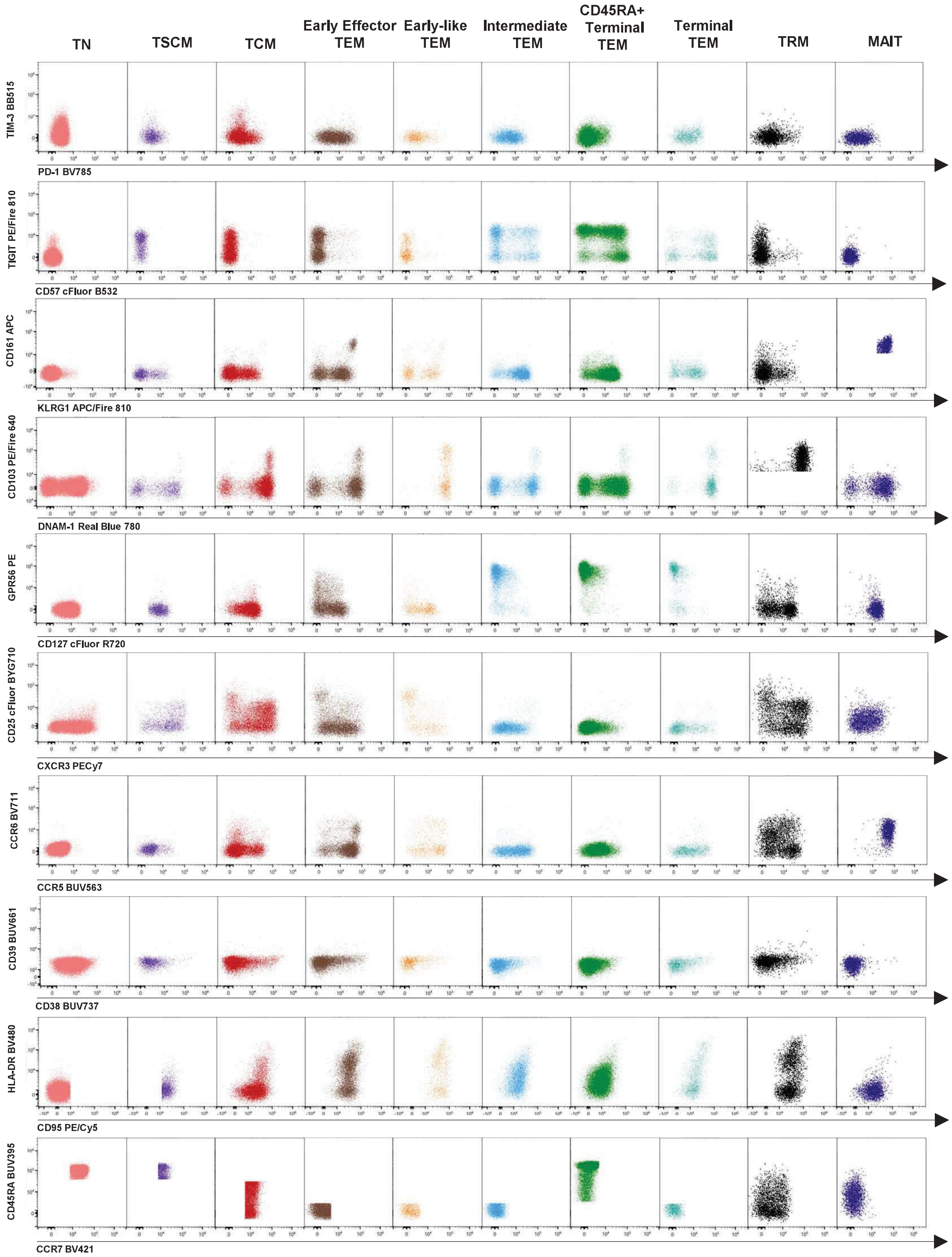

**19B**

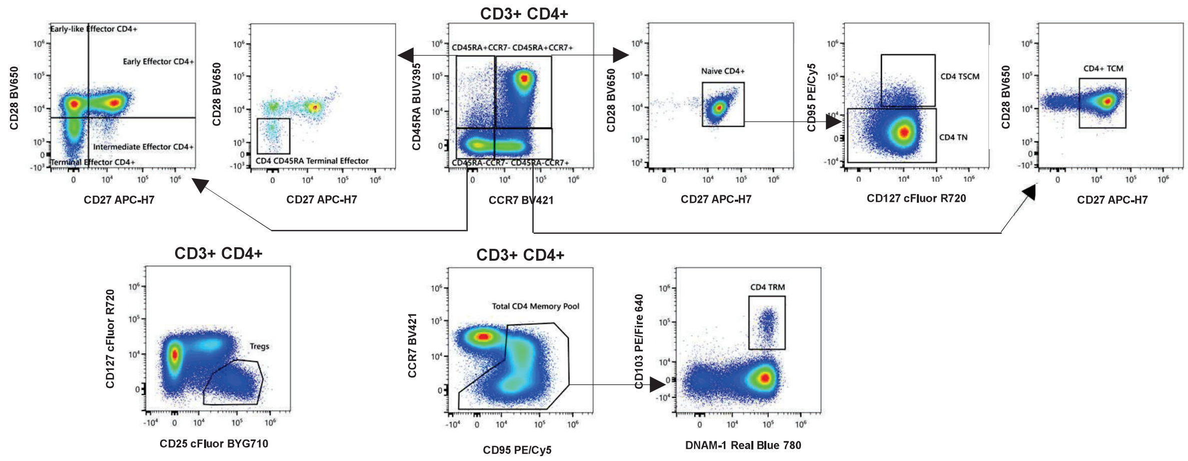

**Exhaustion and Activation in CD4+ T cells**

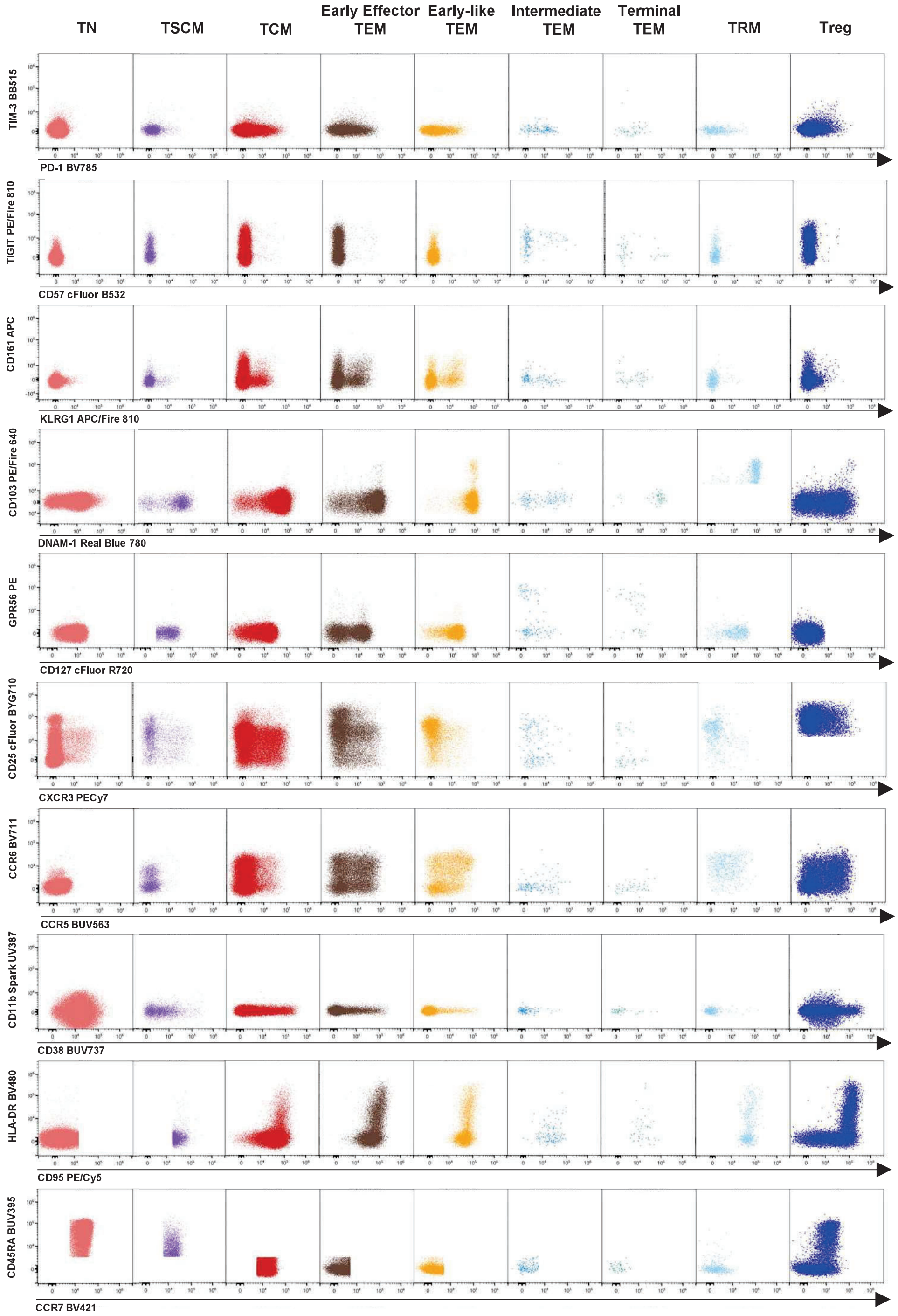

**Online Figure 19. Characterization of Exhaustion and Activation Phenotypes in Various T Cell Subsets** **(A)** The gating strategy to identify TN, TSMC, TCM, early-effector TEM, early-like TEM, intermediate TEM, CD45RA+ Terminal TEM, Terminal TEM, TRM and MAIT is presented for CD8$^{+}$ T cells**.** The changes in level of expression of various markers throughout those different subsets is illustrated in dot plots, based on the expression of TIM-3, PD-1, CD57, TIGIT, CD161, KLRG1, CD103, DNAM-1, GPR56, CD127, CD25, CXCR3, CCR6, CCR5, CD39, CD38, HLA-DR, CD95, CD45RA, and/or CCR7. Each T cell subset is identified with a different color. **(B)** A similar analysis is shown for CD4$^{+}$ T cell subsets (TN, TSCM, TCM, TEM, TRM, and Tregs).

discussion on CD4 T cells. **Online Figure 20A** shows FlowSOM metaclusters projected into 2 UMAP dimensions. The UMAP overlaid scatterplot shows 4 donors concatenated, as an example of the high consistency in cluster location and size across samples (**Online Figure 20B**). Four donors were included in the clustering analysis, since their samples were stained and acquired on the same day. A UMAP for each individual donor is shown in **Online Figure 20C**. Using concatenated files, we generated a hierarchically clustered heatmap for metacluster description. The marker expression intensity is displayed on a scale from red (negative) to green (positive) to indicate relative magnitude (**Online Figure 20D)**. We were able to identify subpopulations out of the classical TN, TSCM, TCM, or TEM subsets, based on the new markers included in the panel. As expected, CD45RA, CCR7, CD28, and CD27 facilitate cluster definition according to memory/effector status. For example, CD27, CD28, CD45RA and CCR7 are not present in metacluster 18, a hallmark for terminal effector cells, accompanied by expression of CCR5, CD57, PD-1, KLRG1, GPR56 and TIGIT, a phenotype highly suggestive of cellular exhaustion. Out of the new markers in the panel, TIGIT was present in 10 of the clusters, DNAM-1 in 21 (with high expression in 10), CD103 and CD161 in 3, GPR56 in 4, and KLRG1 in 5. Of note, CD11b and TIM-3 were only detected in one cluster (metacluster 9). As seen in the heatmap, cluster definition was also supported by markers present in the original 40-color panel, such as CD25, CD127, HLA-DR, CCR5, CCR6, CXCR3, CXCR5, CD38, CD39, CD57, CD95, and CD337. In CD4+ T cells, effector metaclusters 18, 29, and 30 were distant by expression of elevated levels of CD103, consistent with the manual identification of resident cells in the sample. The terminal effector metaclusters 13, 21, 22, 24, and 28 were characterized by high levels of KLRG1, with on/off expression of other terminal differentiation markers, such as GPR56, CD57, CCR5, TIGIT, and PD-1. In contrast, the abundance of CD45RA, CCR7, CD27, and CD28 was present in metacluster 5, classified as naive T cells, and characterized by the absence of CD95 and all the other terminal differentiation markers. As expected, the identified metaclusters displayed marker profiles reminiscent of all memory subsets

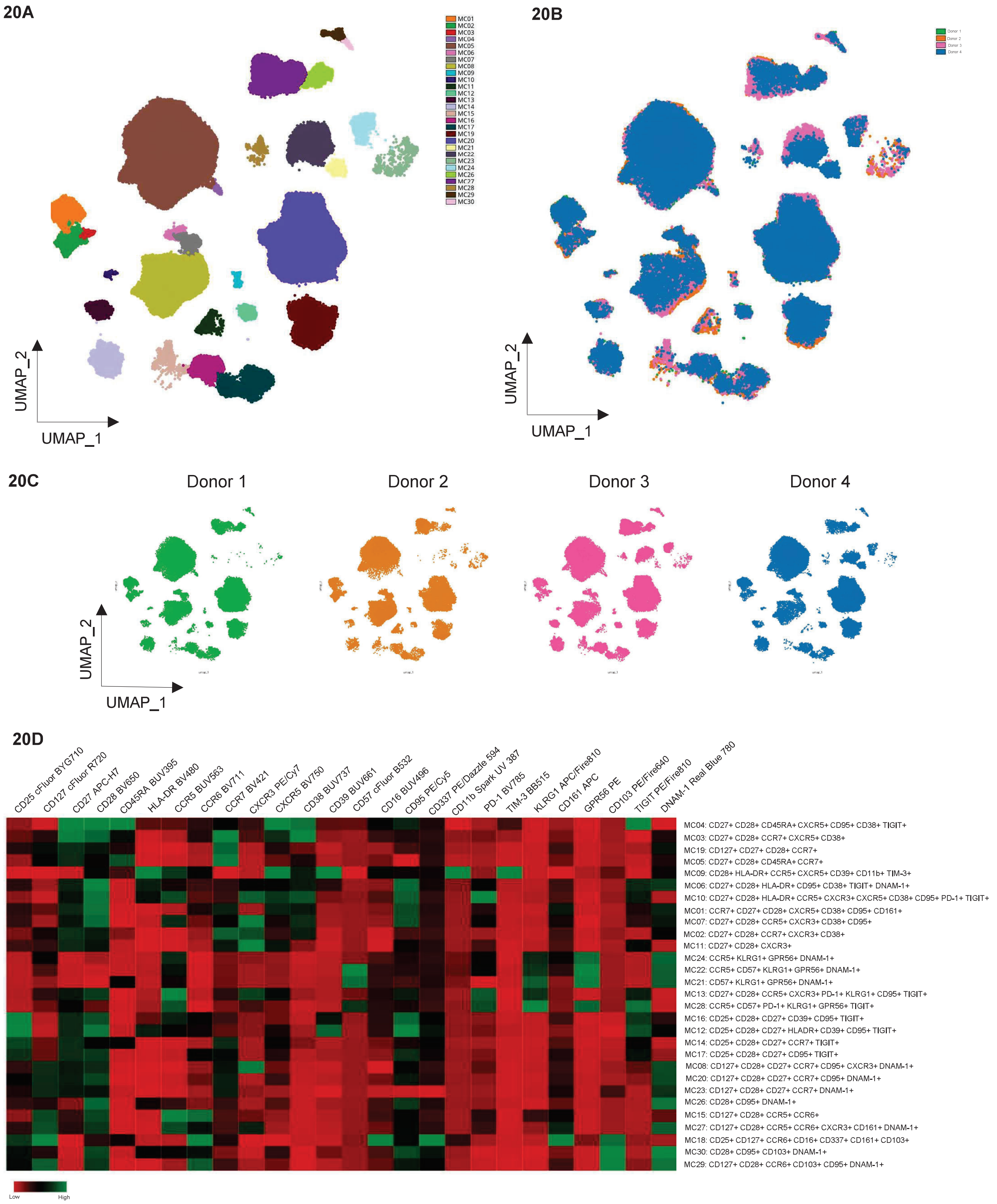

**Online Figure 20. Multidimensional Data Analysis of CD4+ T cells**

Data was analyzed with the OMIQ platform using a pipeline similar to the one used in OMIP-069. PeacoQC was used for data QC and FlowSOM for clustering. After the FlowSOM analysis, dimensionality was reduced, and clusters were displayed in a UMAP map. UMAP settings were as follows: all files used, Neighbors = 80, Minimum Distance = 0.7, Components = 2, Metric = Euclidean, Learning Rate = 1, Epochs = 250, Random Seed = 9346, Embedding Initialization = spectral. **(A)** Metaclusters are visualized on the UMAP parameters for ease of comparison between samples. Samples acquired on the same day from four donors were tested, **(B)** overlaid and **(C)** individual UMAPs are shown. **(D)** Following the UMAP analysis, a heatmap was generated with the resulting populations clustered hierarchically. Marker expression intensity is displayed on a scale from red (negative) to green (positive).

defined manually, with a more comprehensive description of their entire phenotype. We used UMAP plots for each marker in the panel to confirm cluster identification **(Online Figure 21A).** Marker expression was also verified visualizing each cluster in overlayed scatterplots with all markers in the panel included **(Online Figure 21B)**. When comparing clusters across donors, similarities and differences were identified. For example, differences in metacluster 22 abundance were identified across donors, with the highest frequency observed in donor 3 **(Online Figure 21C)**. CD4 T cell Metacluster 22 expresses CD45RA, HLA-DR, CD38, CD57, CD95, KLRG1, CD161, CCR5, GPR56, and DNAM-1, suggesting a memory terminal effector exhausted nature in this population.

A similar strategy was used to dissect the CD8 T cell compartment. **Online Figure 22** shows 30 metaclusters projected in a UMAP, and the overlay of 4 donors, for consistency in cluster definition and frequency **(A-C)**. The heatmap displays the variety of populations identified based on marker intensity **(D)**. To illustrate, three metaclusters expressing CD103, a marker of tissue residency, were identified. Metacluster 11, 15, and 20, characterized also by expression of memory markers, including(CD95, DNAM-1, CD28, and CD27), and the absence of CD45RA and CCR7, pointing towards an early effector profile. Each of these 3 clusters was defined as unique by their on/off expression of CD25, CD127, HLADR, CCR6, CXCR3, and CD337. CD8+ MAIT cells were identified in metaclusters 8 and 12, by coexpression of CD95, CD161, KLRG1, CD28, CCR5, CCR6, and CD337, but with key activation differentiators between them (CD127, CD39, CD38 and HLA-DR). The expression pattern for each marker is displayed in **Online Figure 22E.**

Finally, we used our advance analysis strategy, to evaluate the impact of applying minor manual spillover adjustments to unmixed data. We observed similar UMAP positioning, cluster identification and cluster frequency when comparing data with or without manual corrections. Overlaid scatterplots and correlation of cluster abundance are shown for both CD4 and CD8 T cells in **Online**

21A

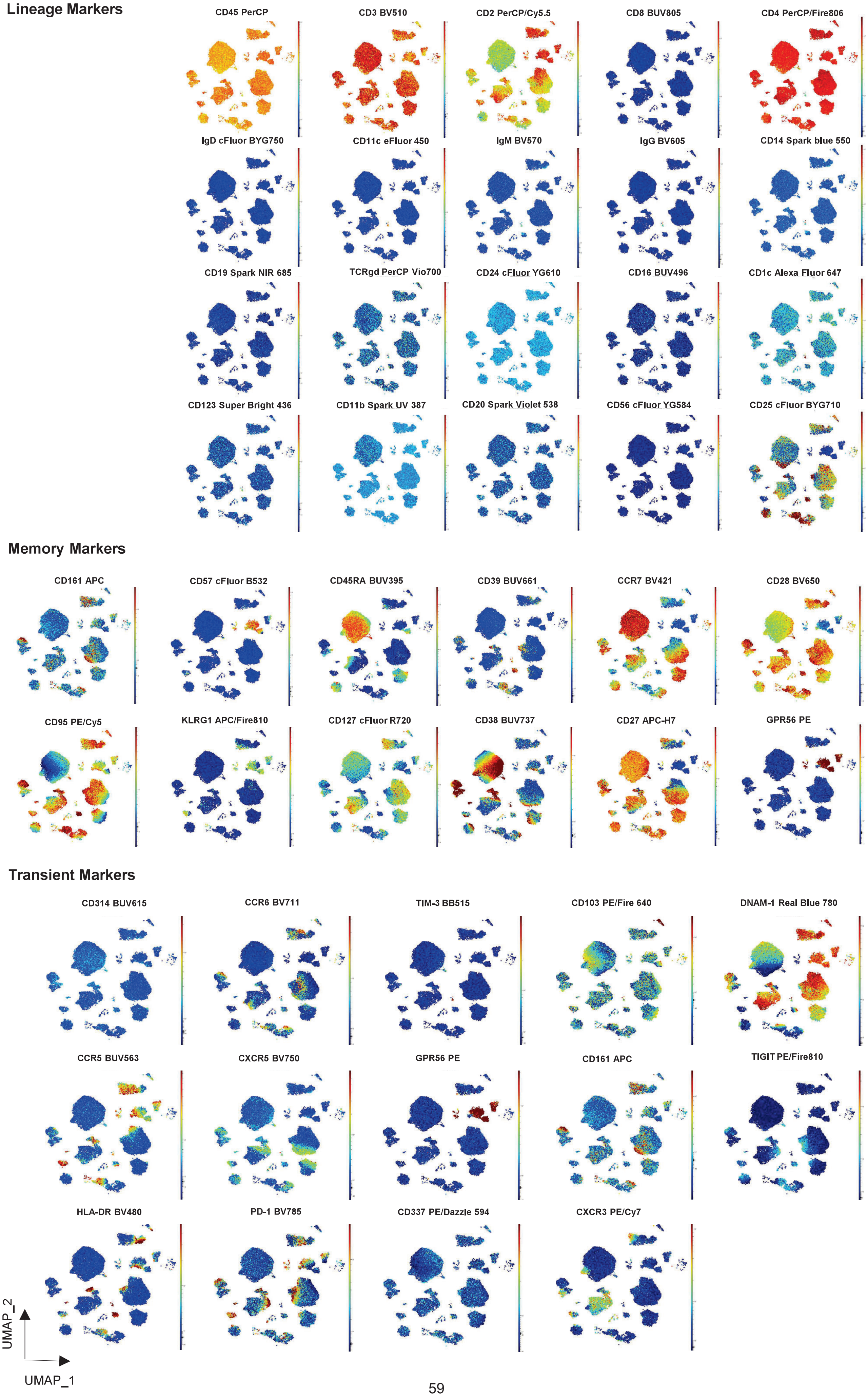

Lineage Markers
CD45 PerCP
CD3 BV510
CD2 PerCP/Cy5.5
CD8 BUV805
CD4 PerCP/Fire806
IgD cFluor BYG750
CD11c eFluor 450
IgM BV570
IgG BV605
CD14 Spark blue 550
CD19 Spark NIR 685
TCRgd PerCP Vio700
CD24 cFluor YG610
CD16 BUV496
CD1c Alexa Fluor 647
CD123 Super Bright 436
CD11b Spark UV 387
CD20 Spark Violet 538
CD56 cFluor YG584
CD25 cFluor BYG710
Memory Markers
CD161 APC
CD57 cFluor B532
CD45RA BUV395
CD39 BUV661
CCR7 BV421
CD28 BV650
CD95 PE/Cy5
KLRG1 APC/Fire810
CD127 cFluor R720
CD38 BUV737
CD27 APC-H7
GPR56 PE
Transient Markers
CD314 BUV615
CCR6 BV711
TIM-3 BB515
CD103 PE/Fire 640
DNAM-1 Real Blue 780
CCR5 BUV563
CXCR5 BV750
GPR56 PE
CD161 APC
TIGIT PE/Fire810
HLA-DR BV480
PD-1 BV785
CD337 PE/Dazzle 594
CXCR3 PE/Cy7
UMAP_2
UMAP_1

**21B**

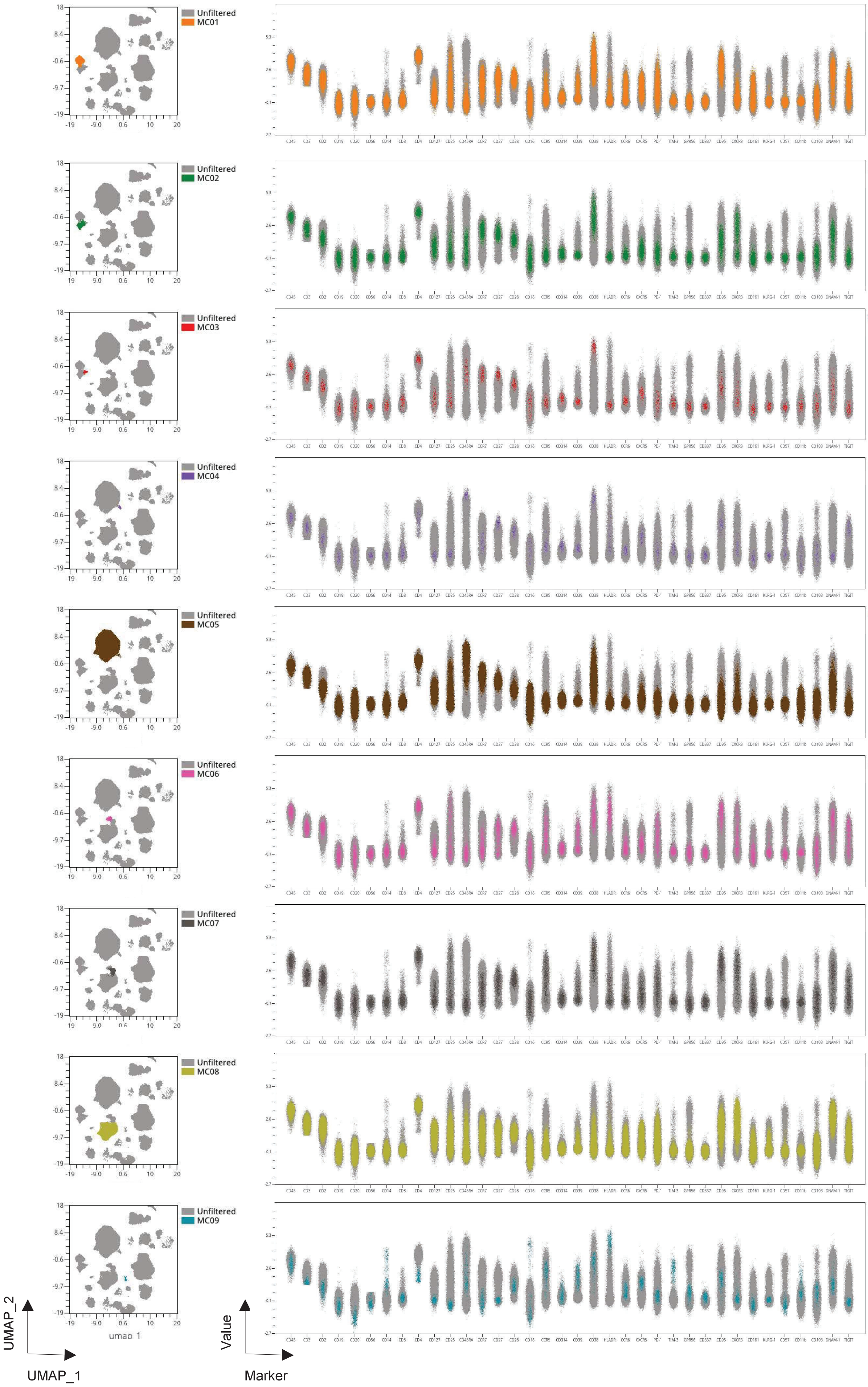

Unfiltered
MC01
Unfiltered
MC02
Unfiltered
MC03
Unfiltered
MC04
Unfiltered
MC05
Unfiltered
MC06
Unfiltered
MC07
Unfiltered
MC08
Unfiltered
MC09
UMAP_2
UMAP_1
Value
Marker

**21B**

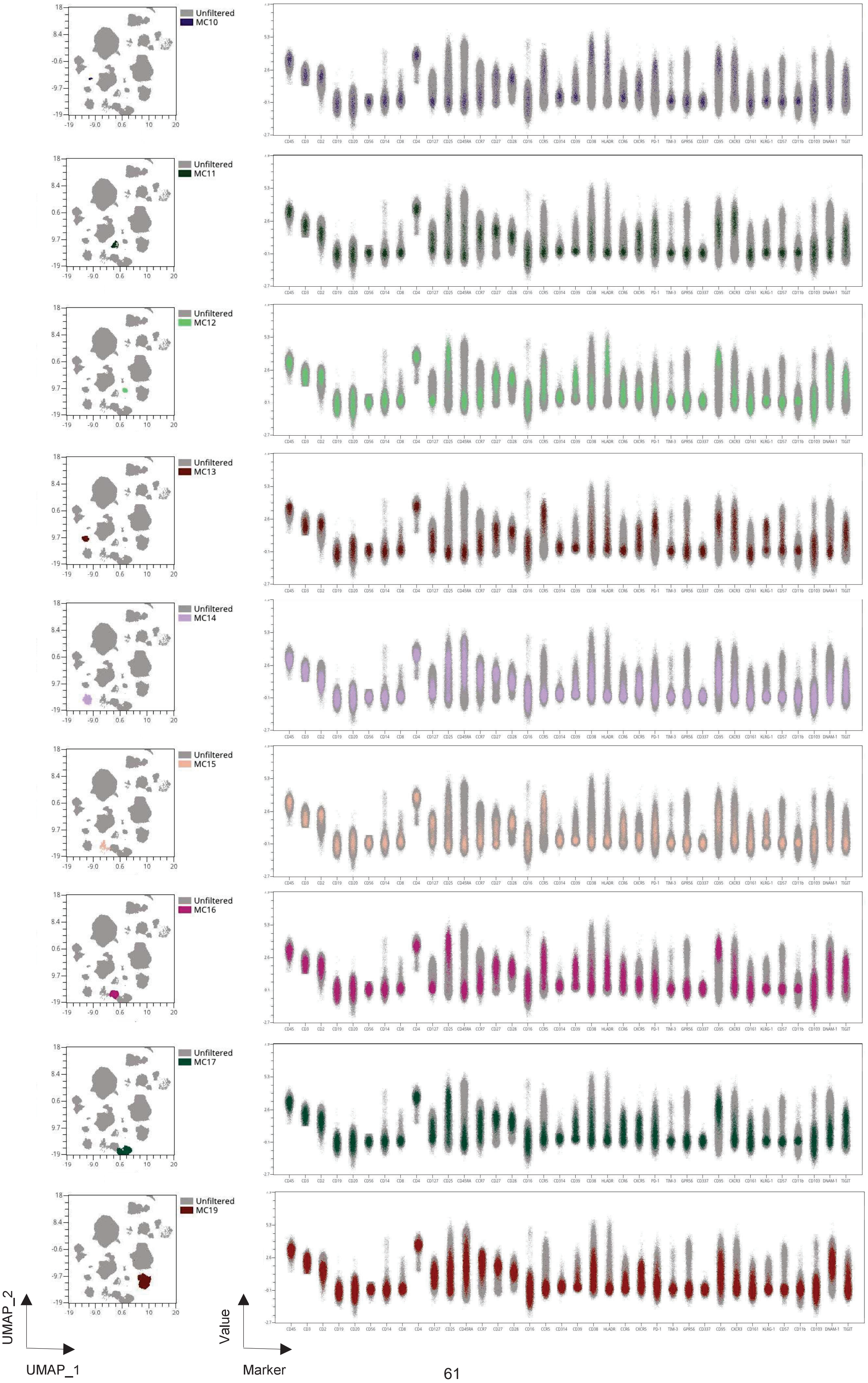
Unfiltered
MC10
MC11
MC12
MC13
MC14
MC15
MC16
MC17
MC19
UMAP_2
UMAP_1
Value
Marker
CD45 CD3 CD2 CD19 CD20 CD56 CD14 CD8 CD4 CD127 CD25 CD45RA CCR7 CD27 CD28 CD16 CCR5 CD314 CD39 CD38 HLADR CCR6 CXCR5 PD-1 TIM-3 GPR56 CD337 CD95 CXCR3 CD161 KLRG-1 CD57 CD11b CD103 DNAM-1 TIGIT

**21B**

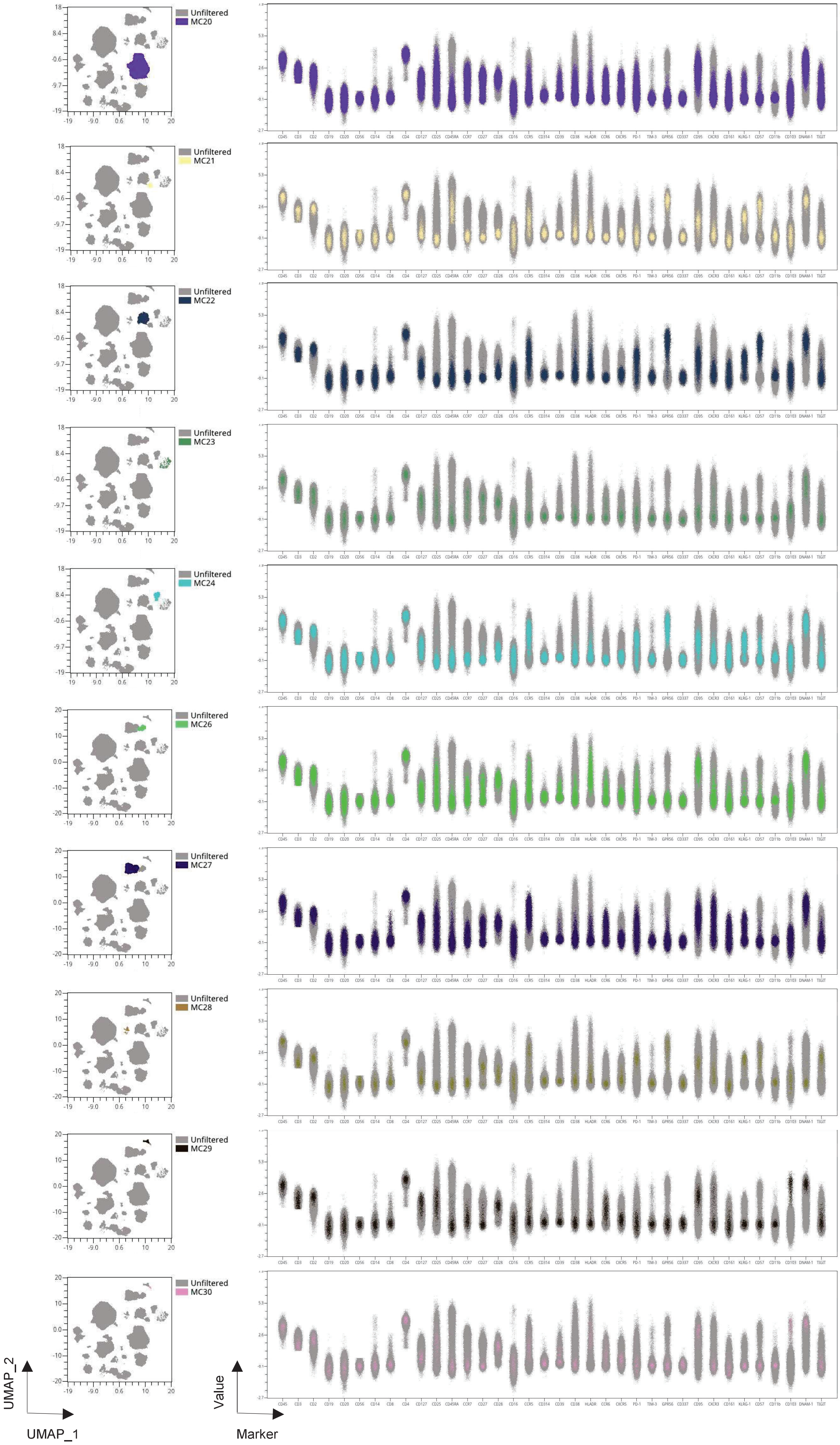
Unfiltered
MC20
Unfiltered
MC21
Unfiltered
MC22
Unfiltered
MC23
Unfiltered
MC24
Unfiltered
MC26
Unfiltered
MC27
Unfiltered
MC28
Unfiltered
MC29
Unfiltered
MC30
UMAP_2
UMAP_1
Value
Marker
CD45 CD3 CD2 CD19 CD20 CD56 CD14 CD8 CD4 CD127 CD25 CD45RA CCR7 CD27 CD28 CD16 CCR5 CD314 CD39 CD38 HLADR CCR6 CXCR5 PD-1 TIM-3 GPR56 CD337 CD95 CXCR3 CD161 KLRG-1 CD57 CD11b CD103 DNAM-1 TIGIT

**21C**

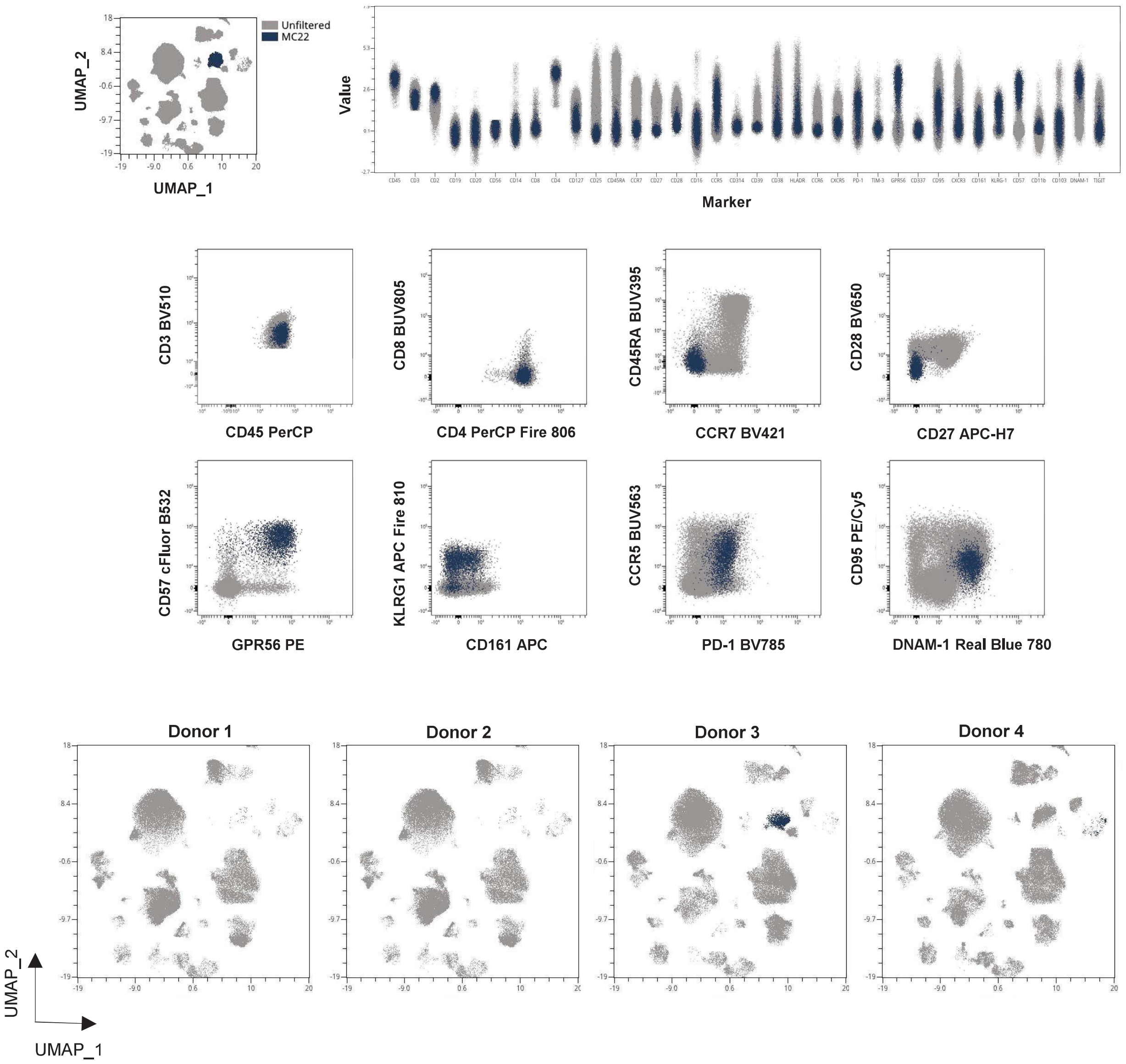

**Online Figure 21. CD4 T cell Cluster Verification**

Metacluster identity was verified in multiple ways. **(A)** First, UMAP colour-continuous scatterplots show the intensity of expression of each marker in the panel, including the ones not used for clustering to verify phenotype. Markers are divided into lineage, memory, and transient groups, to facilitate data exploration. Marker positivity or negativity for lineage markers was confirmed. Marker expression intensity is indicated by the scale bar to the right of each plot, where red is high, and blue is low. Second, cluster phenotype was verified by using overlay scatter plots which display the arc sin transformation value for each critical marker used to define metaclusters. **(B)** For each cluster, UMAP (left) and overlay scatterplots (right) display 2 layers, in grey unfiltered events and each cluster in the corresponding color shown in the clustering map. Last, metaclusters were manually explored through back gating to verify phenotype. **(C)** Metacluster 22 is shown in detail. A UMAP and an overlay scatterplot for all markers show 2 layers, unfiltered events in grey and cluster events in dark blue. The sequential backgating for metacluster 22 is shown, to confirm expression of CD3, CD45, CD4, CD57, GPR56, KLRG1, PD-1, CCR5, DNAM-1 and CD95 dim, and the exhausted memory nature of this subpopulation.

22A

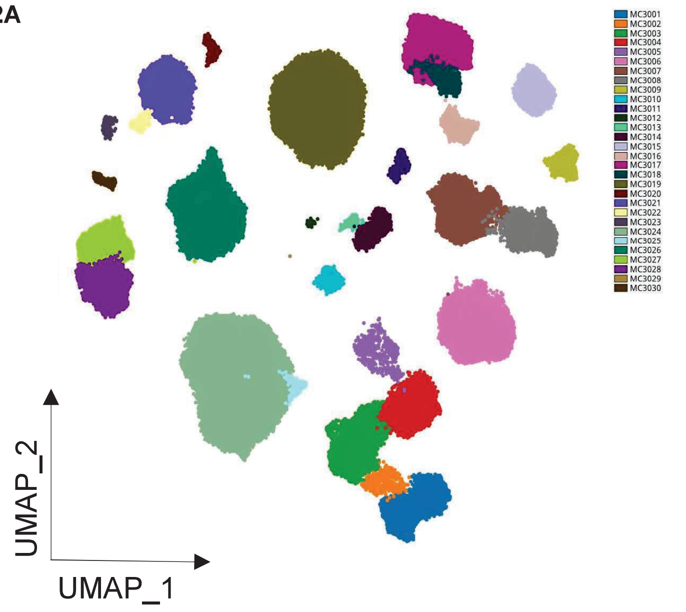

22B

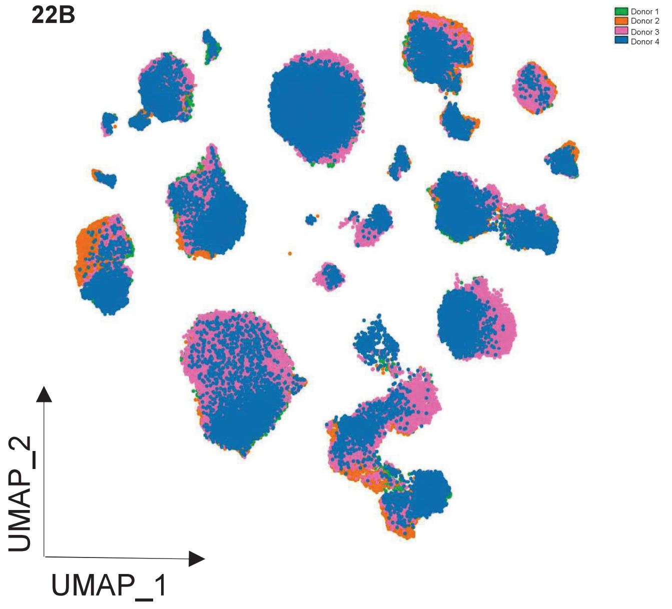

22C

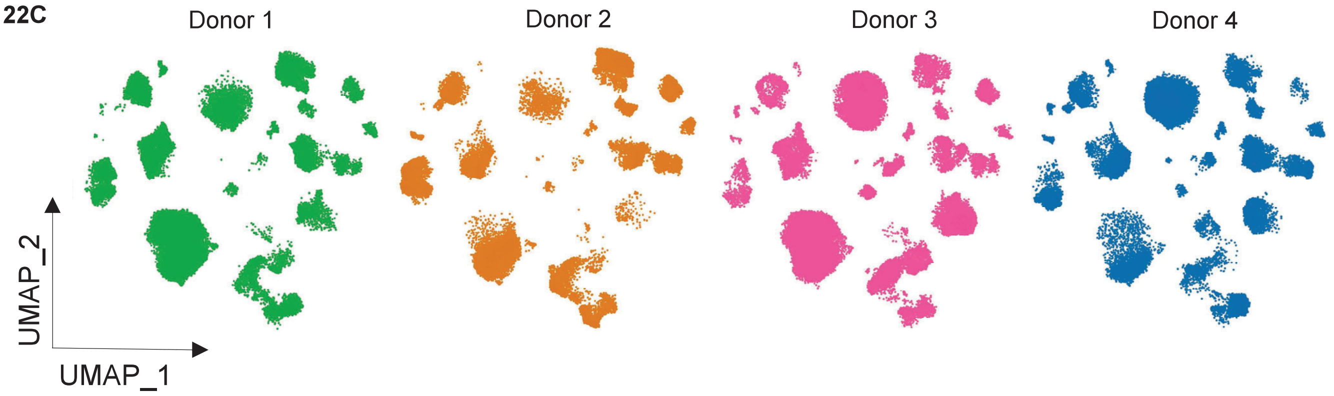

22D

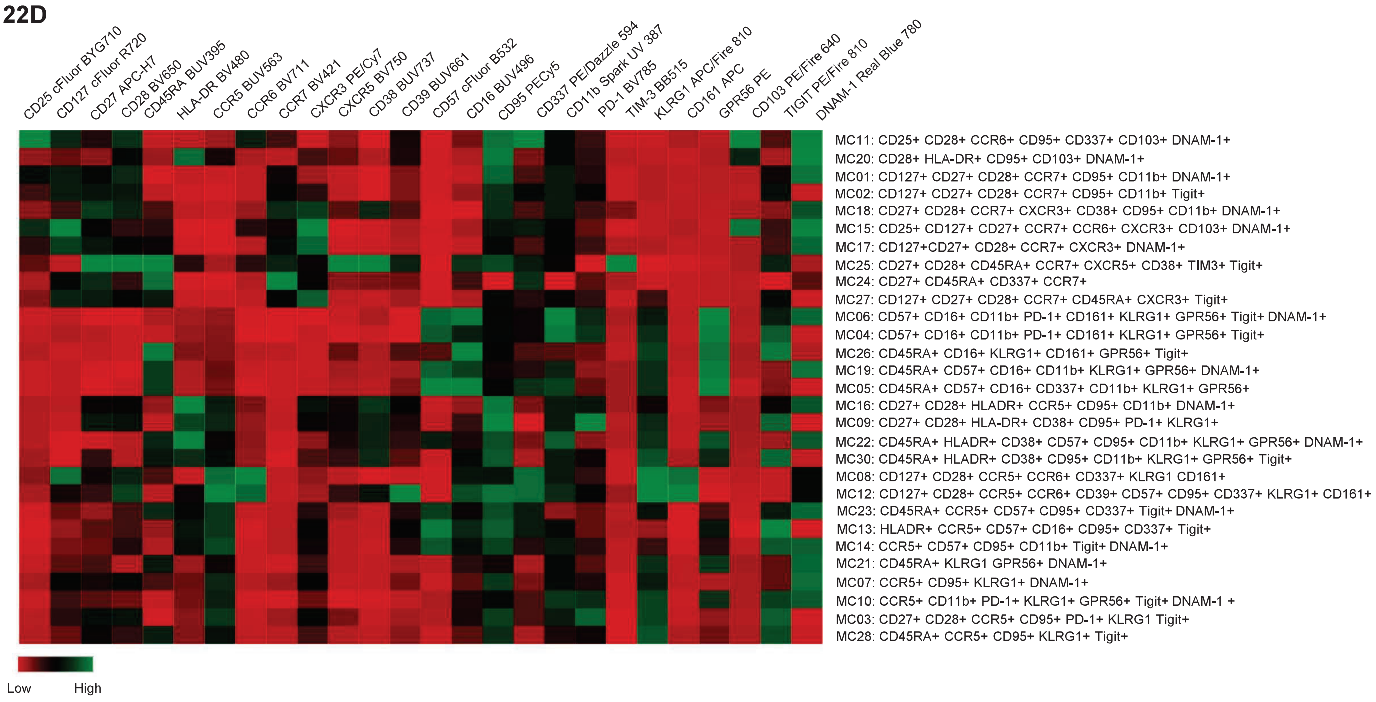

22E

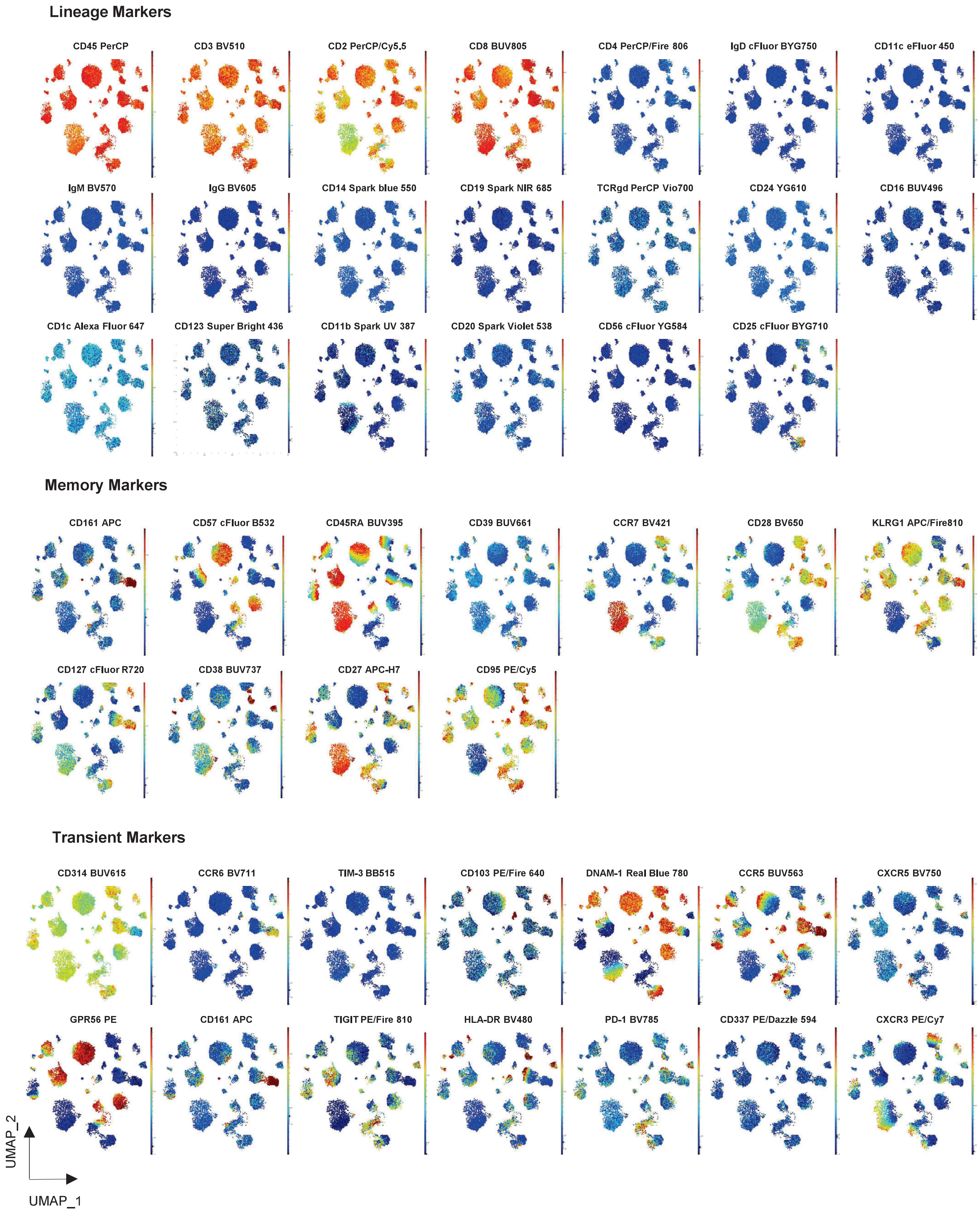

**Online Figure 22. Multidimensional Data Analysis of CD8+ T cells**

Data was analyzed with the OMIQ platform a pipeline similar to the one described for CD4 T cells. **(A)** CD8 metaclusters are displayed in a UMAP map. **(B)** Four donors overlaid, **(C)** donor individual UMAPs, **(D)** clustered hierarchical heatmap, and **(E)** UMAP color-continuous scatterplots for the intensity of expression of each marker in the panel.

**Figures 23A.** A similar comparison was made with all live single CD45+ events in the same sample. We observed no perturbation on cluster definition when manual adjustments were needed (**Online Figure 23**).

#### c. Impact of paraformaldehyde fixation on assay performance

Since it is not always possible to acquire samples right after finishing the staining procedure, fixation with formaldehyde (PFA) is often used. We investigated the impact of fixation with 1% PFA in PBS for 20 minutes followed by a wash with stain buffer and final resuspension in stain buffer (PBS, BSA, NaN3 based), This is our preferred method as it uses a relatively low concentration of PFA and limits the exposure of the fluorochromes, as it is well known that PFA can have a negative impact on the stability of tandem fluorochromes. We compared panel performance without fixation, and immediately, 2, 18 and 24 hours after. Of note, throughout the development of the panel we did not use fixation. Three performance criteria were evaluated: 1) our ability to gate/identify the populations of interest, 2) changes in the frequencies of those populations, and 3) any decrease in intensity (MFI) for any marker. Immediately after fixation, the vast majority of the populations remained well resolved and were easily identifiable without moving the gate positioning established with the fresh sample, as shown for T cells and B cell subsets in **Online Figure 24A**. However, the CD24 cFluor YG610 intensity and the frequency of CD123+ pDCs changed within the first two hours, as presented in **Online Figure 24A**. An unsupervised analysis comparison of fresh vs immediately fixed samples, focused on CD4 and CD8 T cells, is shown in **Online Figure 24B.**

#### d. Assessment of assay performance across analyzers and sorter with the same optical configuration

A key component of any panel is that it can provide reproducible results across cytometers with similar optical configurations, this OMIP is the first to demonstrate this ability. We evaluated the

23A

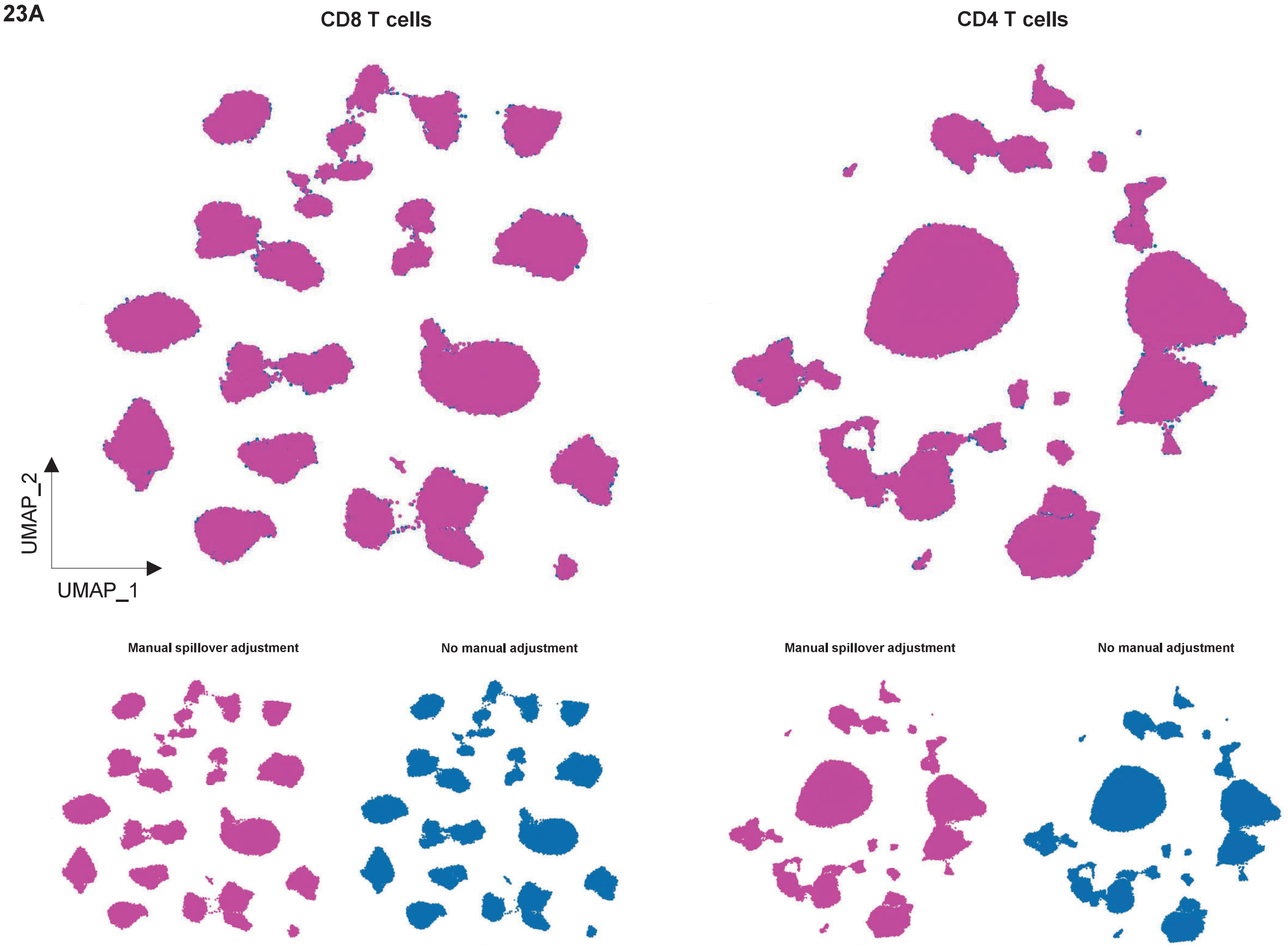

CD8 T cells
CD4 T cells
UMAP_2
UMAP_1
Manual spillover adjustment
No manual adjustment
Manual spillover adjustment
No manual adjustment

23B

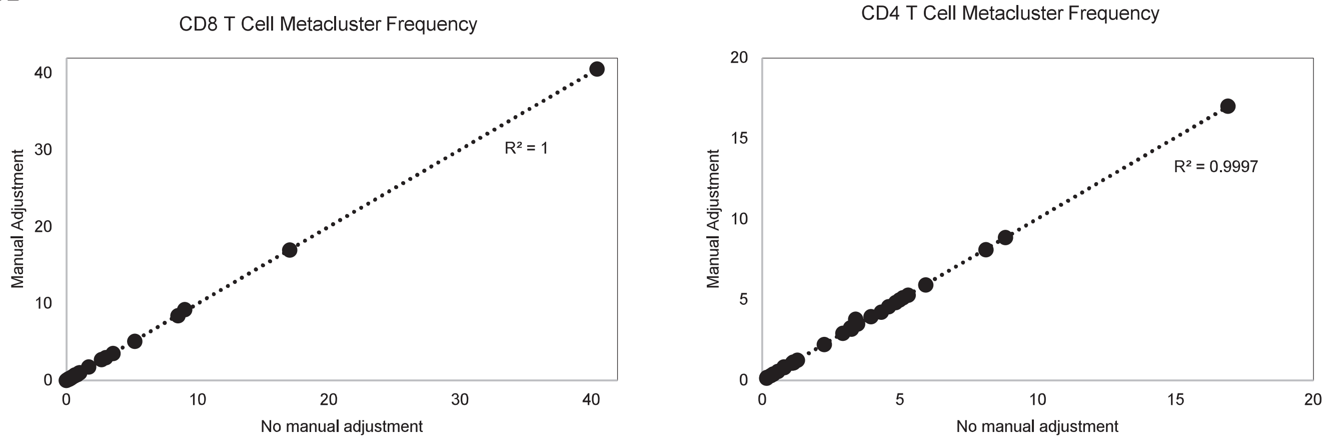

CD8 T Cell Metacluster Frequency
Manual Adjustment
No manual adjustment
0
10
20
30
40
R² = 1
CD4 T Cell Metacluster Frequency
Manual Adjustment
No manual adjustment
0
5
10
15
20
R² = 0.9997

**23C**

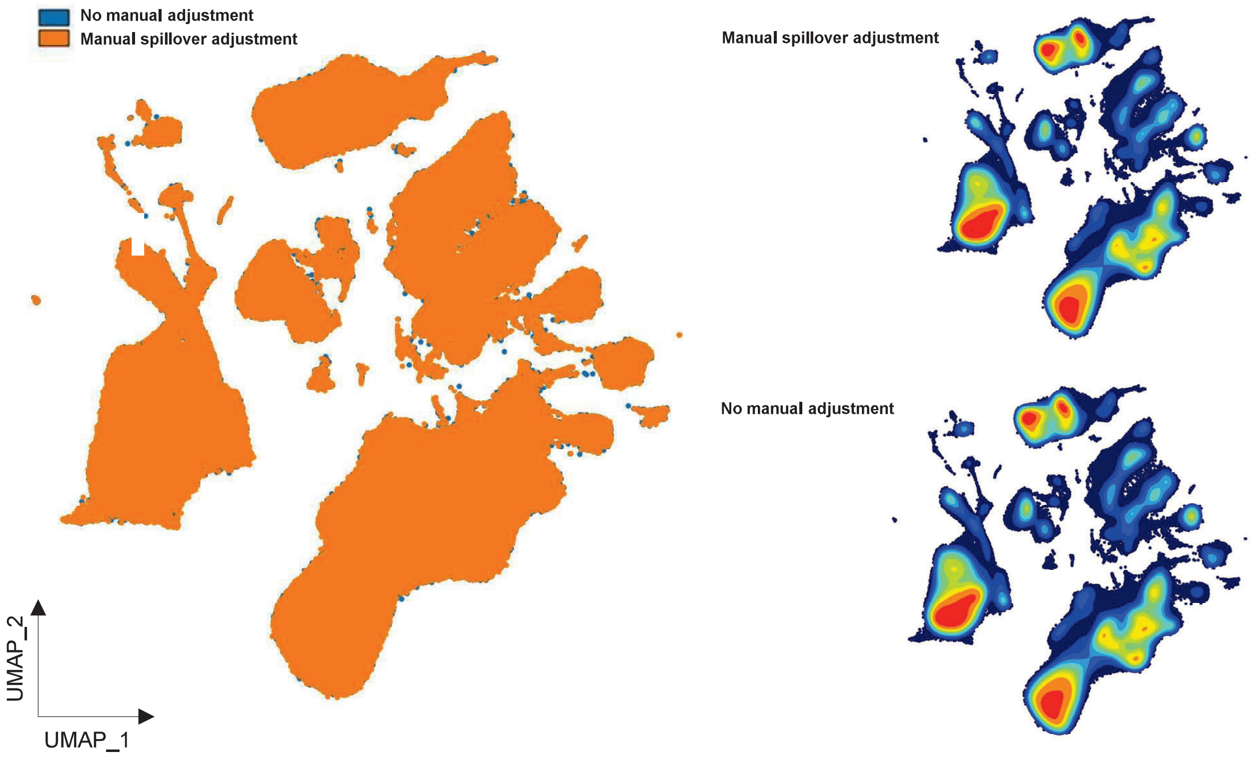

**23D**

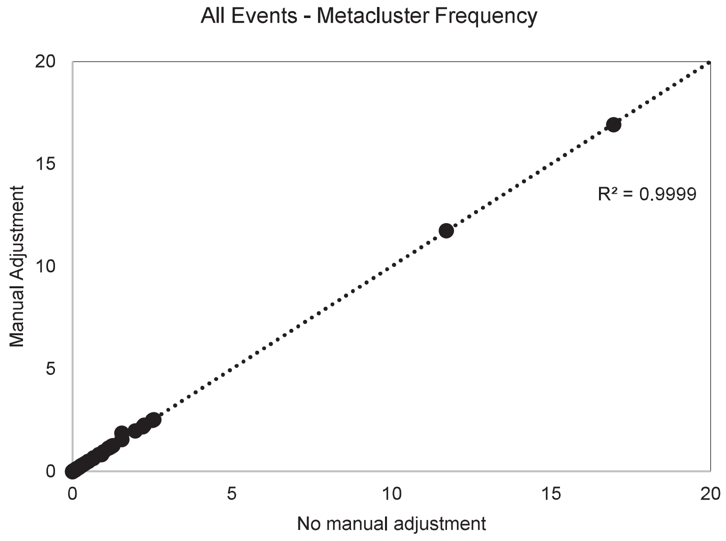

**Online Figure 23. Comparison of Data Analysis With or Without Manual Spillover Adjustments**

Data was analyzed with the OMIQ platform a pipeline similar to the one described before. **(A)** FlowSOM and UMAP analysis is shown for one donor, comparing fcs files without manual spillover adjustments upon unmixing against files with manual corrections. Manual adjustments were included only for aesthetic purposes. Overlaid and individual UMAPs for CD4 and CD8 T cells show the consistency in cluster identification and position. **(B)** Cluster frequency was also compared, with high correlation of abundances between unmixing strategies. Pearson's rho was calculated using Microsoft excel. Clustering and dimensionality reduction were performed in all events (450,000 events per sample) to verify that manual adjustments do not impact metacluster definition. Clustering was performed using all markers, except for CD45 and viability dye. FlowSOM metaclustering was set to 50 and 100 K value. **(C)** The consistency of overlaid UMAPs for a file with manual spillover adjustment vs the same file without correction upon unmixing is presented. Individual UMAPs are also shown for each condition. **(D)** Frequency of the 50 metaclusters was compared, with high correlation of abundances between the 2 unmixing strategies is shown. Pearson's rho was calculated using Microsoft excel.

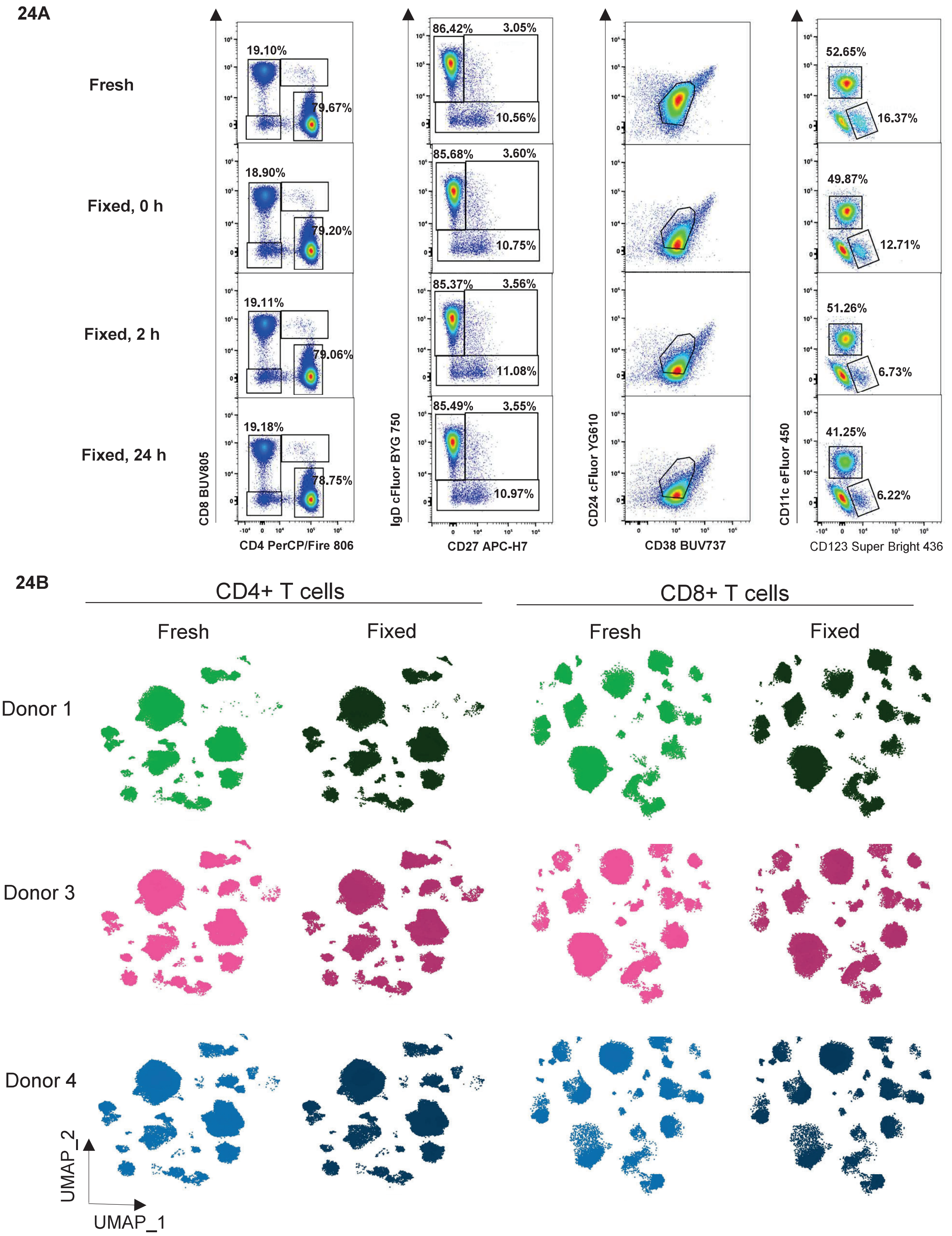

**Online Figure 24. Impact of Paraformaldehyde Fixation on Panel Performance**

Panel performance with or without fixation was compared for 3 donors. Upon staining, cells were fixed with 1% formaldehyde (PFA) in PBS for 30 minutes, followed by a wash and final resuspension in wash buffer. Samples were acquired immediately (fresh), and 2, 18 and 24 hours after fixation. **(A)** CD4-CD8, CD27-IgD, CD24-CD38, and CD123-CD11c bivariate plots are shown for the 4 conditions tested (fresh, fixed 0h, fixed 2h, and fixed 24h). **(B)** A comparison of unsupervised analysis of fresh vs immediately fixed sample is shown for CD4 and CD8 T cells, using FlowSOM and UMAP. Cluster identity and frequency was preserved upon fixation. Individual UMAPs are shown for 3 donors.

consistency of assay performance across three 5-laser Auroras and one 5-laser Aurora CS (Cell Sorter). All instruments were set up at "CytekAssaySetting 2x UV1". The same optimal reference controls (beads and cells per the recommendations listed on Online Table 2) were acquired on all instruments as well as the same multicolor tube, using the same stopping criteria, and the same analysis template. No gate adjustments were needed as all populations were equally well resolved across instruments and were in remarkably similar positions (data not shown). **Online Figure 25** shows the results of the unsupervised analysis for the CD4 and CD8 subsets across instruments, The same clusters with the same location and abundance were obtained. In summary, the data from all 4 instruments is highly comparable and this result gives us reassurance that the panel will perform optimally in other laboratories with access to 5-laser Cytek Aurora and Aurora CS instruments. This panel was only evaluated on Cytek instruments. If using other spectral cytometers, the resolution for some markers may be different depending on optical performance and instrument setup.

#### e. Sorting of T cell memory subsets and functional characterization by intracellular cytokine staining

Finally, we wanted to assess the utility of the panel for cell sorting experiments. Since the panel performed equally well on the Aurora CS as in the analyzers, i.e., all populations were clearly defined as show in section d, making it very straightforward to define the gates needed to identify the populations to be sorted. To explore the functional diversity of effector CD4 T cells, we decided to sort specific T cell subsets that were identified by expression of, or lack of expression of, a high number of markers included in this panel. Those specific subsets are found in circulation in low frequencies in healthy donors, highlighting one of the key utilities of the panel. The following 6 subsets were sorted from the $CD45^+CD3^+CD19^-CD56^-CD14^-CD4^+CD8^-$ population **(Online Figure 26A)**:

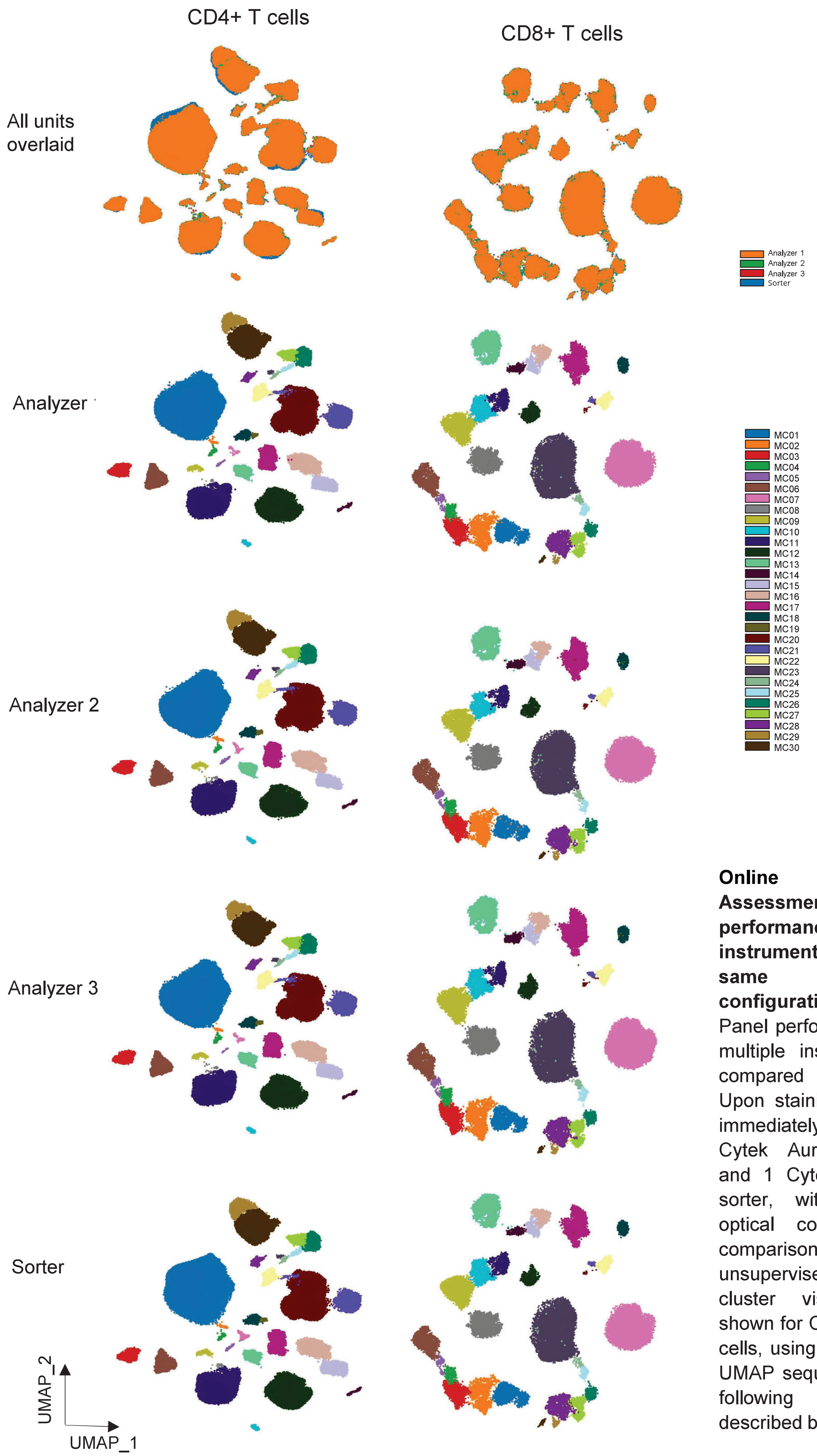

**Online Figure 25. Assessment of assay performance across instruments with the same optical configuration**

Panel performance across multiple instruments was compared for 1 donor. Upon staining, cells were immediately acquired on 3 Cytek Aurora analyzers and 1 Cytek Aurora cell sorter, with the same optical configuration. A comparison of unsupervised analysis and cluster visualization is shown for CD4 and CD8 T cells, using FlowSOM and UMAP sequential analysis following the pipeline described before.

A. TSCM: $CCR7^+CD45RA^+CD27^+CD28^+CD127^+CD95^+$

B. TEM early effector: $CCR7^-CD45RA^-CD27^+CD28^+KLRG1^+TIGIT^-CCR6^-DNAM\text{-}1^+$

C. TEM early effector: $CCR7^-CD45RA^-CD27^+CD28^+KLRG1^+TIGIT^-CCR6^+DNAM\text{-}1^+$

D. TEM early effector: $CCR7^-CD45RA^-CD27^+CD28^+KLRG1^-TIGIT^-CCR6^+$DNAM-1+

E. TEM early effector: $CCR7^-CD45RA^-CD27^+CD28^+KLRG1^-TIGIT^-CCR6^-$DNAM-1+

F. TEMRA: $CCR7\ ^-CD45RA^+CD27^+CD28^+CD127^+CD95^+$

Cells were sorted at an average of 4500 events/second using a 100 µm nozzle. From $5x10^7$ total cells, we recovered $1.1x10^4$, $7x10^3$, $2x10^3$, $3x10^4$, $7x10^3$, and $5x10^4$ cells of populations A-F, respectively. After sorting, the cells were spun down and resuspended in 200 µL of complete RPMI and rested overnight at 37ºC, 5% CO2. The next day the cells were stimulated with PMA ionomycin to analyze their ability to secrete cytokines upon mitogenic stimulation.

The strategy to design the panel for intracellular staining was straightforward. Since the phenotype of the sorted cells was clearly defined, several markers from the 45-color panel were not expressed in these cells and therefore the fluorochromes assigned to those markers could be paired with anti-cytokine antibodies. Specifically, Spark NIR 685, cFluor 584, eFluor 450, and BV605 were available since the sorted cells did not express CD19, CD56, CD11c, or IgG, respectively. Moreover, in our laboratory, we had previously optimized intracellular cytokines assays and we had already identified good performing clones for the 4 cytokines we were interested in analyzing. The following antibodies were selected: IFNγ Spark NIR 685 (clone 4S. B3, Biolegend, Cat. 502551), IL-2 Real Yellow 586 (equivalent to cFluor 584, similarity index 1, clone MQ1-17H12, BD Biosciences Cat. 568468) TNF-αPacific Blue (equivalent to eFluor 450, similarity index 1, clone Mab11, BioLegend, Cat. 502920), and IL-17 BV605 (clone BL168, BioLegend, Cat. 512325). Because of the small number of cells available from the sorted populations, we set apart five million cells that were not sorted as the experimental control for the cytokine assay. Both stimulation and -non-stimulation conditions were evaluated. **Online Figure 26B** shows a summary of the results.

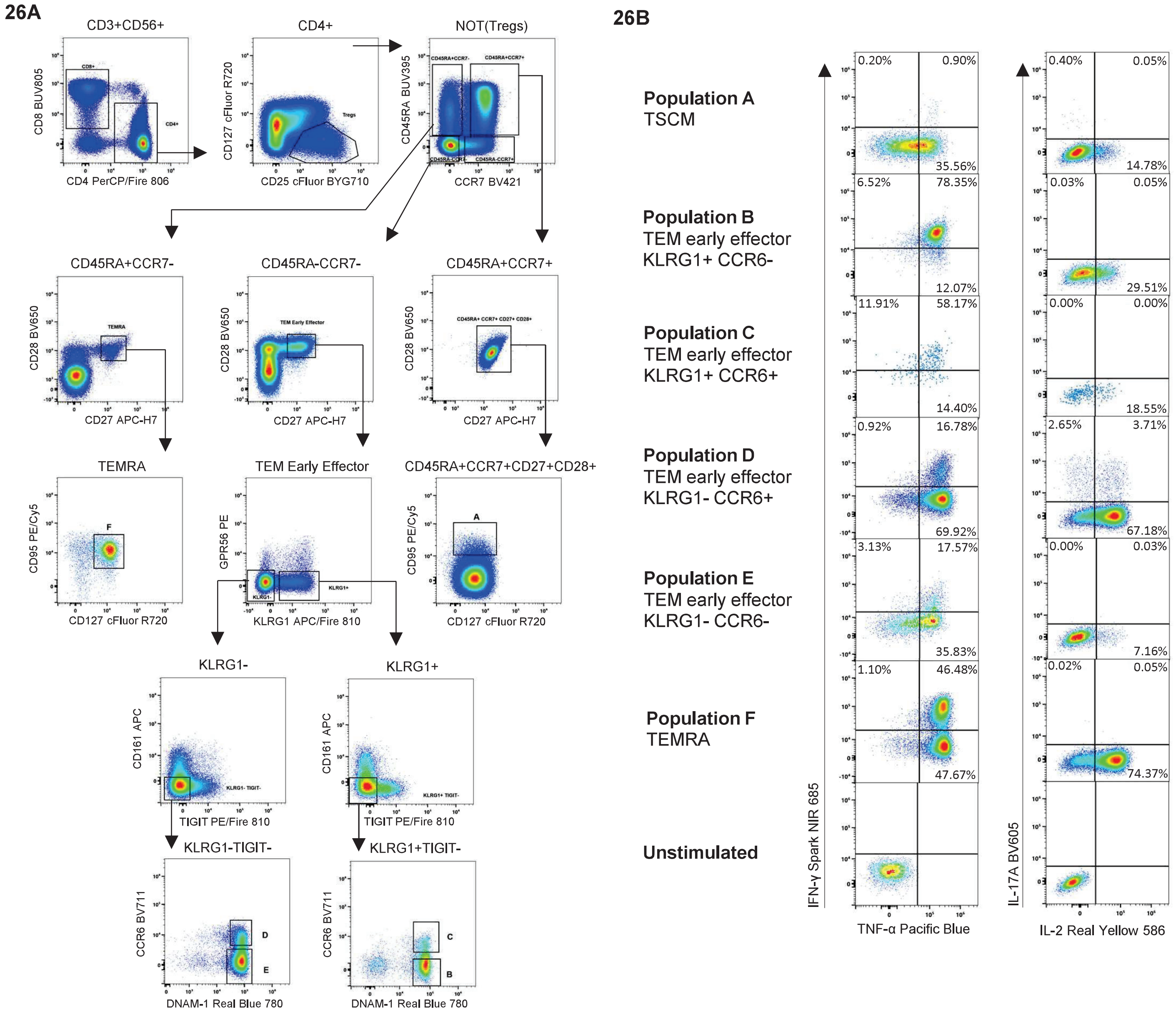

**Online Figure 26. Sorting and intracellular cytokine staining**

**(A)** CD4 T cells characterized with the 45-color panel, were further gated to distinguish 6 populations:

A. TSCM: CCR7$^+$CD45RA$^+$CD27$^+$CD28$^+$CD127$^+$CD95$^+$

B. TEM early effector: CCR7$^-$CD45RA$^-$CD27$^+$CD28$^+$KLRG1$^+$TIGIT$^-$CCR6$^-$DNAM-1$^+$

C. TEM early effector: CCR7$^-$CD45RA$^-$CD27$^+$CD28$^+$KLRG1$^+$TIGIT$^-$CCR6$^+$DNAM-1$^+$

D. TEM early effector: CCR7$^-$CD45RA$^-$CD27$^+$CD28$^+$KLRG1$^-$TIGIT$^-$CCR6$^+$DNAM-1$^+$

E. TEM early effector: CCR7$^-$CD45RA$^-$CD27$^+$CD28$^+$KLRG1$^-$TIGIT$^-$CCR6$^-$DNAM-1$^+$

F. TEMRA: CCR7$^-$CD45RA$^+$CD27$^+$CD28$^+$CD127$^+$CD95$^+$

Cells were sorted at an average of 4500 events/second using a 100 µm nozzle. After sorting, cells were spun down and resuspended in 200 µL of complete RPMI and rested overnight at 37ºC, 5% CO2. Cells were stimulated with 50ng/mL PMA and 1µg/mL ionomycin for 4h to induce cytokine production. 1 hour after treatment, BFA and monensin were added at 5 µg/mL, each, to prevent cytokine secretion. Four intracellular cytokines were evaluated by flow cytometry: IFNγ Spark NIR 685 (clone 4S. B3, Biolegend, Cat. 502551), IL-2 Real Yellow 586 (clone MQ1-17H12, BD Biosciences Cat. 568468) TNF-α Pacific Blue (clone Mab11, BioLegend, Cat. 502920), and IL-17 BV605 (clone BL168, BioLegend, Cat. 512325). Unstimulated cells were used as a control. **(B)** IFNγ vs TNF-α and IL-2 vs IL-17A plots are shown for each population post-sort, post-stimulation. The sorted subpopulations had very distinct functional cytokine profiles.

Briefly, all sorted populations were activated, and each one had a distinct functional pattern. All subsets secreted elevated levels of TNF $\alpha$ (from 70 to 95% of the cells) except for population A (36%) in which most of the cells were negative or expressed intermediate levels of this cytokine, as expected for TSCM cells. Populations A and E secreted extremely low levels of IFN$\gamma$ (1.10% and 20.70% respectively). In contrast to cells in subset B, in which 84.87% secreted IFN$\gamma$. Interestingly, subsets D and F had the highest frequency of cells producing IL-2 (70.89% and 74.41% respectively), in agreement with the lack of expression of the exhaustion KLRG1 marker in both populations. The other 4 subsets had low levels of IL-2. Finally, subset D, early effector TEM cells positive for CCR6, was the only subset with cells able to produce IL-17 (6.36%), with 58% coexpression of IL-2. Unstimulated cells are shown as a gating control. These results indicate that the distinct phenotypes identified within the CD4 T cell effector compartment have different functionality, and hence would play a different role in immune responses. The fine delineation and sorting of these functionally diverse subsets is only possible with a highly parametric panel as the one described here. Characterization of these and other subsets in patient samples would highly likely shed light on their biological significance and role in disease progression and response to treatment.

### 6.Sample preparation and staining procedure

<u>Materials and Reagents</u>

12x75 mm Falcon® 5 mL Polypropylene (PP) Test Tubes: Corning, catalog #352063

Ficoll® Paque Plus: Sigma, 17-1440-02

Paraformaldehyde solution 4% in PBS, Santa Cruz Biotechnology, Cat. sc-281692

RPMI-1640 Medium: Sigma, catalog # R8758-500 mL

Fetal Bovine Serum (FBS): Gibco, catalog #16000044

Penicillin-Streptomycin: Gibco, catalog #15140122

PBS, pH 7.2: Gibco, catalog #20012-027

BD Horizon™ Brilliant Stain Buffer Plus : BD Biosciences, catalog #568264

True-Stain Monocyte Blocker™: BioLegend, catalog #426101

UltraComp eBeads™ Plus Compensation Beads: Thermo Fisher, catalog #01-3333-41

Wash/Stain Buffer: BD Pharmingen™ Stain Buffer (BSA), BD Biosciences catalog #554657

Antibodies specified in Table 1

Prepared buffers: complete RPMI: add 50 mL of FBS and 5 mL of Penicillin-Streptomycin into 500 mL of RPMI 1640 (with L-glutamine and sodium bicarbonate)

Thawing of PBMCs

1. Pre-warm complete RPMI at 37˚C for a least 30 min
2. Thaw as quickly as possible
    a. Thaw cryo-vial ($25x10^6$ cells) in 37˚C water bath, until only a small piece of ice remains.
    b. Transfer contents of cryo-vial to 50 mL conical tube
    c. Add 1 mL of warm complete RPMI to cryo-vial. Leave aside until step f.
    d. Drop-by-drop add 5 mL of warm complete RPMI to the cells in the 50 mL tube. While adding, gently mix the 50 mL tube (with a pipette in one hand and in the other the 50 mL tube, add the complete RPMI while you gently swirl the tube).
    e. After the first 5 mL of complete RPMI has been added, add the next 5 mL a little bit faster (a few drops at a time).
    f. After 10 mL have been added, pour the contents of the cryo-vial into the 50 mL tube.

g. Add additional volume of complete RPMI to 20 mL.

3. Spin at 400 x *g* for 8 minutes
4. Decant supernatant carefully without disturbing the pellet.
5. Gently resuspend pellet in 2 mL of warm complete RPMI.
6. Add additional volume of complete RPMI to 20 mL.
7. Repeat steps 3-5.
8. Resuspend in 7.5 mL of complete RPMI or to an approximate concentration of $3.5x10^6$/ml).
9. Leave in incubator until ready to proceed (rest is not necessary, but once staining has begun, samples will need to be processed without delay).

Preparation of freshly isolated PBMCs

Note: Handling of human blood components should be done in accordance with regional and institutional biosafety policies and/or requirements.

Note: Fresh blood samples should be kept at 18ºC to 20ºC throughout the protocol.

1. Collect 15 mL of whole blood into heparin tubes.
2. Using a 50 mL conical tube, dilute the blood in PBS at 1:1 ratio.
3. Invert the tube gently 5 times to mix.
4. Bring the Ficoll® Paque Plus media bottle to room temperature.
5. Invert the Ficoll® several times to mix thoroughly.
6. Add 3 mL of Ficoll® to the bottom of 15 mL conical tubes, avoiding getting the solution on the tube wall.
7. Tilt the 15 mL tube with Ficoll® media and slowly layer 7 mL of the diluted blood sample.

8. Close the cap and centrifuge at 400 x *g* for 40 minutes, room temperature (RT) with the break turned off.

9. Remove the top layer of plasma using a serological pipet while avoiding disturbing the mononuclear cell layer.

10. Carefully take the layer of mononuclear cells and transfer to a separate 15 mL conical tube.

11. Complete the volume to 14 mL with Wash/ Stain buffer, mix well by inversion of the tube.

12. Centrifuge at 400 x *g* for 8 minutes at RT.

13. Decant

14. Carefully resuspend the cell pellet in 1 mL of Wash/Stain buffer.

15. Repeat steps 11-13.

16. Resuspend in 5 mL of Wash/Stain buffer and count cells.

17. Adjust the concentration to 3 x $10^6$ cells/mL.

<u>Viability dye Preparation</u>

1. Completely thaw DMSO. Add 50 µL DMSO to the lyophilized LIVE/DEAD Blue Dead Cell Stain Kit, following the manufacturer's recommendation.
2. Vortex to mix thoroughly.
3. Aliquot 7 µL and store at -20°C until use.
4. Thaw a fresh vial of aliquot of the stock solution at room temperature, protected from light, before each use.
5. Dilute the stock solution at 1:20 in PBS (5 µL stock solution in 195 µL 1X PBS).
6. Use the working solution at 10 µL per test.

Antibody Cocktail Preparation

1. Prepare multicolor antibody cocktail in a 1.7 mL microcentrifuge tube by first adding, per sample, 10 μL of Brilliant Stain Buffer Plus to the tube, then all the antibodies at pre-determined titers EXCEPT CCR5 BUV563, CD123 Super Bright 426, IgG BV605, TCRγδ PerCP-Vio 700, CXCR3 PE/Cy7, CCR7 BV421, CXCR5 BV750, PD-1 BV785, and CD11c eFluor 450.

Note: The nine reagents listed above need to be added sequentially for optimal results.

Note: Prepare one extra test for the multicolor cocktail to account for any reagent loss.

Viability dye Reference Control Staining

1. Label a 12x75 mm Falcon® 5 mL PP Test Tube for Viability dye Reference Control.
2. Add 100 μL of resuspended cells to the tube (~$3x10^5$ cells).
3. Add 1X PBS to complete the final volume to 3 mL.

Note: Viability dye is an amine reactive dye and will be bound by any proteins in the buffer. Therefore, it is important to make sure the wash buffer is free of protein.

4. Centrifuge at 400 x *g*, 5 minutes at RT.
5. Decant supernatant and blot on paper towel.
6. Vortex thoroughly.
7. Repeat steps 3-6.
8. Add 10 μL of working solution LIVE/DEAD Fixable Blue Dead Cell Stain Kit to the cell pellet.
9. Vortex thoroughly.
10. Incubate for 15 minutes at RT, protected from light.
11. Add 3 mL of Stain Buffer.
12. Centrifuge at 400 x *g*, 5 minutes at RT.

13. Decant supernatant and blot on paper towel.
14. Vortex thoroughly.
15. Repeat steps 11-14.
16. Resuspend in 300 µL Stain Buffer.
17. Acquire at medium flow rate.

Reference Controls Staining

1. Label 12x75 mm Falcon® 5 mL PP Test Tubes for Single Stain (SS) controls.
2. Add 100 µL of resuspended cells to each tube (~$3x10^5$ cells).

Note: We recommend using all controls as cells, if possible. However, beads can be used as reference controls for some antibody reagents that showed minimal to no difference in unmixing accuracy compared to cell reference controls. Refer to **Online Table 4** for the reference control type recommendation. Please note that the recommendation is made using UltraComp eBeads™ Plus Compensation Beads and that beads from other manufacturers may perform differently.

3. Add pre-determined optimal amount of antibody to the appropriate tube.

Note: Add 10 µL of Brilliant Stain Buffer Plus to BUV805 CD8 SS Tube only.

4. Incubate for 20 minutes at RT, protected from light.
5. Add 3 mL of Stain Buffer.
6. Centrifuge at 400 x *g*, 5 minutes at RT.
7. Decant supernatant and blot on paper towel.
8. Vortex thoroughly.
9. Repeat steps 5-8.
10. Resuspend in 300 µL Stain Buffer.
11. Acquire at medium flow rate.

Multicolor Staining

1. Label a 12x75 mm Falcon® 5 mL PP Test Tubes for multicolor sample.
2. Add 1 mL of resuspended cells to the tube (~$3x10^6$ cells).
3. Add 1X PBS to complete the final volume to 3 mL.

Note: Viability dye is an amine reactive dye and will be bound by any proteins in the buffer. Therefore, it is important to make sure the wash buffer is free of protein.

4. Centrifuge at 400 x *g*, 5 minutes at RT.
5. Decant supernatant and blot on paper towel.
6. Vortex thoroughly.
7. Repeat steps 3-6.
8. Add 10 µL of working solution LIVE/DEAD Fixable Blue Dead Cell Stain Kit to the cell pellet.
9. Vortex thoroughly.
10. Incubate for 15 minutes at RT, protected from light.
11. Add 3 mL of Stain Buffer.
12. Repeat steps 3-6.

12\. Add 5 µL of Tue-Stain Monocyte Blocker™ and 10 µl of Brilliant Stain Buffer Plus to all multicolor sample tubes. Vortex well.

13. Add CCR5 BUV563 antibody, vortex.
14. Incubate for 5 minutes at RT, protected from light.
15. Add CD123 Super Bright 436 and IgG BV605 antibodies, vortex.
16. Incubate for 10 minutes at RT, protected from light.
17. Add TCRγδ PerCP-Vio 700 antibody, vortex.
18. Incubate for 5 minutes at RT, protected from light.

19. Add CXCR3 PE/Cy7, CCR7 BV421, CXCR5 BV750, PD-1 BV785, and CD11c eFluor 450, vortex.
20. Incubate for 10 minutes at RT, protected from light.
21. Add the remaining antibodies from the multicolor cocktail mix, vortex.
22. Incubate for 20 minutes at RT, protected from light.
23. Add 3 mL of Stain Buffer.
24. Centrifuge at 400 x *g*, 5 min. at RT.
25. Decant supernatant and blot on paper towel.
25. Resuspend in 400 µL Stain Buffer.
26. Acquire at medium flow rate.

<u>Paraformaldehyde (PFA) Fixation</u>

Note: If the samples need to be stored at 4ºC for more than 2 hours prior to data collection, follow these steps to fix the samples in 1% PFA and acquire within 24 hours of fixation.

Note: For accurate unmixing/compensation, SS controls need to be treated the same .as the multicolor stained sample, including the fixation step. This is true whether beads or cells are used as SS controls. Fluorochrome emission properties can be slightly altered depending on their microenvironment.

1. Dilute 4% PFA in 1X PBS to make 1% paraformaldehyde solution.
2. After the final washing step of each staining process, add 500 µL of 1% PFA in PBS to the cell pellet, vortex.
3. Incubate for 20 minutes at RT, protected from light.
4. Add 3 mL of Stain Buffer.
5. Centrifuge at 400 x *g,* 5 minutes at RT.
6. Decant supernatant and blot on paper towel.

7. Vortex thoroughly.
8. Resuspend in appropriate volume in Stain Buffer.
9. Store at 4ºC, protected from light and acquire within 24 hours post-fixation.

**Bulk staining for sorting**

1. Adjust the cell concentration to 20 to 25 x $10^6$ cells/mL in PBS and aliquot 2 mL of cell suspension per multicolor tube.
2. Add 50 µL of diluted Live Dead Blue solution to each multicolor tube.
3. Vortex thoroughly.
4. Incubate for 15 minutes at RT, protected from light.
5. Add 2 mL of Wash/Stain buffer.
6. Centrifuge at 400 x *g* for 5 minutes at RT.
7. Decant.
8. Add 4 mL of Wash/Stain Buffer.
9. Repeat steps 6-7.
10. Resuspend in 2 mL of Wash/Stain Buffer
11. Add 4x the titer of each antibody, following the sequential staining steps.
12. Complete the volume to 4 mL of Wash/Stain buffer.
13. Centrifuge at 400 x *g* for 5 minutes at RT.
14. Decant
15. Repeat steps 12-14
16. Resuspend in 2 mL of desired buffer for sorting.
